# Leopard Cats Occupied Human Settlements in China for 3,500 years before the Arrival of Domestic cats in 600-900 CE around the Tang Dynasty

**DOI:** 10.1101/2025.01.31.635809

**Authors:** Yu Han, Songmei Hu, Ke Liu, Xiao Xu, Sean Doherty, Alexandra Elizabeth Jamieson, Aurelie Manin, Sofia Granja Martins, Miaomiao Yang, Chong Yu, Juan Wang, Zhuang Wu, Canping Chen, Sicheng Han, Daowei Lu, Lanhui Peng, Xianzhu Wu, Ziyi Li, Wenquan Fan, Quanfa Cai, Zongliang Cui, Jing Yuan, Zihan Li, Yang Liu, Zhipeng Li, Zhendong Liu, Qian Ma, Jing Shao, Zhouyong Sun, Fulai Xing, Wuzhan Yang, Shugang Yang, Lianjian Yue, Pengcheng Zhang, Weilin Wang, Huanyuan Zhang, Yan Zhuang, Xin Sun, Yan Pan, Xiaohong Wu, Laurent A. F. Frantz, He Yu, Joel M. Alves, Greger Larson, Shu-Jin Luo

## Abstract

The earliest cats in human settlements in China were not domestic cats (*Felis catus*), but native leopard cats (*Prionailurus bengalensis*). To trace when and how domestic cats arrived in East Asia, we analyzed 22 feline bones from 14 sites across China spanning 5,000 years. Nuclear and mitochondrial genomes revealed that leopard cats began occupying anthropogenic scenes around 5,400 years ago and last appeared in 150 CE. Following several centuries’ gap of archeological feline remains, the first known domestic cat (706–883 CE) in China was identified in Shaanxi during the Tang Dynasty. Genomic analysis suggested a white or partially white coat and a link to a contemporaneous domestic cat from Kazakhstan, indicating a likely dispersal route via the Silk Road. The two felids once independently occupied ancient anthropogenic environments in China but followed divergent paths and reached different destinations in human-animal interactions.

## Main Text

The African wildcat (*Felis lybica*) is the ancestor of all modern domestic cats (*F. catus*) (*1–3*). Though the timing and circumstances of cat domestication are uncertain, zooarchaeological evidence demonstrates that the northward dispersal of domestic cats into Europe took place by 3,000 BP (*4*,*5*). Domestic cats were also transported eastwards across Eurasia into the Far East and China, but the route and timing of this dispersal remain unresolved.

A previous study of 5,400-year-old felid remains from the site of Quanhucun in western China concluded that cats had been predating rodents within human settlements, thus implying that domestic cats were present in China by the late Neolithic (*6*). Subsequent morphological (*7*) and ancient mtDNA (*8*) analyses of the same specimens revealed that the remains did not belong to the domestic cat but were instead leopard cats (*Prionailurus bengalensis*), a small wild felid native to continental South, Southeast, and East Asia (*9, 10*). These results thus require a reconsideration of when domestic cats arrived in China, how long leopard cats persisted in archaeological contexts, and whether the two felids coexisted.

To address these questions, we retrieved whole genome sequences from 22 ancient small felid bones from 14 Chinese archaeological sites dated from c.3,500 BCE to 1,800 CE (Fig. 1A and table S1 and table S2). This sample set represents the vast majority of feline excavations from China thus far. We directly radiocarbon dated six individuals and generated seven nuclear (2.4× - 6.0×) and 22 mitochondrial genomes (12× - 304×) through direct sequencing or targeted sequencing with hybridization capture (Fig. 1). We then analyzed these data in the context of global cat genetic variation, including 63 nuclear and 108 mitochondrial genomes (table S1) previously published from ancient and modern specimens.

**Figure 1.**
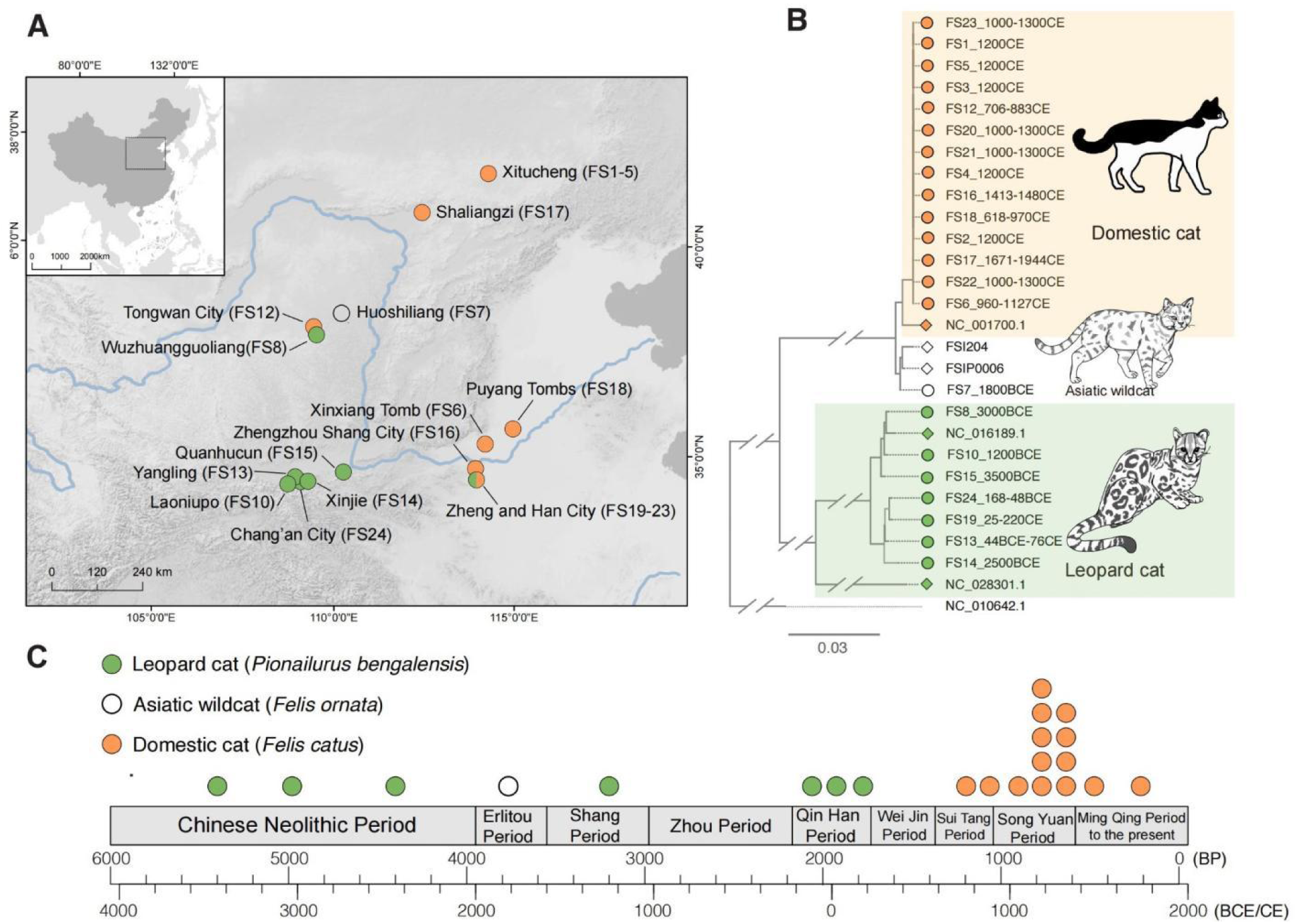
Sampling location, species identification, and age of small feline specimens from ancient China. **(A)** Archaeological sites of small felid specimen sampling in China. The colors of the dots correspond to the species of small felid found at the sites. The IDs of small felids excavated at each site are displayed in parentheses after the site names. The Zheng and Han City is the only site where both leopard cats and domestic cats were found. **(B)** A maximum likelihood (ML) phylogenetic tree constructed based on mitogenomes of ancient feline samples from China. All 22 cat samples in archaeological contexts are included in this analysis in addition to mitogenomes from *F. catus* (NC_001700.1), *F. ornata* (FSI204, FSIP0006), *P. bengalensis* (NC_016189.1, NC_028301.1), and the tiger (NC_010642.1) as an outgroup. **(C)** The temporal distribution of small felids from archeological sites in this study, with the horizontal time scale showing the historical periods of China. The dots represent specimens excavated from the corresponding period and are color-coded according to their species identification.

### Timing of the arrival of domestic cats in China

Among the 22 felid bones from China, 14 individuals spanning >1,000 years from c.730 CE to 1,800 CE were identified as domestic cats (*F. catus*) (Fig. 1 and table S2). These 14 individual cats were identified in seven archeological sites across multiple dynasties, including the Tang (618–907 CE), Song (960–1,279 CE), Yuan (1,279–1,368 CE), Ming (1,368–1,644 CE), and the Qing Dynasty (1,644–1,911 CE). The most ancient domestic cat in our dataset (FS12), identified using both nuclear and mitochondrial genomes, was excavated from Tongwan City (*11*) in Shaanxi, western China, and directly radiocarbon dated to c.730 CE (706–883 cal CE 95%) during the mid-Tang Dynasty (Fig. 1 and table S2). This result supports a relatively recent arrival of domestic cats in eastern Eurasia and undermines the commonly held assumption that domestic cats were present in China as early as the Han Dynasty (206 BCE–220 CE) (*12*).

All 14 Chinese archaeological cats cluster within mtDNA clade IV-B in the phylogenetic tree (Fig. 2 and fig. S3) (*1,2*). Though cats that possess clade IV-B haplotypes are rare in both ancient and modern individuals from western Eurasia (*5*,*13*), this clade includes a contemporaneous domestic cat from Dhzankent of Kazakhstan dated to 775–940 cal CE (*14*) (table S1), a contemporary African wildcat (FSI48) from the Levant, and 30 out of 38 (78.9%) modern domestic cats from China (*3*) (table S1).

**Figure 2.**
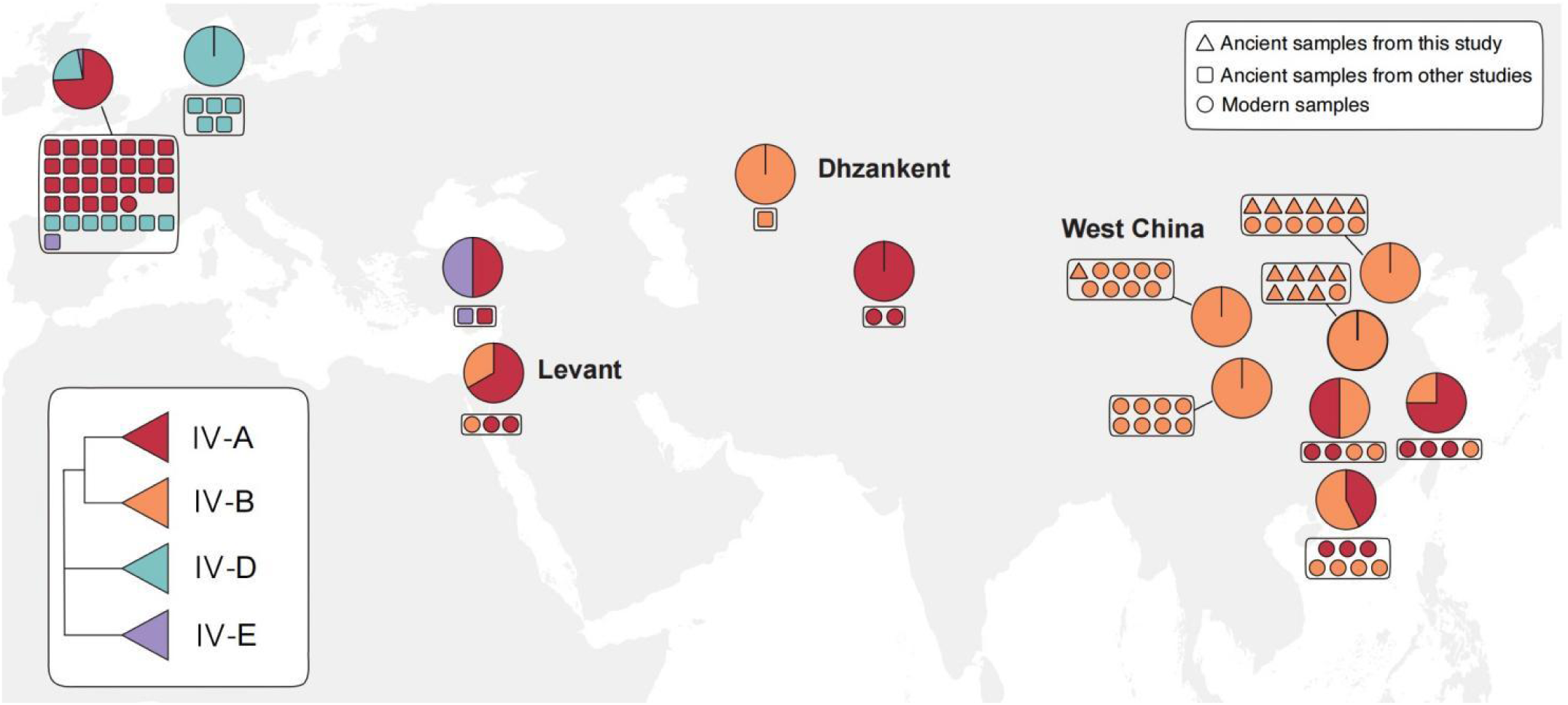
The geographical distribution of the mtDNA haplogroups of Eurasian domestic cats. A maximum likelihood (ML) phylogenetic tree is reconstructed based on mitogenomes from 14 ancient domestic cats in China and other cats from Eurasia. A simplified diagram of the phylogenetic tree is shown in the lower left corner of the figure. Each small icon corresponds to one specimen, and icons within the same box represent samples from the same location. The color of the icon corresponds to the mtDNA clade it belongs to, as shown in the simplified tree in the bottom left corner. The mtDNA clade coding system follows the previous description (*1*). Red triangles represent newly generated ancient samples in this study, black triangles represent published ancient samples, and circles represent published modern samples, as indicated in the upper right corner.

A principal component analysis (PCA) of nuclear genomes highlights three major clusters that correspond to domestic cat and/or African wildcats from western Eurasia, China, Central Asia, and the Levant (Fig. 3A). The PC1 axis separates western Eurasian samples from the rest of the dataset, and the PC2 axis distinguishes all major groups, including Chinese/Central Asian domestic cats, western Eurasian domestic cats, and Levantine African wildcats (Fig. 3A). All ancient and modern domestic cats from China form a cluster that is closely associated with an ancient cat from Dhzankent, Kazakhstan (*14*), which is thus far the earliest domestic cat on the Silk Road (Fig. 3A).

**Figure 3.**
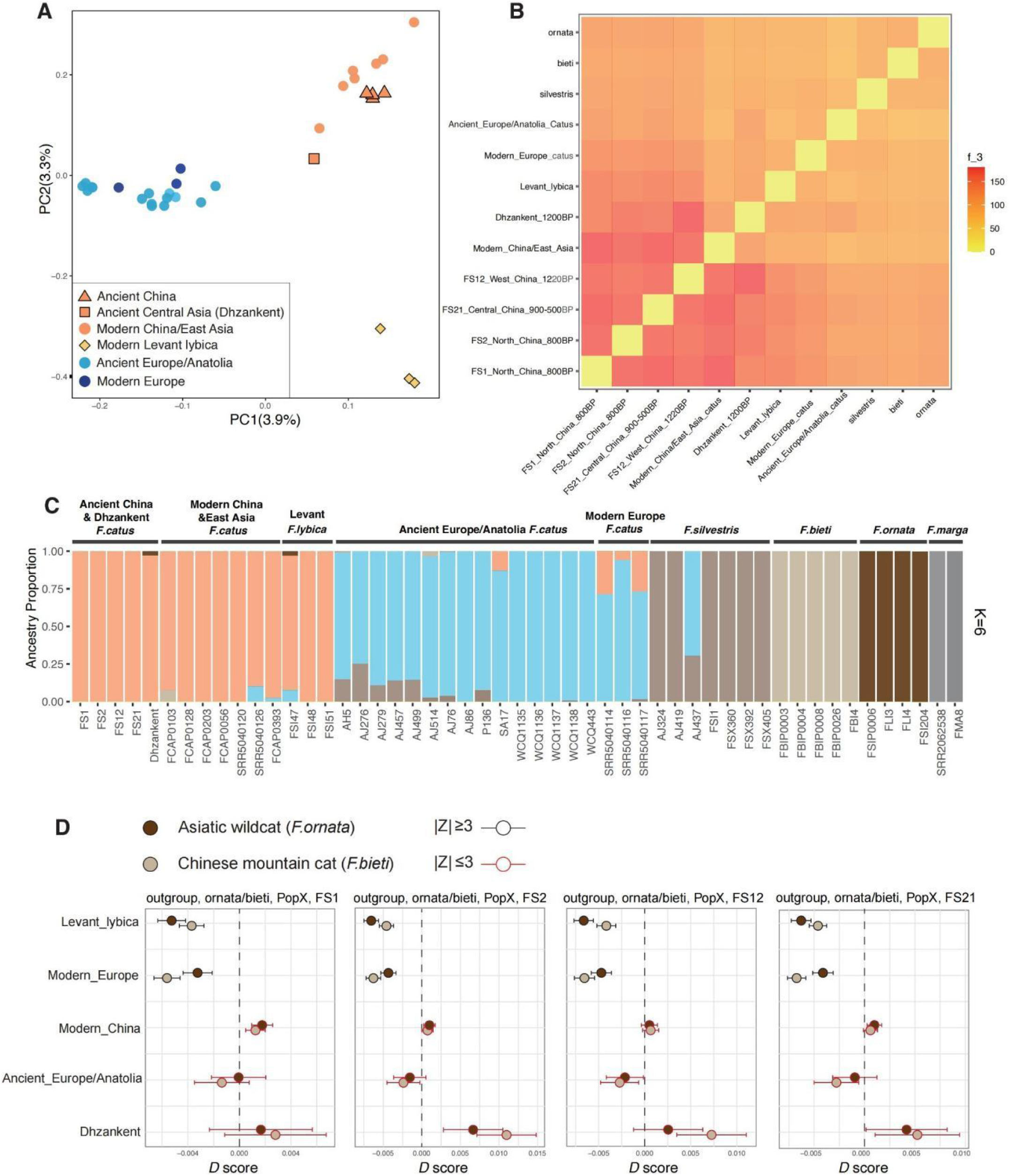
Nuclear genome analysis of ancient domestic cats in China. Four ancient Chinese domestic cats (FS1, FS2, FS12, FS21) with whole-genome sequencing data are included in the analysis. **(A)** The result of a Principal Component Analysis (PCA). Data from modern samples are used to construct an initial PCA followed by a subsequent projection of ancient samples. **(B)** The result of outgroup *f_3_-*statistics statistics. The shared drifts of different pairs of individuals from ancient domestic cats and other populations are tested. The sand cat (*F. margarita*) is used as an outgroup. The color gradient, ranging from yellow to red, signifies the magnitude of *f_3_* values. **(C)** Genetic clustering of domestic cats and wildcats inferred by ADMIXTURE (K=6). Each individual is represented by a vertical bar, which is partitioned into K colored segments and represents the individual affiliation to each cluster. See supplementary fig. S9. for clustering results for K from 2 to 4. **(D)** *D-statistics* test for gene flow between ancient Chinese domestic cats and *F. bieti* or *F. ornata,* with the analysis setup as *D* (Outgroup, bieti/ornata, PopX, Ancient Chinese domestic cats). The sand cat is used as an outgroup and PopX is labeled on the Y-axis. Results with significant *Z* values are outlined in black, while those with non-significant *Z*-values in red.

This pattern supports a genomic affinity between domestic cats from China and Central Asia and African wildcats from the Levant. The finding is corroborated by ADMIXTURE analyses, where all these individuals share the same ancestry component at *K*=6 (Fig. 3C and fig. S4). Our outgroup *f_3_* analysis is also consistent with these patterns and demonstrates that relative to other ancient Chinese specimens, FS12, the earliest (Fig. 1C and table S2) and westernmost (Fig. 1A) domestic cat sampled in China, shows the strongest genetic association and temporal coincidence with the ancient Dhzankent cat (*14*).

Combined, this evidence suggests that the dispersal of domestic cats towards East Asia occurred via Central Asia before reaching western China. It is possible that this movement was facilitated by trade along the Silk Road since the earliest Chinese domestic cat in our study (FS12) is dated to about 1,200 years ago (706-883 CE) during the Mid-Tang Dynasty, an era in Chinese history when trade linking the East and West along this route was at its peak (*14*,*15*). Since 21.1% (8 of 38 specimens) of contemporary Chinese domestic cats, particularly those from the southern and eastern coastlines, possess mtDNA clade IV-A which is uncommon for cats from northern China (Fig. 2 and table S1), it is possible that cats were also introduced to China via maritime trade routes.

### Phenotypic reconstruction of the earliest domestic cat in China

In numerous domestic taxa, shifts in selective pressures associated with living within anthropogenic environments enabled the emergence of phenotypes that distinguish them from their wild counterparts. These include a loss of camouflage and the appearance of novel coat colors and patterns, though the timing of the origin of different variants varies dramatically (*17*). For example, a high frequency of the distinguishing blotched-tabby coat pattern was evident in domestic cats in Europe only after the Middle Ages (*2*). To reconstruct the phenotype of the oldest domestic cat in China (FS12), we generated a 16× coverage of the nuclear genome (table S1) and assessed known genotypes previously identified in domestic cats (tables S7A-E). We characterized 94 variants associated with a wide range of traits, including coat color and pattern (tables S7A to C), tail type (table S7D), and common genetic diseases identified in modern cat breeds (table S7E).

FS12 was a male without any deleterious mutations associated with feline health defects tested in the study. The wild-type alleles present in seven genes suggested that FS12 had short hair (*FGF5*, *HR*, *KRT71*, *LPAR6*, *LIPH*) (*18-21*) (Fig. 4C and table S7C) and a long tail (*HES7*, *T*) (*22*,*23*) (Fig. 4C and table S7D). The variant associated with the recessive pigmentation allele on *ASIP* (*24*) indicated that FS12 did not possess any black hair (Fig. 4C and table S7A). Furthermore, we identified a *FERV1* insertion within the *KIT* gene that is associated with an all or partially white coat common in domestic cats (*25*) (Fig. 4C, fig. S12, and table S7A). Since we excluded the possibility that FS12 carried the blotched-tabby *Taqpep* allele (*26*) (table S7B), it is possible that this cat was either an all- white cat or a mackerel-tabby cat with white patches (Fig. 4C).

**Figure 4.**
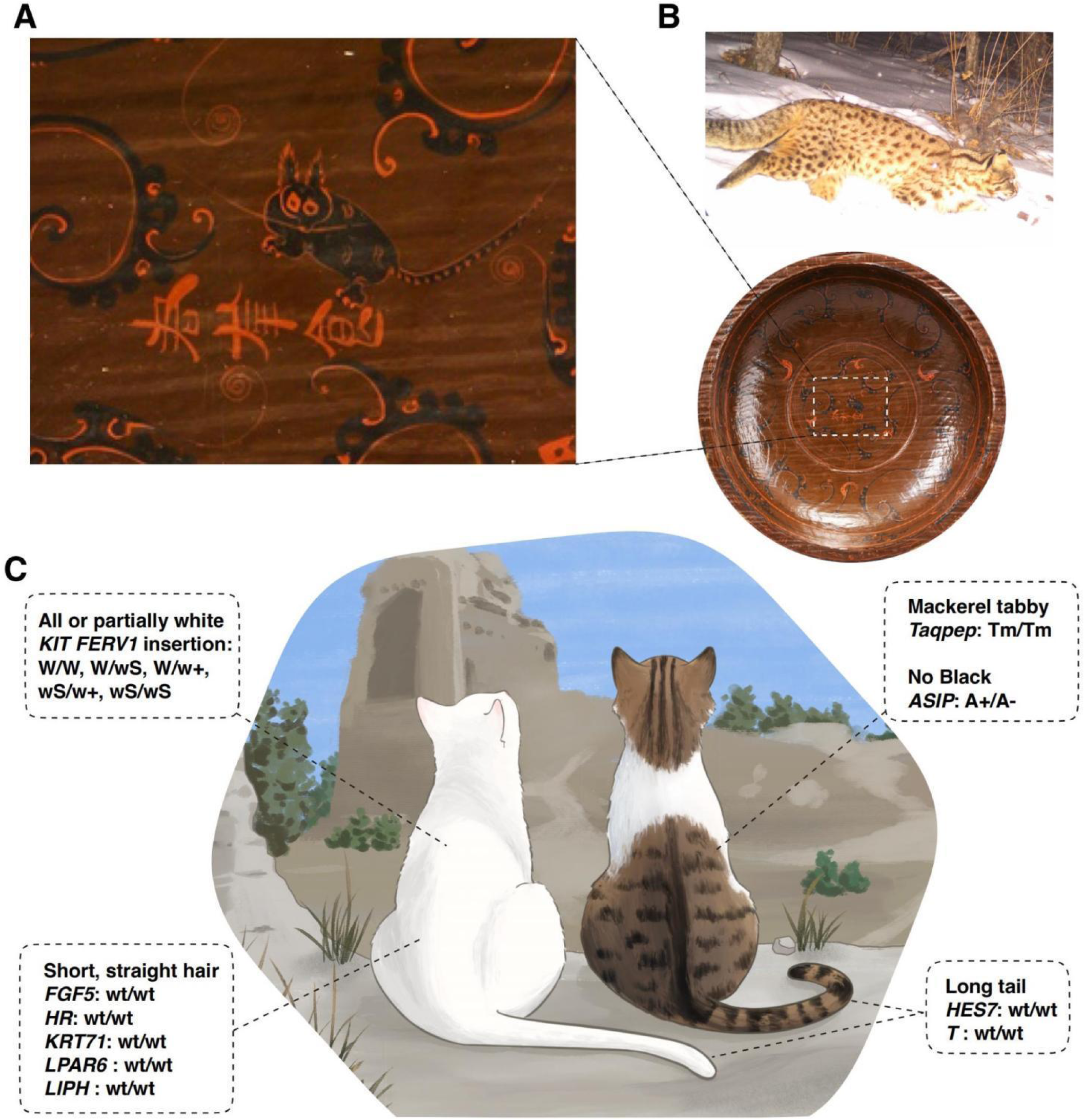
Phenotypic reconstruction of the earliest domestic cat in China and an artistic depiction of a leopard cat-like figure from ancient China. **(A)** Painting of one of the earliest known cat figures on a decorative artifact excavated from the Mawangdui Han Tomb (c.168 BCE), whose distinct spotted coat closely resembles that of a leopard cat (Source: Hunan Museum Collection Database, https://www.hnmuseum.com/zh-hans/zuixin_tuijie). **(B)** A wild leopard cat from Beijing (Credit: School of Life Sciences, Peking University), highlighting the similarity between its spotted coat and the artistic depiction of a cat on the archeological artifact. **(C)** Phenotypic reconstruction, based on whole genome sequencing at 16× coverage, of FS12, the earliest domestic cat discovered in China (706–883 cal CE) from the Tang Dynasty. Genomic inference suggests FS12 was either an all-white cat (left) or a partially white cat with a mackerel tabby pattern (right). The background is an artistic illustration of ancient Tongwan City, Shaanxi, where FS12 was excavated (Image credit: Lanhui Peng).

The presence of white fur in the earliest Chinese domestic cat is consistent with the observation that in East Asia, modern domestic cats possess a greater proportion of white or white-spotting coloration relative to other regions of the world (*27*). This pattern is also reflected in an iconographic survey in which 85% of the cat images identified in 33 ancient paintings from the Tang Dynasty to the 20th century (827–1,930 CE) showed white coat coloration (table S8).

### Assessing gene flow from local wildcat species into ancient Chinese domestic cats

Two Indigenous wildcat species in China, the Chinese mountain cat (*Felis bieti*) and Asiatic wildcat (*F. ornata*), can interbreed with domestic cats and produce fertile offspring (*1,3*). In Europe, analyses of ancient and modern domestic cats and European wildcats (*F. silvestris*) demonstrated that until recently, gene flow between the two taxa was rare (*13*,*28*). In Asia, neither ADMIXTURE (Fig. 3C and fig. S4) nor *D-*statistics (Fig. 3D and fig. S5) revealed evidence for gene flow between the native wildcat species and the ancient domestic cats in China.

This could be because the restricted distribution of the Chinese mountain cat on the eastern part of the Qinghai-Tibetan Plateau does not overlap with the locations where domestic cats were first introduced to China, thus limiting opportunities for interbreeding. Domestic cats likely reached the Qinghai-Tibetan Plateau only within the past centuries and began encountering their alpine relatives. A recent genetic analysis indicated that, as a result of this sympatry, modern domestic cats in this region now possess 1-10% ancestry from Chinese mountain cats (*3*).

### Leopard cats occupied human settlements prior to the arrival of domestic cats

Prior to the arrival of domestic cats, all but one of the seven feline remains found in Chinese archaeological sites were identified as leopard cats (*P. bengalensis*). The only exception is an individual that possessed an Asiatic wildcat (*F. ornata*) mtDNA haplotype (Fig. 1 and fig. S2) found at the site of Huoshiliang (FS7, c.1,800 BCE) (*29*). The earliest archaeological leopard cat specimen (FS15) in our dataset is from the site of Quanhucun dated to ∼5,400 years ago (*6*,*8*) (Fig. 1 and fig. S6). Six additional feline excavations from six archaeological sites dated between c.3,000 BCE and c.150 CE (spanning from the Chinese Late Neolithic to the terminal Han Dynasty) were all identified as leopard cats using both mtDNA (Fig. 1 and fig. S6) and nuclear DNA analyses (figs. S7-S11). The most recently dated individual was a skeleton (FS19) found within human burials at the site of Zheng and Han City in c.150 CE during the late Han Dynasty (*30*,*31*).

The integration of leopard cats into human culture and society is also reflected in ancient Chinese literature (*32*) and art (Fig. 4A). Decorative paintings on a series of culinary artifacts excavated from the Mawangdui Han Tomb (c.168 BCE) feature one of the earliest known cat depictions in Chinese archaeological history (*33*,*34*). These images portray a spotted feline that more closely resembles a leopard cat than a domestic cat (Figs. 1, 4A and B).

Leopard cats may have been attracted to and begun living within human settlements prior to 3,400 BCE (Quanhucun), at least 4,000 years earlier than the arrival of domestic cats (*6*-*8*) (Fig. 1). A previous stable isotopic analysis suggested that leopard cats predated on rodents within human settlements (*6*), implying that this species may have occupied the same anthropogenic niche that was later filled by domestic cats. This relationship lasted at least 3,500 years given that the most recent leopard cat (FS19) in our dataset was dated to c.200 CE (Fig. 1 and table S2). Following this finding, there is a gap of ∼600 years prior to the earliest appearance of a domestic cat (Fig. 1). In fact, over these six centuries, zooarchaeological research in China has thus far failed to identify any small cat remains (*35*). This absence may therefore reflect a real gap during which cats were either absent or extremely rare in human settlements.

The disappearance of leopard cats coincides with a turbulent era in Chinese history after the breakdown of the Han Dynasty and before the Tang Dynasty (Fig. 1). This historical period witnessed a shift toward a colder, dryer climate, dwindling agricultural yields, frequent social conflict, and a demographic contraction that persisted for 400 years (*36*,*37*). These conditions likely contributed to, or were accompanied by, the collapse of the Han Dynasty (*38*), possibly resulting in the decline of agriculture and a contraction of the niche that led to the cohabitation of leopard cats and humans. A study of ancient black rats in Europe demonstrated that, at least in Northern Europe, black rat populations disappeared alongside the demise of the Roman Empire, only to re-emerge from a different source population following the recovery of economic activity (*39*). In each case, the decline of dynasties on both sides of Eurasia may have led to the disappearance of a commensal animal whose presence was dependent on a suitable anthropogenic niche.

The disappearance of leopard cats was followed six centuries later by the arrival of domestic cats during the rise of the Tang Dynasty (*15*). (Fig. 1). It is possible that the arrival of domestic cats may have prevented the re-establishment of the relationship between leopard cats and humans since both species occupy similar ecological niches. Other contributing factors that prevented a second appearance of leopard cats within human settlements may have included the rise of poultry farming in ancient China after the Han Dynasty (*40*), which could have led to human-leopard cat conflict due to the leopard cat’s predation of poultry chicken (*10*,*41*).

## Discussion

The zooarchaeological and genetic evidence demonstrates that the earliest commensal felids found within archaeological contexts were leopard cats. They first appeared at least 5,400 years ago and were then present across China for 3,500 years, during which they were incorporated into human burials and depicted in material culture. Following the end of the Han Dynasty in c.200 CE, leopard cats vanished from both zooarchaeological assemblages and artistic depictions. The disappearance of leopard cats may have begun with the socio-economic disruptions associated with dynastic changes, a shift mirrored in the temporary disappearance of black rats in Northern Europe following the end of the Roman Empire (*39*).

Following a six-century period in which no felid has yet been identified within archaeological contexts, domestic cats arrived in China during the Tang Dynasty during the 8th–9th centuries CE (Fig. 1 and table S2). This result first undermines the commonly held assumption that domestic cats were present in China during either the Neolithic (*6*) or Han Dynasty (*12*). Secondly, this evidence demonstrates that cats were introduced to China far more recently than other Eurasian domestic animals, including cattle, sheep, and goats (c.3,000-c.2,000 BCE) (*42*), and horses (c.1,200 BCE) (*43*). In addition, genomic associations with an ancient Central Asian cat suggest that cats were likely introduced via the Silk Road, thus highlighting the significance of ancient trade routes in facilitating the dispersal of domestic animals.

From a conservation perspective, the introduction of domestic cats has had significant negative consequences for indigenous wildcat species. In Europe, gene flow between the earliest introduced domestic cats and the indigenous European wildcat was minimal (*13*). In the 20th century, however, substantial hybridization between the two taxa has led to the emergence of a hybrid swarm, and the eradication of European wildcats free of domestic cat ancestry at least in Scotland (*28*). Likewise, although we found no evidence of introgression in ancient China between domestic cats and local wildcat species (*F. ornata* and *F. bieti*), contemporary and ongoing admixture between modern domestic cats and *F. bieti* has been reported following the arrival of domestic cats on the Qinghai-Tibetan Plateau (*3*). This pattern raises concerns over the impact of the growing domestic cat population on the genomic integrity of native Chinese wildcats.

By contrast, the leopard cat belongs to *Prionailurus* genus that would not naturally breed with *Felis* species, and human-mediated crossing only occurred in the 1980s, giving rise to the Bengal domestic cat breed (*44*). The evidence presented here hence reveals an intriguing succession of two distinct small felids, each from a separate genus, that independently and consecutively occupied ancient anthropogenic environments in China. Though both were integrated into human cultural expressions in history, they ultimately followed divergent paths in human-animal interactions and reached different destinations.

## Acknowledgments

We thank Yuxin Hou, Yeyizhou Fu, and Zhenyan Sui for their help to the project.

## Author Contributions

Y.H., S.H., G.L., and S.J.L. designed the research; Y.H., J.A., K.L., A.E.J., and H.Y. performed the research; A.M., S.G.M., Y.Z., X.S., D.L., S.H., L.P., X.X., and L.A.F.F. assisted in the laboratory work and data analysis; Y.H., M.Y., C.Y., J.W., Z.W., C.C., X.W., Z.L., W.F., Q.C., Z.C., J.Y., Z.L., Y.L., Z.L., Q.M., J.S., Z.S., W.W., F.X., W.Y., S.Y., L.Y., H.Z., Z.P.L, X.L., and P.Z. carried out fieldwork and sample collection; C.Y., Y.P., and X.W. conducted radiocarbon dating; Y.H., J.A., S.D., G.L., and S.J.L. wrote the manuscript with comments from all authors.

## Data and materials availability

The sequencing data generated in this study are available in the NCBI Sequence Read Archive (PRJNA1178732). The NCBI accession numbers of mitochondrial haplotypes are PQ553222 to PQ553243.

## Supplementary Materials

### Supplementary Results and Discussions

#### Population structure of ancient leopard cats from the Middle Yellow River Basin

The leopard cat has a widespread distribution in Asia and appears to be more tolerant of deforestation and human disturbance than most other Asian wild felids (*1*–*5*), and it often is found in plantations and near villages, showing an affinity to human settlements (*6*–*8*). Previous studies demonstrated that the leopard cat can be divided into two deeply diverging lineages to the level of speciation. or the mainland leopard cat (*P. bengalensis*) and Sunda leopard cat (*P. javanensis*). The mainland lineage contains leopard cats ranging from the Russian Far East to Indochina, while the Sunda leopard cat includes populations from Malay Peninsula and Sundaland. Research on the population structure of leopard cats in the middle reaches of the Yellow River was limited (*5*).

To better understand the population dynamics of ancient leopard cats in this region, we analyzed a dataset consisting of 30 mitochondrial genomes. This included seven newly sequenced ancient samples from the Middle Yellow River Basin and contemporary samples in various regions Russian Far East (n=3), Indochina (n=6), Malay (n=6) and Sundaland (n=3) (table S1D). Additionally, mitochondrial genomes from two fishing cats and a flat-headed cat were included for comparison. For the nuclear DNA analysis, we examined 18 genomes, including three ancient samples from China and 15 modern leopard cats: two from the Russian Far East, six from Indochina, four from Malay, and three from Sundaland. Comparative analysis also included nuclear genomes from a fishing cat and two domestic cats. After SNP calling and filtering, we identified 20,436,894 SNPs in our nuclear genome dataset (table S1).

An ML (Maximum Likelihood) tree was constructed to elucidate the maternal genetic relationships among ancient leopard cats and other populations, as detailed in fig. S6. The analysis divided the leopard cats into two principal clades: Mainland and Insular. Within the Mainland clade, two sub-clades emerged, primarily comprising individuals from the Russian Far East and Indochina, respectively. Notably, our ancient samples from China, though from the same geographical area, were distributed across these sub-clades. FS13, FS14, FS19, and FS24 were grouped with the sub-clade dominated by Indochina individuals, whereas FS8, FS10, and FS15 were aligned with the sub-clade associated with the Russian Far East. This distribution indicates that the ancient leopard cats from the Middle Yellow River Basin shared mtDNA with both Indochina and the Russian Far East, suggesting a complex and interconnected maternal structure in this region during ancient times.

The phylogenetic tree constructed from autosomal SNPs, with robust statistical support for each branch (bootstrap ≥99%), clearly segregates the leopard cat populations into four primary groups: Sundaland, Malay, Indochina, and Russian Far East (fig. S7). Notably, our three ancient samples, FS8, FS15, and FS19, form a monophyletic group closely aligned with the Russian Far East population. This pattern suggests a close relationship between the leopard cats from the Middle Yellow River Basin and those from the Russian Far East.

PCA (fig. S8) demonstrated that the ancient samples form a distinct clade closely associated with both the Russian Far East and Indochina populations along the PC1. Along the PC2, these ancient samples are positioned intermediately between the Russian Far East and Indochina populations, as shown in Figure S8. In the ADMIXTURE analysis (fig. S9), from *K*=2 to *K*=4, the ancient samples consistently show a predominant genetic affinity with the Russian Far East population. This relationship is further supported by the outgroup *f_3_*-statistics (fig. S10; table S3), which indicate that the Russian Far East population shares a more extended evolutionary history with the ancient leopard cats.

Our analyses also point to potential gene flows between the populations of Indochina and the Middle Yellow River Basin. As revealed by the phylogenetic analysis, sample FS19, which clusters with the Indochina clade in the tree constructed using mtDNA, generally aligned with the Russian Far East population in the nuclear DNA analysis (figs. S6-S7). However, in the PCA (fig. S8), ancient Chinese leopard cats were distributing between Russian Far East and Indochina populations. This inconsistency suggested potential genetic exchanges.

To confirm this, we applied *D-*statistics analysis. For the combination (domestic cats, Indochina, Russian Far East, ancient China), the *D* values consistently yielded positive results coupled with significant Z values (|Z| > 3) (fig. S11; table S4). Since our samples from the Middle Yellow River Basin, this results likely suggests that gene flow occurred between ancient population from the Middle Yellow River Basin and the ancestral population of Indochina leopard cats in history.

Combined, these results suggested a significant genetic linkage between the leopard cats from the Middle Yellow River Basin and those from the Russian Far East, indicating they were more closely related genetically than with other contemporary populations. This close relationship may reflect historical gene flow or a shared ancestral population between these regions.

#### Phenotypic reconstruction of the earliest domestic cat of China

A domestic cat remain FS12 was found among the Tang Dynasty’s remains of the Tongwan City in northwest China and directly radiocarbon dated to 706 - 883 CE. It is thus far the earliest domestic cat discovered in China. Comparison of the ratio of the X chromosome to the autosomal chromosomes (approximately 0.5) indicated it as a male individual. FS12 was sequenced to a genome coverage of 16× to infer its phenotypes, including coat color (table S7A), pigmentation pattern (table S7B), fur type (table S7C), tail type (table S7D), and potential genetic diseases (table S7E).

Genomic analysis indicated that FS12 has an all-white or partially white pelage. White coloration in domestic cats is attributed to a feline endogenous retrovirus (*FERV1*) insertion within the *KIT* (*proto-oncogene, receptor tyrosine kinase*) gene’s intron 1. The repercussions of this insertion can lead to body surface depigmentation **(*9*)**. While distinguishing between homozygosity (fig. S12A and C) and heterozygosity (fig. S12A and D) for the *FERV1* insertion within the *KIT* gene is challenging, the distinct alignment patterns observed when compared to the wildtype (fig. S12A and B) suggest that FS12 likely carries either a heterozygous or homozygous *FERV1* insertion (*61*). This genetic configuration is associated with white coloration, indicating that FS12 may exhibit either pure white or white-spotted coat color (Fig. 4C).

Our analysis indicated that FS12 does not possessed black fur, another coat color for domestic cats. This trait stems from a homozygous two-nucleotide deletion (Felis_catus_9.0 GCA_000181335.4, ChrA3:25086566-25086567del) in the *ASIP* gene (*10*). FS12’s heterozygous genotype negates black coloration. We also examined other coloration loci with well-documented genetic mechanisms in FS12, finding that all of these loci are homozygous for the wildtype alleles, which prevents the expression of those specific colors (*11*–*16*). The genomic regions associated with the Mocha (Felis_catus_9.0 GCA_000181335.4, ChrD1:45898609-45898771dup) (*17*) and Dilution (Felis_catus_9.0 GCA_000181335.4, ChrC1:219396820del) (*18*) colorations are absent in FS12, leaving the status of these color patterns inconclusive (table S7A and fig. 4C).. Considering that the Mocha and Dilution colorations are primarily found in Western cat breeds, it is unlikely that FS12 exhibits these two colors.

We assessed the genetic variants associated with the blotched (*19*) and ticked tabby to elucidate FS12’s pigmentation pattern (*20*) (table S7B). Since no mutation associated with the blotched or ticked variant was identified, we infer FS12 to be a wild-type mackerel tabby, akin to that of *Chinese Lihua* or *Dragon Li* cats (fig. S4C; table S7B).

While it is known that FS12 displays white coloration, the current information is inadequate to ascertain the precise extent of the white coverage (*9*). If FS12 possessed a all-white coat, the tabby pattern would not appear due to the all-white coat variant’s epistatic effect. On the other side, FS12 could show a partially white fur with patches of wild-type mackerel tabby pattern. Either coat color morph, an all-white cat or a mackerel tabby cat with white patches, is possible for FS12 (Fig. 4C).

To investigate the hair length morphology, we examined the genetic variants responsible for short hair, long hair, curly hair, hairlessness, and Lykoi-style hair (table S7C) and revealed no variant in FS12 that would result in these specific coat types (*21*–*27*). Although the two mutations in *HR* gene associated with the Lykoi pattern were not available for analysis (*24*), considering that the Lykoi pattern is found in modern Western commercial breeds, it is unlikely to be present in an ancient individual from China. Therefore, we exclude this variation. Based on these findings, we inferred that FS12 has straight and short hair as a wild type (Fig. 4C).

The tail morphology and associated mutations are detailed in table S7D. Since FS12 carries wild-type alleles at all loci (*28*, *29*), it is apparent that it has a full-length, normal tail (Fig. 4C).

We assessed 52 deleterious variants associated with 37 genetic diseases found in modern cats. Detailed information about these can be found in table S7E. No defect variant was found in FS12 (*24*, *30*–*69*).

In summary, phenotypic reconstruction based on 16× coverage of whole genome sequencing data suggests FS12 the earliest domestic cat found in China thus far as an all-white cat or partially white cat with mackerel tabby pattern. It has short hair and a long tail (Fig. 4C; table S7). In addition, no genetic defect commonly found in modern domestic cat breeds are detected in FS12 (table S7E).

We further explored the appearance of cat color in historical artwork with a specific focus on two prominent pelage morphs, the black and white (*70*). We collected 33 historically significant feline-themed paintings spanning from the Tang Dynasty to modern times (827 - 1930 CE) (table S8). Iconographic analysis revealed that 68% of these paintings illustrated cats with black coloration and a staggering 85% portrayed white cats (table S8). Artwork often reflects societal preference, and the high prevalence of white cats in historical Chinese paintings are consistent with an ancient proclivity for white animals in East Asian culture. Phenotypic reconstruction based on genome data and artistic reflection of cat images from ancient artwork jointly confirmed that a preference towards white color may exist historically in China. This preference in feline aesthetics, merging genetics with art, provides a fascinating glimpse into the cultural and biological expression in East Asia. Further research will be essential to illuminate these insights.

### Supplementary Materials and Methods

#### Archaeological sites description

##### Tongwan City Site (Contact: Songmei Hu)

Tongwan City was founded in northwestern China in 413 CE by Helian Bobo, the founder of the Daxia Kingdom. Due to its strategic lofelidion in the heart of the Ordos region, it gradually became a crucial hub for both commercial and military activities within the Inner Asian realm. Its importance was further emphasized by its position along the Silk Road. After the dissolution of Daxia, Tongwancheng continued to be inhabited and used by subsequent periods (*71*). In the present study, a domestic felid sample (FS12) retrieved from an excavation site within the city was subjected to radiocarbon dating. The results indicated that this sample can be dated to 706 - 883 cal CE within the era of the Tang Dynasty.

##### Puyang Tombs (Contact: Juan Wang)

The Puyang is in the Puyang City, Henan Province. The excavation of the site has revealed several burials. Faunal remains identified in these tombs included a variety of species, including the domestic cattle, horse, felid, hare, canid, badger, deer, Chinese zokor, and pheasant. One sample (FS18) from the Tomb M10 was analyzed in this study. This individual was believed to as early as the late Tang Dynasty to the Five Dynasties and Ten Kingdoms (618 - 970 CE) (*72*).

##### Xitucheng City Site (Contact: Zhuang Wu)

The Xitucheng City Site, situated in the westernmost part of Xitucheng Village within Erhao Bu Township, Kangbao County, Zhangjiakou City, Hebei Province, occupies a strategic location. Archaeological evidence suggests that this city site dates back to the Jin Dynasty (1,115 - 1,234 CE), making it approximately 800 years old (*73*, *74*). Five domestic felid samples from the site (labeled FS1, FS2, FS3, FS4, and FS5) have been analyzed in this study.

##### Xinxiang Song Tomb (Contact: Xianzhu Wu)

The tomb is located within the construction project of the new campus of Henan Institute of Technology. Excavation revealed that the site served as a family burial ground during the Northern Song Dynasty, spanning from 960 - 1,127 CE. The four tombs discovered shared a similar architectural style, all mimicking wooden structures with brick chambers. Each tomb consisted of three distinct sections: a burial shaft, an entrance, and a burial chamber. The burial shafts were oriented towards the south, and the stratigraphic accumulation of the burial ground was relatively simplistic. In Tomb 2012XJZM4, apart from the discovery of burial goods such as porcelain and copper coins, a collection of animal bones was also found, including those of a domestic felid. The felid was buried adjacent to the head of the tomb occupant (*75*). This sample has been used in this study labeled as FS6.

##### Zheng and Han City Site (Contact: Ziyi Li)

Zheng and Han City is in the heartland of Central China. Throughout the Spring and Autumn and Warring States periods (770 – 221 BCE), it served as the capital for both the Zheng and Han states successively (Ma, 1978). The felid bones used in this study did not unearthed from a tomb dating to the late Eastern Han period (25 - 220 CE) and radiocarbon dated to 200 - 380 cal CE, while samples FS20, FS21, FS22, and FS23 were derived from pits dating between the Song and Yuan Periods (960-1,368 CE).

##### Zhengzhou Shang City Site (Contact: Juan Wang)

The Zhengzhou Shang City Site is located within Henan Province in Central China and served as the capital city during the Shang Dynasty (*76*, *77*). This region has maintained its usage as a residential area, consistently utilized throughout successive eras. The current study involves a felid bone excavated from a pit radiocarbon dated to the Ming Dynasty (1,413 - 1,480 cal CE) within this site. This sample has been analyzed in this study and is labeled as FS16.

##### Shaliangzi City Site (Contact: Chong Yu)

The Shaliangzi City Site, located in Inner Mongolia’s Hohhot City, encompasses an area of roughly 110,000 square meters. At its center, archaeologists have discovered and excavated a large, single-structure rammed-earth platform dating back to the Han Dynasty. Artifacts recovered from the site primarily consist of architectural materials such as flat tiles, tube tiles, eave tiles, and square bricks with geometric patterns (*78*). A felid skeleton (FS17) was found under the foundation of the granary. Although the main body of the foundation was identified as a creation of the Han Dynasty, radiocarbon dating indicated that the felid sample was originated in the Qing Dynasty (1,671 - 1,944 CE).

##### Quanhucun Site (Contact: Songmei Hu)

The Quanhucun Site is in the village of the same name within Hua County, Shaanxi Province, in Central China. Archaeological remains at the site can be attributed to the Middle to Late Yangshao Culture (4,000 - 2,700 BCE). Agricultural practices of the Yangshao culture encompassed the cultivation of foxtail millet and common millet, and domestication of animals such as pigs and dogs. Studies identified that leopard felids were having a commensal relationship with humans in the site(*6*–*8*). This research incorporated one felid sample (FS15) that can be dated to approximately 5,400 years ago unearthed from the Quanhucun site (*6*–*8*).

##### Wuzhuangguoliang Site (Contact: Songmei Hu)

The Wuzhuang Guoliang archaeological site is situated in the Jingbian County, Shaanxi Province, northwestern China. Occupying about 600,000 square meters, the site spanned across five mountain ridges. The site contains cultural remains ranging from the Neolithic Age to the Warring States period (∼3,000 - 221 BCE) (*7*, *79*). One felid sample (FS8) dated back to the Neolithic Period was analyzed in this research.

##### Huoshiliang Site (Contact: Songmei Hu)

The Huoshiliang Site, situated in the desert interior of Yulin City, Shaanxi Province, northwestern China, spans about 100,000 square meters. A significant number of animal bones were also unearthed. Preliminary analysis identified that the primary cultural content of the Huoshiliang site ranges from the late Longshan period of Chinese Neolithic Age to the Erlitou Period (∼1,800 BCE) (*80*). One felid sample (FS7) unearthed from the site was included in this study.

##### Xinjie Site (Contact: Songmei Hu)

The Xinjie site is a settlement located on the loess plateau on the eastern bank of the Ba River, in the Wei River basin of Shaanxi Province, China. The site encompassed the period from the late Yangshao period to the Longshan period during the Chinese Neolithic Age (∼3,500 - 1,800 BCE), with most of the remains belonging to the late Yangshao period. The site extends approximately 600 meters east to west and about 500 meters north to south, covering a total area of approximately 300,000 square meters. A felid sample (FS14) from the Longshan context was included in this study.

##### Laoniupo Site (Contact: Songmei Hu)

The site of Laoniupo is located at the eastern suburb of Xi’an City, Shaanxi Province, China. It was occupied by many archaeological cultures spanning from the Neolithic period to the Shang Dynasty (∼5,000 - 1,200 BCE) (*81*). One sample (FS10) from the Shang remains of the site was included in this study.

##### Han Chang’an City Site (Contact: Songmei Hu)

The Han Chang’an City is in Weiyang District, Xi’an City, Shaanxi Province, China. It was the capital city of the Western Han Dynasty built and expanded by Emperor Gaozu of Han and his successors on the foundation of the Qin Dynasty’s Xingle Palace. In 2002, the Han Chang’an City Working Group from the Institute of Archaeology, Chinese Academy of Social Sciences, conducted an excavation at the southwest corner site of the Han Chang’an City wall (*82*). During the excavation, a tibia of felid (FS24) was discovered. This sample has been analyzed and radiocarbon dated to 44 cal BCE - 76 cal CE in this research, further confirming its assignment to the Han Dynasty.

##### Yangling Mausoleum (Contact: Songmei Hu)

The Yangling Mausoleum is the joint burial tomb of Emperor Jing of the Western Han Dynasty, Liu Qi, and his queen (157-126 BCE). It is situated in Xianyang City, Shaanxi Province, China. This mausoleum is the easternmost of the emperor tombs located on the Xianyang Plain of the Western Han Dynasty (83). During excavations, a felid skeleton (FS13) was discovered in the external burial pit on the east side of the tomb mound within the mausoleum garden and analyzed in this study. Radiocarbon dating confirmed that the age of the sample is 168 - 48 cal BCE falling into the range of the Western Han Dynasty.

#### Biological specimens

All the specimens newly assembled for this study were obtained from archaeological excavations in collaboration with respective excavators or zooarchaeologists. Each sample has clear records of the archaeological remain or stratigraphic layers from which it was retrieved. This ensures the accuracy of the archaeological context of the samples.

In total, 22 ancient felid bones excavated from 14 archaeological sites dated from ∼3,500 BCE to 1,800 CE were collected (table S1 and Sites Description in Supplementary Materials). From which, 3 nuclear genomes (FS8, FS15, FS19) (3.3× - 6×) and 7 mitochondrial genomes (11.5× - 91×) belong to leopard cats (*P. bengalensis*); 4 nuclear genomes (2.4×-3.5×), 14 mitochondrial genomes (34.7× - 304×) of domestic cats (*Felis catus*) and a mitochondrial genome (16.9×) of Asiatic wild cat (*F. ornata*) or Chinese mountain cat (*F. bieti*) were generated. To obtain phenotypic information of the earliest domestic cats so far found in China. We further deep sequenced FS12 and increased its depth of nuclear genome from 3.5× to16× (table S1A).

In addition to ancient samples newly analyzed in this research, mitochondrial genomes of 42 ancient domestic cats from the British Isle (n=34), Demark (n=5), Near East (n=2) (provided by School of Archaeology, University of Oxford) (*84*), Central Asia (n=1) (*85*) (table S1B), 38 modern domestic cats from China (provided by School of Life Sciences, Peking University) are analyzed jointly in this research. A VCF files containing a total of 43 ancient and modern *Felis* samples (provided by School of Archaeology, University of Oxford and School of Life Sciences, Peking University) (*84*, *86*) were also jointly analyzed with new ancient Chinese samples. The VCF file includes modern Chinese domestic cats (n=5), modern East Asia domestic cats (n=2), ancient Central Aisa domestic cat (n=1), modern Levantine African wildcats (*F. lybica*) (n=3), ancient or historical western Eurasia domestic cats (n=15), modern western Eurasia domestic cats (n=3), ancient European wildcats (*F. silvestris*) (n=3), modern European wildcats (n=4), modern Asiatic wildcats (*F. ornata*) (n=4), modern Chinese mountain cats (*F. bieti*) (n=5), and modern Sand cats (*F. margarita*) (n=2) (table S1C).

In addition to *Felis* species, modern mitochondrial genomes of 18 leopard cats (*P. bengalensis*) from the Russian Far East (RFE, n=3), Indochina (n=6), Malay (n=6) and Sundaland (n=3) and a fishing cat (*P. viverrinus*) were jointly analyzed with ancient leopard cats. Together with 3 nuclear genomes of ancient Chinese leopard cats, modern nuclear genomes of 15 leopard cats from RFE (n=2), Indochina (n=6), Malay Peninsula (n=4), Sundaland (n=3). Together with this dataset one fishing cat and two domestic cats are also combined and analyzed with the ancient leopard cats from China. All leopard cats and fishing cat data were originally generated and provided by School of Life Sciences, Peking University.

#### Radiocarbon dating

Six samples were submitted for accelerator mass spectrometry (AMS) dating. Three samples (FS16, FS17, and FS19) were dated at *Beta Analytic* in the United States and the remaining samples (FS12, FS13, and FS24) were processed at the School of Archaeology, Peking University, China. All radiocarbon dates reported were calibrated using the INTCAL20 calibration curve (*87*).

#### Ancient DNA laboratory procedures

DNA extraction was performed in the dedicated ancient DNA laboratory at the School of Life Sciences, Peking University. All sterilized centrifuge tubes and other plastic consumables used in the following procedures were sterilized using a UV crosslinker with high-intensity UV radiation for 15 minutes. In this study, we utilized the Dabney method for ancient sample DNA extraction and optimized it further.

##### Sample preparation

Bone samples, weighing between 100 - 250 mg, were prepared using a Dremel multi-tool drill. The bone surface was carefully removed with a circular cutting disk to eliminate any exogenous DNA. To minimize the risk of cross-contamination, the drill was cleaned with a 10% bleach solution, and the spindle and disposable disk were replaced after processing each sample. The benchtop was also cleaned with 10% bleach and 75% ethanol between each sample, and gloves were changed frequently. Negative controls were employed as blanks for each batch of tested samples at every stage of the laboratory procedures, from extraction to capture. The bones were pulverized using a hammer. Any equipment that had come in contact with the samples was thoroughly cleaned between sample processing using 10% bleach and 75% ethanol.

##### DNA Extraction

All samples were processed using Dabney method (*88*) with slight modification by adding a pre-treatment step. Initially, samples were pre-treated to remove external DNA contamination, which included pre-digestion (*89*). For this step, 1 mL of pre-digestion premix containing EDTA and proteinase K (see Table A.1) was added to the sample tube. The sample tube was then placed in a four-dimensional rotary mixer located inside a constant temperature oven and incubated at 37°C for 10 minutes with rotation; this was followed by centrifugation at 12,000 rpm for 1 minute and subsequent removal of the supernatant.

Following pre-treatment, samples underwent extended digestion using a digestion premix containing EDTA and proteinase K to release internal DNA. DNA extraction was then completed using the MinElute PCR Purification Kit according to the manufacturer’s instructions. The specific steps were as follows: 4 mL of digestion premix was added to the 5 mL centrifuge tube containing the pre-treated sample, and the sample was incubated overnight at 55°C with rotation (16 - 24 hours). The next day, MinElute spin columns, sterile plastic funnels, and 50 mL sterile centrifuge tubes were assembled. 10 times the volume of the digestion fluid of weakly acidic PB buffer was added to another batch of 5 mL sterile centrifuge tubes. The tube containing the digestion fluid was centrifuged at 12,000 rpm for 1 minute, and the supernatant was transferred to a 50 mL centrifuge tube and thoroughly mixed with the binding buffer. The mixture of digestion fluid and binding buffer was then transferred in batches to the assembly tube using a sterile 5 mL pipette and centrifuged at 400 g at 4°C until the mixture had completely passed through the MinElute spin column. After the transfer was complete, the MinElute spin column was removed from the assembly tube and placed in a collection tube and centrifuged at 12,000 rpm at 4°C for 1 minute. Then 700 μL of PE buffer was added to the MinElute spin column, centrifuged at 8,000 g for 1 minute, and the waste liquid was discarded. This step was repeated once. A dry spin at 12,000 rpm for 1 minute was performed; the MinElute spin column was then transferred to a 1.5 mL DNA LoBind tube, 50 μL of EB buffer was added to the filter membrane of the MinElute spin column and incubated at 37°C for 10 minutes. The eluate was collected by centrifuging at 12,000 rpm for 1 minute, the MinElute spin column was discarded, and the DNA extract was stored at 4°C.

##### Library construction

This study employed the Blunt-End Single-Tube double-stranded library building method (BEST protocol) (*90*). The BEST protocol involves three main steps: end repair, adaptor ligation, and fill-in. First, 16 μL of DNA extract is added to a 0.2 mL PCR tube and diluted with 16 μL of EB buffer. Then, 8 μL of end repair premix is added, mixed thoroughly, and subjected to a PCR program of 30 minutes at 20 °C followed by 30 minutes at 65°C. For adaptor ligation, the appropriate concentration and volume of adaptor for each sample is calculated based on the concentration of the DNA eluate, with volumes not exceeding 2 μL; if less is needed, EB buffer is used to make up the volume. Then, 8 μL of ligation enzyme premix is added, mixed thoroughly, and subjected to a PCR program of 30 minutes at 20°C followed by 10 minutes at 65°C. For the fill-in step, 10 μL of fill-in premix is added and the mixture is incubated in a PCR machine for 15 minutes at 65°C followed by 15 minutes at 80°C. The DNA is then purified according to the SPRI bead protocol, eluted in 34 μL of EB buffer, and collected in a 1.5 mL DNA LoBind tube, stored at 4°C.

##### qPCR

For quantification and amplification of the DNA library, 1 μL of DNA library liquid and 19 μL of qPCR premix are taken in a new 0.2 mL PCR tube and subjected to a qPCR program. Based on the qPCR amplification curve, suitable indexing PCR cycle numbers are selected for each sample; 15 μL of DNA library liquid and 35 μL of Indexing PCR premix are mixed and subjected to the Indexing PCR program. The amplified DNA library is purified using SPRI beads, eluted in 40 μL of EB buffer, and collected in a 1.5 mL DNA LoBind tube. The concentration of the amplified DNA library liquid is determined using the Qubit 4.0 fluorometer as per the manufacturer’s instructions.

##### Library amplification

Libraries was subsequently amplified and externally indexed (*91*)using standard Illumina P5 and P7 primers with double indexes in a 50 μL PCR reaction, consisting of 15 μL of library, 25 μL Gold Taq 360 Master Mix, 5 μL Reaction Enhancer, 3 μL H_2_O, 1 μL of P5 primer, and 1 μL of P7 primer (10nM). Thermocycling conditions included an initial denaturation step of 2 minutes at 95°C, followed by cycles of 15 seconds denaturation at 95°C, 30 seconds annealing at 60°C, and 30 seconds extension at 72°C for the specified number of cycles for each sample. A final extension step of 7 minutes at 72°C was conducted. The amplified libraries were purified using SPRI beads (*Beckman Coulter*), and quality control was performed on a Fragment Analyzer (*Spectro Scientific*) prior to sequencing.

##### Sample screening

All samples underwent screening sequencing on *NovaSeq* 6000 instrument (*92*) at Novogene, China. Approximately 3 Gb of raw data were obtained for each sample. Following the screening sequencing, the ratio of nucleotides mapped to the domestic cat whole nuclear reference genome (Felis_catus_9.0 GCA_000181335.4) (endogenous DNA) and the number of reads mapped to the mitochondrial genome (NC_001700.1) were calculated. For leopard cat samples we estimated endogenous DNA content by calculating the ratio of nucleotides that mapped to the leopard cat reference genome (Genome assembly Fcat_Pben_1.1_paternal_pri GCF_016509475.1) or the number of reads mapped to the leopard cat reference mitochondrial genome (NC_028301.1). We then determined further sequencing strategies based on these results.

##### Whole genome resequencing

Samples representing the spatial and temporal contexts or possessing more than 10% endogenous DNA in screening were selected for deeper sequencing. In total, four domestic cats (FS1, FS2, FS12, FS21) and three leopard cats (FS8, FS15, FS19) are deep sequenced. All samples were sequenced on a *NovaSeq* 6000 instrument at Novogene, Beijing. We obtained approximately 100 Gb of data each for FS1, FS2, FS12, and FS22. FS8, FS15, FS19 are processed in a whole lane. To reveal the phenotypic information of FS12, the most ancient domestic cats so far found in China, we further sequenced this sample in a single lane of *NovaSeq* 6000 instrument at Novogene, Beijing, and obtained an additional approximately 900 Gb data.

##### Mitochondrial genome sequencing

An additional 4 Gb of sequencing data were obtained to acquire full mitochondrial genomes for samples that did not qualify for whole nuclear genome sequencing but had more than 2,000 reads mapped to the mitochondrial reference genome (NC_001700.1). Samples FS3, FS4, FS5, and FS17 were processed using this strategy and were sequenced on an Illumina *NovaSeq* 6000 instrument at Novogene, China.

##### Ancient mitochondrial DNA capture sequencing

In-solution targeted capture of the cat mitochondrial genome was conducted on 14 domestic cat samples (FS6, FS7 FS8, FS10, FS13, FS14, FS15, FS16, FS17, FS18, FS19, FS20, FS21, FS22, FS23) using the *myBaits* (*Arbor Biosciences*) Hybridization Capture for Targeted Next-Generation Sequencing protocol (*93*), with a hybridization temperature of 55°C and a hybridization time of 48 hours.

The domestic cat mitochondrial reference genome (NC_001700.1) was employed to synthesize baits for capture. Due to the degraded nature of ancient DNA (*94*, *95*), the amount of endogenous DNA per sample is considerably lower than that of modern samples, resulting in the potential for probes in a single capture reaction to be significantly overwhelmed for each sample. To achieve optimal cost-effectiveness, more than one sample was pooled per reaction. Consequently, three reactions, each containing 2-3 ancient libraries pooled equimolarly, were prepared. The libraries combined in a single pool had distinct indexes. The three reactions were then sequenced on an Illumina *NovaSeq* 6000 instrument at Novogene, China.

#### Genomic data processing

##### Raw data processing

Raw sequencing data were processed by trimming adapters with default setting and merges completely overlapping paired-end reads into a single sequence (--collapse) using AdapterRemoval v2.3.0 (*96*). Subsequently, Cutadapt v3.5 (*97*) was employed to remove 5 bases from both the 5’ and 3’ ends (-u 5 -u -5) of the reads to minimize the impact of deamination (*94*, *95*). The same software was also utilized to trim low-quality reads shorter than 25 bp (--minimum-length 25) and containing over 10% Ns (--max-n 0.1).

##### Mitochondrial genome data mapping and filtering

###### Leopard cats

The filtered data of leopard cats were mapped to the leopard cat mitochondrial reference genome (NC_028301.1) using the “aln” function in BWA v0.1.17 (*98*) with specifying the maximum allowed mismatch ratio to be 0.01(-n 0.01), allowing for up to two gap openings (insertions or deletions) in a read (-o 2), and a seed length to be 1024 (-l 1024). SAMtools v.1.14 (*99*) were used to sort the bam files with “sort” function, and to filter out PCR duplicates by “markdup” function with a parameter “-r” added. Unmapped reads and reads with a mapping quality less than 30 were removed by the parameters “-F 4”, and “-q 30” of the “view” function in SAMtools.

###### Domestic cats

The filtered data of domestic cats were then mapped to the leopard cat mitochondrial reference genome (NC_001700.1) using the “aln” function in BWA v0.1.17 (H. Li, 2013) with specifying the maximum allowed mismatch ratio to be 0.01(-n 0.01), allowing for up to two gap openings (insertions or deletions) in a read (-o 2), and a seed length to be 1024 (-l 1024). SAMtools v.1.14 (*99*) were used to sort the bam files with “sort” function, and to filter out PCR duplicates by “markdup” function with a parameter “-r” added. Unmapped reads and reads with a mapping quality less than 30 were removed by the parameters “-F 4”, and “-q 30” of the “view” function in SAMtools.

HTSbox (https://github.com/lh3/htsbox) was used to call consensus mitochondrial sequences with the default settings. Control regions of the consensus sequences were then excluded manually, and the remaining sequences were then aligned to each other using MUSCLE v.5 (*100*) with default setting.

##### Nuclear genome data mapping

###### Leopard cat

The filtered data of leopard cats was then mapped to the leopard cat whole nuclear reference genome (Genome assembly Fcat_Pben_1.1_paternal_pri GCF_016509475.1) using the “aln” function in BWA v0.1.17 (*98*) with specifying the maximum allowed mismatch ratio to be 0.01(-n 0.01), allowing for up to two gap openings (insertions or deletions) in a read (-o 2), and a seed length to be 1024 (-l 1024). SAMtools v.1.14 (*99*) were used to sort the bam files with “sort” function, and to filter out PCR duplicates by “markdup” function with a parameter “-r” added. Unmapped reads and the reads with a mapping quality less than 30 were removed by the parameters “-F 4”, and “-q 30” of the “view” function in SAMtools.

###### Domestic cat

Filtered data of domestic cats were mapped to the domestic cat nuclear reference genomes (Felis_catus_9.0 GCA_000181335.4) using the “aln” function in BWA v0.1.17 (*98*) with specifying the maximum allowed mismatch ratio to be 0.01(-n 0.01), allowing for up to two gap openings (insertions or deletions) in a read (-o 2), and a seed length to be 1024 (-l 1024). SAMtools v.1.14 (*99*) were used to sort the bam files with “sort” function, and to filter out PCR duplicates by “markdup” function with a parameter “-r” added. Unmapped reads and the reads with a mapping quality less than 30 were removed by the parameters “-F 4”, and “-q 30” of the “view” function in SAMtools.

###### Validation of ancient data

To evaluate the authenticity of the ancient DNA samples, we employed the mapDamage v2.0 (*101*, *102*). During the initial processing of our data, we trim the first and last 5 base pairs (bp) from each read to avoid the effects of deamination. However, this practice contradicts the principle behind using mapDamage (*101*, *102*), which is designed to show signals of ancient DNA damage. Therefore, for each sample, we also prepare a version with the ends untrimmed, and we use this version for analysis with mapDamage. The mapDamage analysis revealed clear and distinct signals of deamination on the 5’ and 3’ ends of all 22 ancient Chinese samples. The observed C-to-T or A-to-G misincorporations were consistent with known patterns of ancient DNA damage, providing strong evidence for the authenticity of the ancient DNA in these samples (Fig. S1).

##### SNP variant calling

###### Leopard cat

The genomic data of three ancient leopard cats, 15 modern leopard cats from Russian Far East (n=2), Indochina (n=6), Malay (n=4), and Sundaland (n=3), and one fishing cat, in addition to two domestic cats as outgroup, were used to generate SNPs. The SNPs were identified using GATK v.4.2 (*103*–*105*). Subsequently, the variants were filtered. Bcftools v 1.15.1 (*99*) “filter” was used to filter out all transition-type single nucleotide variants, retaining only transversion-type single nucleotide variants to avoid analytical errors caused by cytosine deamination. The variant sites were further filtered using Plink v1.90b6.26 software (*106*), including keeping biallelic SNPs only (--biallelic- only) setting a minimum allele frequency with ’--maf 0.01’, a missing genotype rate with ’--geno 0.2’, and filtering for linkage disequilibrium (LD) by pruning with a window of 10kb, step of 2 SNPs, and an *R*^2^ threshold of 0.5 with ’--indep-pairwise 10K 2 0.5’. Only the autosomal were kept for downstream analysis. After filtering, we obtained the SNP dataset covering 18 autosomes of leopard cats. Through all these steps, a total of 20,436,894 SNPs were called and used for downstream analysis.

###### Domestic cat

SNPs of 4 ancient Chinese domestic cats were called with GATK v.4.2 (*103*–*105*). Subsequently, the variants were filtered following the methods as described in Jamieson et al 2023. Specifically, BEDtools v.2.30.0 (*107*) was employed to exclude all non-neutral loci. Bcftools v 1.15.1 (*99*) “filter” was used to filter out all transition-type single nucleotide variants, retaining only transversion-type single nucleotide variants to avoid analytical errors caused by cytosine deamination. The variant sites were further filtered using Plink software v1.90b6.26, including keeping biallelic SNPs only (-- biallelic-only) setting a minimum allele frequency with ’--maf 0.01’, a missing genotype rate with ’--geno 0.2’, and filtering for linkage disequilibrium (LD) by pruning with a window of 10kb, step of 2 SNPs, and an *R*^2^ threshold of 0.5 with ’--indep-pairwise 10K 2 0.5’. After filtering, we obtained the SNP dataset covering 825,403 SNPs in 18 autosomes of domestic cats.

VCF files containing a total of 43 ancient and modern *Felis* samples (*84*, *86*) was then merged with our ancient data. We eventually get a VCF file containing ancient Chinese domestic cats (n=4) modern Chinese domestic cats (n=5), modern East Asia domestic cats (n=2), ancient Central Aisa domestic cat (n=1), modern Levantine African wildcats (*F. lybica*) (n=3), ancient or historical Western Eurasian domestic cats (n=15), modern western Eurasia domestic cats (n=3), ancient European wildcats (*F. silvestris*) (n=3), modern European wildcats (n=4), modern Asiatic wildcats (*F. ornata*) (n=4), modern Chinese mountain cats (*F. bieti*) (n=5), modern sand cats (*F. margarita*) (n=2). Through these steps, a total of 825,403SNPs were generated for downstream analysis.

##### Species identification through phylogenetic clustering

A maximum likelihood phylogenetic tree (ML) was performed using IQ-tree v2.3.6 (*108*, *109*) to identify the species of the samples. This analysis included all 22 ancient samples, and modern mitochondrial genomes of domestic cats, wild cats, and leopard cats, with the tiger (NC_010642.1) chosen as the outgroup. We implemented the Tamura-Nei 93 (TN93) nucleotide substitution model (*110*), incorporating both a gamma distribution (G) and a proportion of invariant sites (I) (-m TN93+I+G) following previously published procedure (*86*). The bootstrap replicates were set to be 1,000 (-B 1000). The trees were visualized using FigTree v1.4.4 (http://tree.bio.ed.ac.uk/software/figtree/) and iTOL v5 an online tool for tree visualization (*111*, *112*).

##### Population genetic and phylogenetic analyses of leopard cats

A Maximum likelihood phylogenetic tree was constructed using IQ-tree v2.3.6 (*108*, *109*). The tree included 7 ancient samples and 20 modern leopard cats, comprising four individuals from Russian Far East, 6 individuals from Indochina, 4 individuals from Sundaland, and 6 from Malay. In addition to leopard cats, 2 fishing cats, and a flat- headed () cat were used for comparison. A tiger (NC_010642.1) was used as outgroup. The tree construction using TN93+I+G model (-m TN93+I+G) with 1000 bootstrap replicates (-B 1000). The trees were visualized using FigTree v1.4.4 (http://tree.bio.ed.ac.uk/software/figtree/).

For whole-genome data, we used the vcf2aln software (*113*) to convert the VCF format variation information files of ancient and modern samples into aligned FASTA sequence files to fit the requirement of input file format of IQ-tree v2.3.6 software (*108*, *109*). We set the parameter “-s” to ignore missing site information and “-O” to print only one haplotype for diploid data. A ML (maximum likelihood) phylogenetic tree was constructed based on the 20 million SNPs with IQ-tree. The bootstrap was set to 200 replicates (-B 200). The best model was automatically found by the IQ-tree software by adding “-m MF” A total of 3 ancient leopard cats, 18 samples, including 2 Chinese domestic cats and 15 modern leopard cats from Russian Far East (n=2), Indochina (n=6), Malay (n=4), Sundaland (n=3), and a fishing cat were included in the analysis. Domestic cat was used as outgroup. The trees were visualized using FigTree v1.4.4 (http://tree.bio.ed.ac.uk/software/figtree/).

Principal component analysis (PCA) was performed using SmartPCA v16000 in EIGANSTRAT v7.2.1 (*114*, *115*) with a total of 18 samples, including 3 ancient Chinese leopard cats and 15 modern leopard cats from Russian Far East (n=2), Indochina (n=6), Malay (n=4), and Sundaland (n=3), along with a fishing cat. Due to the large amount of missing data in ancient samples that could impact PCA results, the PCA was first constructed using high-quality modern data. The ancient leopard cats were then projected onto it by applying the “LsqProject: Yes” function.

ADMIXTURE analysis was performed using ADMIXTURE 1.3.0 (*116*) on over 20 million SNPs derived from a total of 18 samples, which included 3 ancient Chinese leopard cats, 12 modern leopard cats from Russian Far East (n=2), Indochina (n=6), Malay (n=4), Sundaland (n=3), and a fishing cat. We performed the analysis with K value from 1 to 8. Three duplications (CV=3) for each K value. The convergence threshold was set to 0.0001 (-c 0.0001). The results were visualized using an R script.

Outgroup *f_3_*-statistics were conducted through the qp3Pop method in EIGENSTRAT v7.2.1 (*115*, *117*) to test the gene flow between Indochina population and ancient leopard cats from China. Over 20 million SNPs from a total of 20 samples, including 3 ancient Chinese leopard cats, 15 modern leopard cats from Russian Far East (n=2), Indochina (n=6), Malay (n=4), and Sundaland (n=3), a fishing cat, and two domestic cats, were used. Domestic cat was used as the outgroup.

*D*-statistics analysis was conducted using *qpDstat* method in EIGENSTRAT v7.2.1 (*115*, *117*). The same dataset as the outgroup *f_3_*-statistics was used. The population combinations were set as *D* (Outgroup, P.bengal_X, Russian Far East, ancient China) mainly to test the potential gene flow between the leopard cats in ancient China and Indochina. Domesttic cats were used as the outgroup. Leopard cats from Indochina, Malay and Sundaland are then individually placed in P.bengle_X.

##### Population genetic and phylogenetic analyses of domestic cats

Maximum Likelihood (ML) phylogenetic analysis of whole mitochondrial genomes was used 14 ancient Chinese domestic cats, 42 ancient or historical domestic cats from western Eurasia, 1 ancient Domestic cat from central Asia, 50 modern domestic cats from China, 2 modern domestic cats from Central Asia, 3 modern African wildcats from the Levant, and 1 modern domestic cat from Western Eurasia. The phylogenetic analysis was performed using IQ-tree v2.3.6 software (*108*, *109*), implementing TN93+I+G model (-m TN93+I+G) with 1,000 replicates (-B 1000). The trees were visualized using FigTree v1.4.4 (http://tree.bio.ed.ac.uk/software/figtree/). Phylogeographical assignment of mtDNA haplotype is conducted as previously described (*84*, *118*).

Using 825,403 SNPS, PCA were performed with SmartPCA in EIGENSTRAT v7.2.1 (*114*, *115*). To elucidate high-resolution geographic patterns among different domestic cat populations; therefore, European wildcats, Asiatic wild cats and Chinese mountain cats were excluded from the analysis. As African wild cats are the progenitors of domestic cats, they were retained. In total, 4 ancient Chinese domestic cats, 5 modern Chinese domestic cats, 2 modern East Asia domestic cats, 1 ancient Central Asia domestic cat, 3 modern Levantine African wildcats, 15 ancient or historical Western Eurasia domestic cats, 3 modern western Eurasia domestic cats were included in this analysis. Since ancient samples may contain a considerable amount of missing data, potentially impacting PCA results, the “LsqProject: Yes” function was adopted to project all ancient data onto the PCA initially defined by high-quality modern data. The results were visualized with a R ADMIXTURE analysis employed 825,403 SNPS. The assay was conducted using ADMIXTURE 1.3.0 (*116*). We performed the analysis with K value from 2 to 7. Three duplications (CV=3) for each K value and selected K = 6 based on the cross-validation (CV) error. The convergence threshold was set to 0.0001 (-c 0.0001). The results were visualized using an R script. This analysis has included 4 ancient Chinese domestic cats, 5 modern Chinese domestic cats, 2 modern East Asia domestic cats, 1 ancient Central Aisa domestic cat, 3 modern Levantine African wildcats, 15 ancient or historical Western Eurasia domestic cats, 3 modern western Eurasia domestic cats, 3 ancient European wildcats, 4 modern European wildcats, 4 modern Asiatic wildcats, 5 modern Chinese mountain cats, and 2 modern Sand cats. The results were visualized with a R

Outgroup *f_3_*-statistics in EIGENSTRAT v7.2.1(*115*, *117*) was utilized to inform the genetic affinity among different populations of domestic cats. Sand cat (*F. margarita*) was selected as the outgroup.

Subsequently, qp*D*stat in EIGENSTRAT v7.2.1 (*115*, *117*) was employed to further explore potential admixtures between ancient Chinese individuals and Asiatic wildcats or Chinese mountain cats (*D*(Outgroup,, ancient European domestic cat, ancient Chinese domestic cat)), where sand cat was used as outgroup. This analysis has included 4 ancient Chinese domestic cats, 5 modern Chinese domestic cats, 2 modern East Asia domestic cats, 1 ancient Central Asia domestic cat, 3 modern Levantine African wildcats, 15 ancient or historical Western Eurasia domestic cats, 3 modern western Eurasia domestic cats, 3 ancient European wildcats, 4 modern European wildcats, 4 modern Asiatic wildcats, 5 modern Chinese mountain cats, and 2 modern Sand cats.

##### Phenotypic inference of the earliest domestic cat of China

Whole genome sequencing of FS12 for a 16x genome coverage was obtained and mapped to the domestic cat nuclear reference genome (Felis_catus_9.0 GCA_000181335.4) and filtered using the same methods as mentioned above using BWA v0.1.17 (*98*) and SAMtools v.1.14 (*99*). The resulted BAM file for FS12 is then used for downstream analysis for phenotypic inference.

In determining the sex of the specimen FS12, we ascertained the sequencing depths of its X chromosome and autosomes using SAMtools v.1.14 (*99*), followed by calculating the ratio of sequencing depth between the two. A ratio near 1 suggests a female individual, whereas a ratio approximating 0.5 indicates a male.

We procured the reference sets for phenotype-related variants of domestic cats from the database *Online Mendelian Inheritance in Animals* (OMIA) (*119*) (https://omia.org/home/). Incorporated within these variants are 61 single nucleotide polymorphisms (SNPs) and 31 deletions or insertions (Indels), associated with thirteen coat colors, five coat forms, three pigmentation patterns, two tail type variations, and 37 genetic diseases (Table S7). Next, from the generated alignment files, we extracted the specific genomic locus related to a particular phenotype, as well as the 200bp regions before and after this locus, using SAMtools v.1.14 (*99*) with the command “view -h -b”. Afterwards, we used BCFtools v 1.15.1 (*99*) with the command “mpileup -B -C50 -q 30 - Q 30” to obtain high-quality genotype data for the corresponding regions. Only loci with a sequencing depth greater than 10x were considered reliable.

With respect to mutations in the *KIT* gene responsible for white pigmentation (*9*) given that the domestic cat reference genome (Felis_catus_9.0 GCA_000181335.4) lacks the retroviral (*FERV1*) insertion, aligning FS12 to this reference posed a challenge in discerning its genuine genotype. To solve the problem, we manually constructed an reference genome, integrating the retroviral (*FERV1*) fragment into the domestic cat’s *KIT* gene (refer to KIT_FERV1_10K). The raw data of FS12 was subsequently aligned to the KIT_FERV1_10K with the same methods as mentioned above using BWA. The outcome is then visualized using IGV v2.16.2. For comparative purposes, we also sourced individuals exhibiting homozygosity, heterozygosity for the *KIT* white-related mutation, and wildtype individuals. Using BWA v0.1.17 mem (*120*) and SAMtools v1.14 (*99*), these specimens were aligned against KIT_FERV1_10K. The results are visualized with the same method as FS12. The alignment pattern of FS12 is then compared to the wildtype, homozygosity and heterozygosity individuals to determine the genotype of its own.

#### Assembly and analysis of cat images in ancient artwork from China

Artwork from a certain historical period may to some extent reflect the aesthetic view and preference from contemporaneous society. We examined the coat color of domestic cats displayed in ancient paintings from China to evaluate the anthropogenic preference of specific cat’s coat color, with a particular focus on the white cat.

Historical paintings were assembled from published archaeological reports and the digital repository Artlib (http://www.artlib.cn). The dataset includes 91 domestic cats from 33 pieces of paintings, covering a historical period from the Tang Dynasty to modern time (827 - 1933 CE). Coat color of the cats was discerned through visual assessment. A primary focus was to discern cats with black and white color and to calculate the proportion of cats bearing these specific coat color.

## Supplementary Figures

**Fig. S1.**
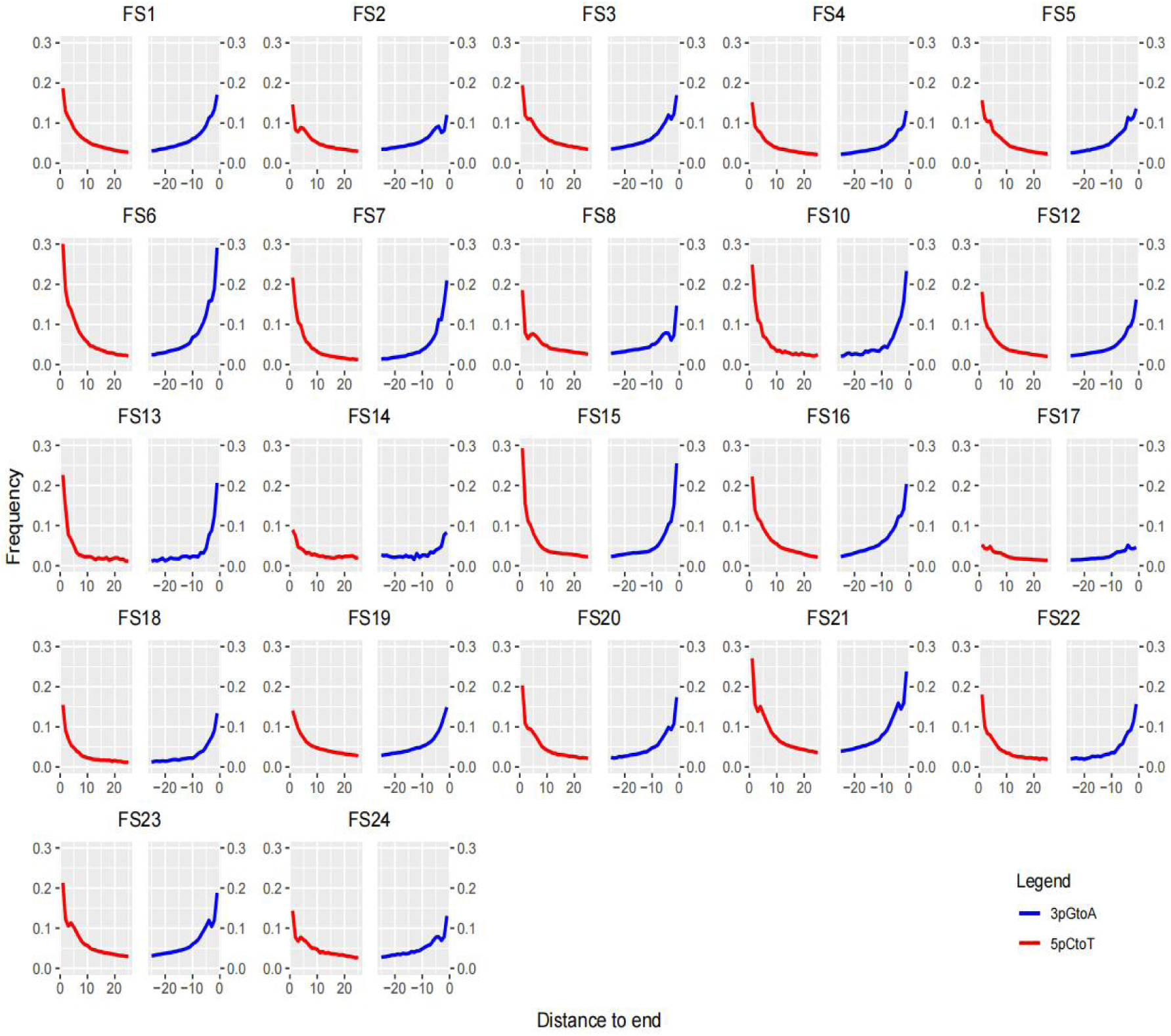
Plots of mapDamage analysis showing damage patterns of 22 ancient Chinese cat specimens used in this study. The 5’ and 3’ ends of all samples exhibit a clear deamination pattern signature for ancient DNA.

**Fig. S2.**
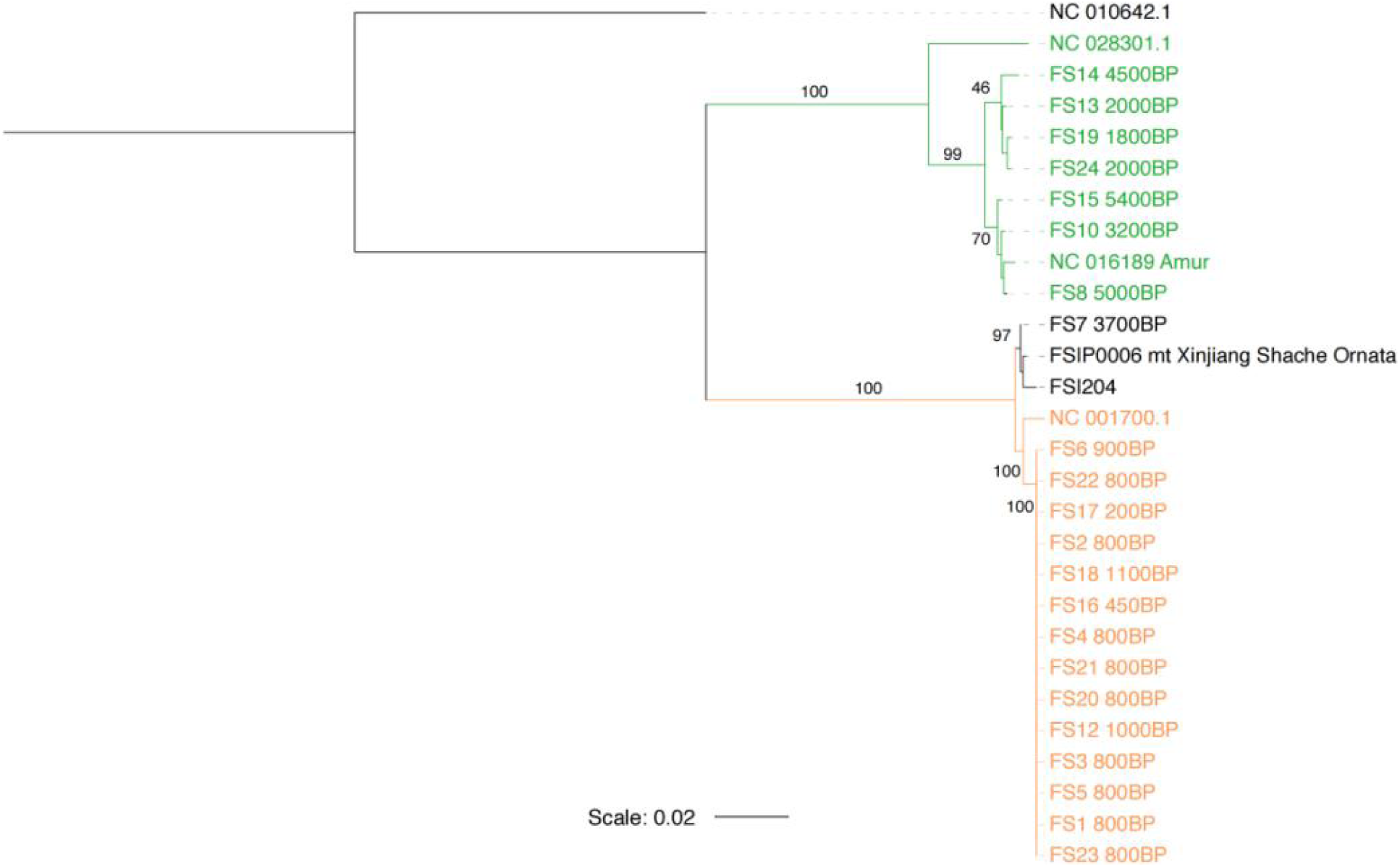
A maximum likelihood (ML) phylogenetic tree constructed based on the mitochondrial genomes of 22 ancient felid samples collected from China. A TN93+I+G model is implemented with 1,000 bootstrap replicates and a tiger genome (NC_010642.1) is used as an outgroup. The numbers on branches represent the statistical support values based on 1,000 bootstrap replicates. A refined version of the tree is in Fig. 1B.

**Fig. S3.**
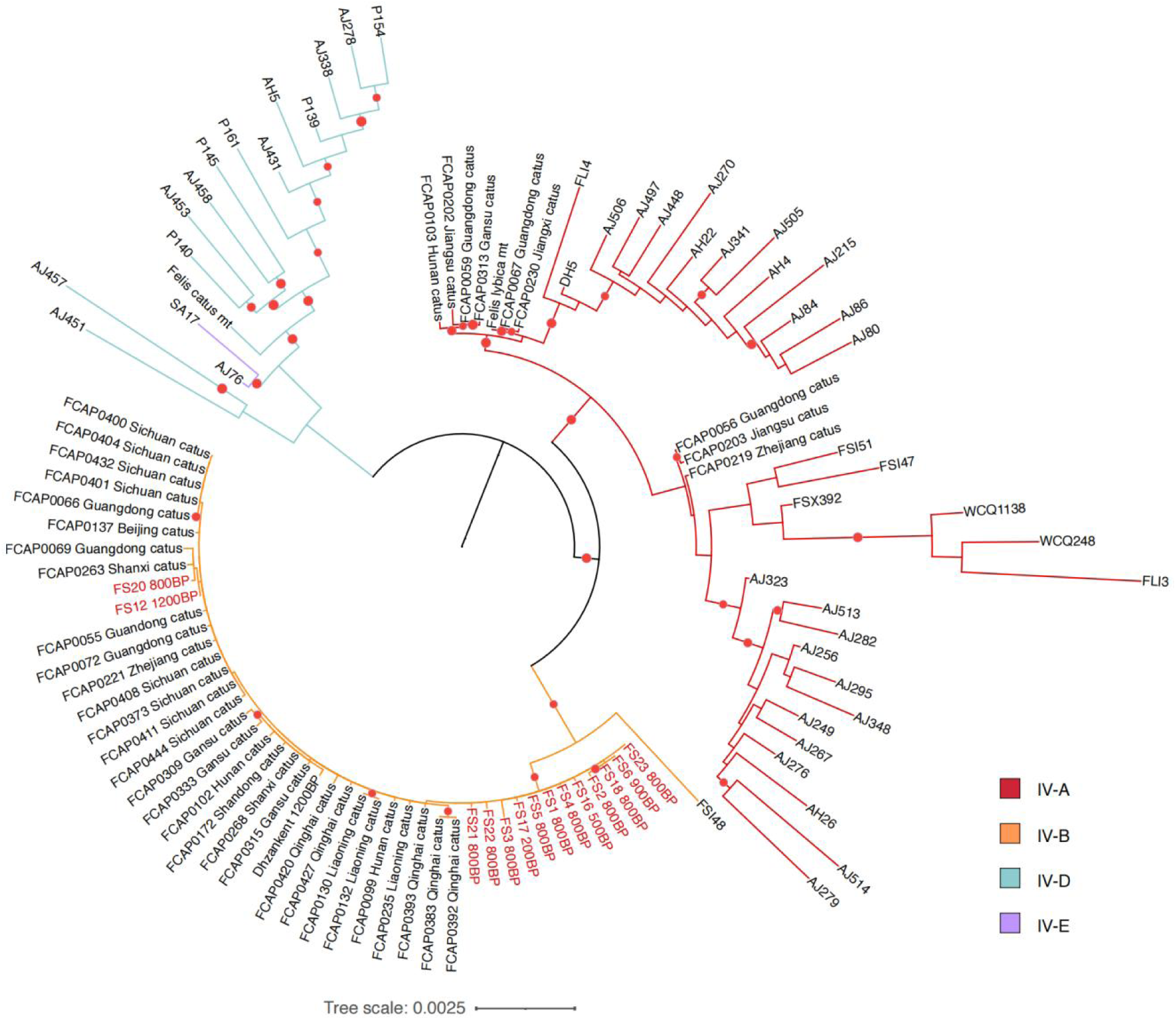
A maximum likelihood (ML) phylogenetic tree based on mitogenomes of ancient and modern domestic cats worldwide, including 14 ancient domestic cats from China. A TN93+I+G model is implemented with 1,000 bootstrap replicates. The missing data rate for Chinese samples is less than 2% each and the sequencing depth is greater than 30×. The missing data rate for European or Central Asian samples is generally less than 10% and the sequencing depth is greater than 10×. Nodes with significant statistical support (≥ 70%) based on 1,000 bootstrap replicates are labeled with red dots.

**Fig. S4.**
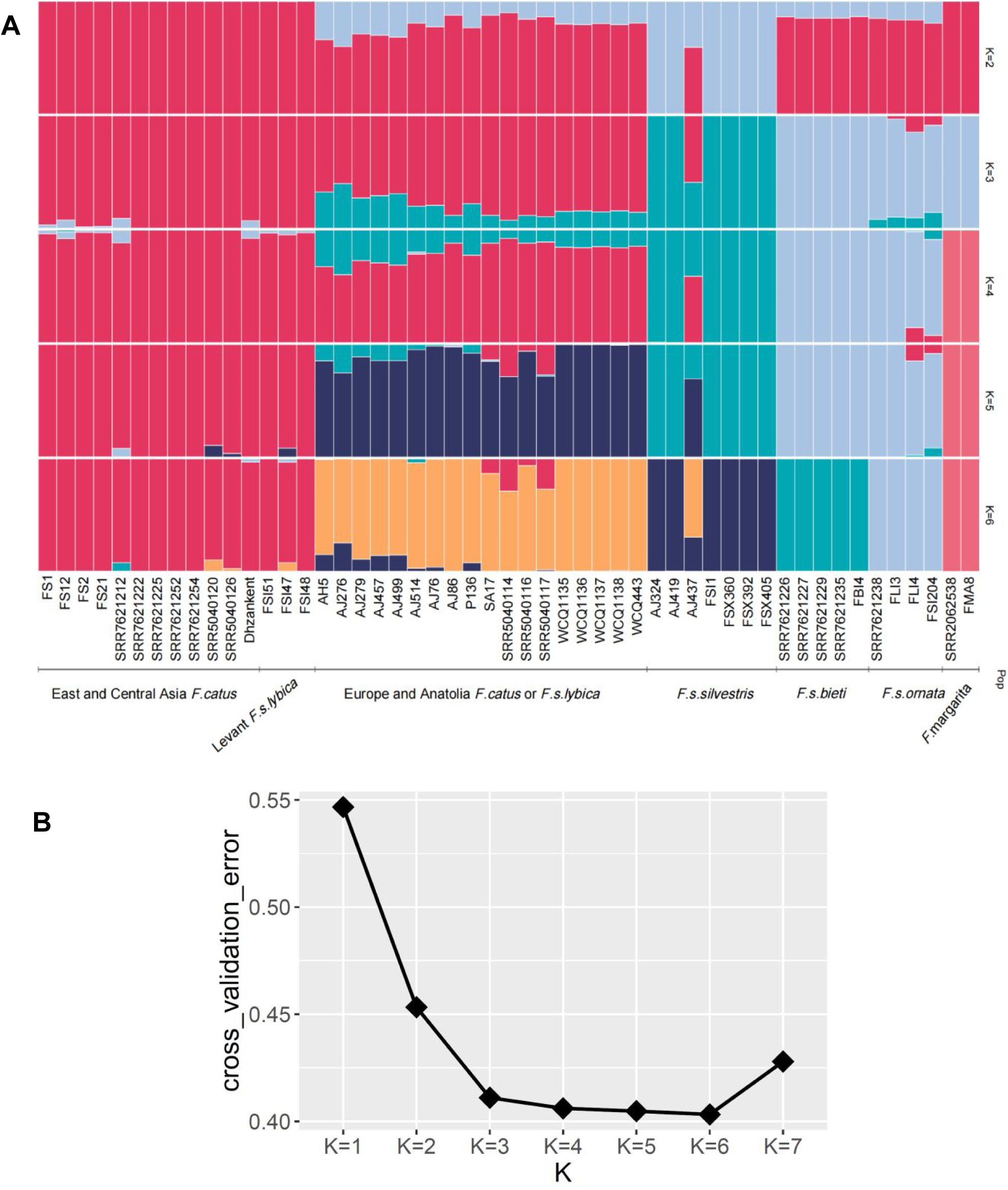
ADMIXTURE analysis for ancient and modern domestic cats and wildcats from Eurasia. **(A)** ADMIXTURE clustering results based on the number of assumed ancestral populations (*K*) from 2 to 6. **(B)** Cross-validation (CV) errors for each number of assumed ancestral populations (*K*) from 1 to 7.

**Fig. S5.**
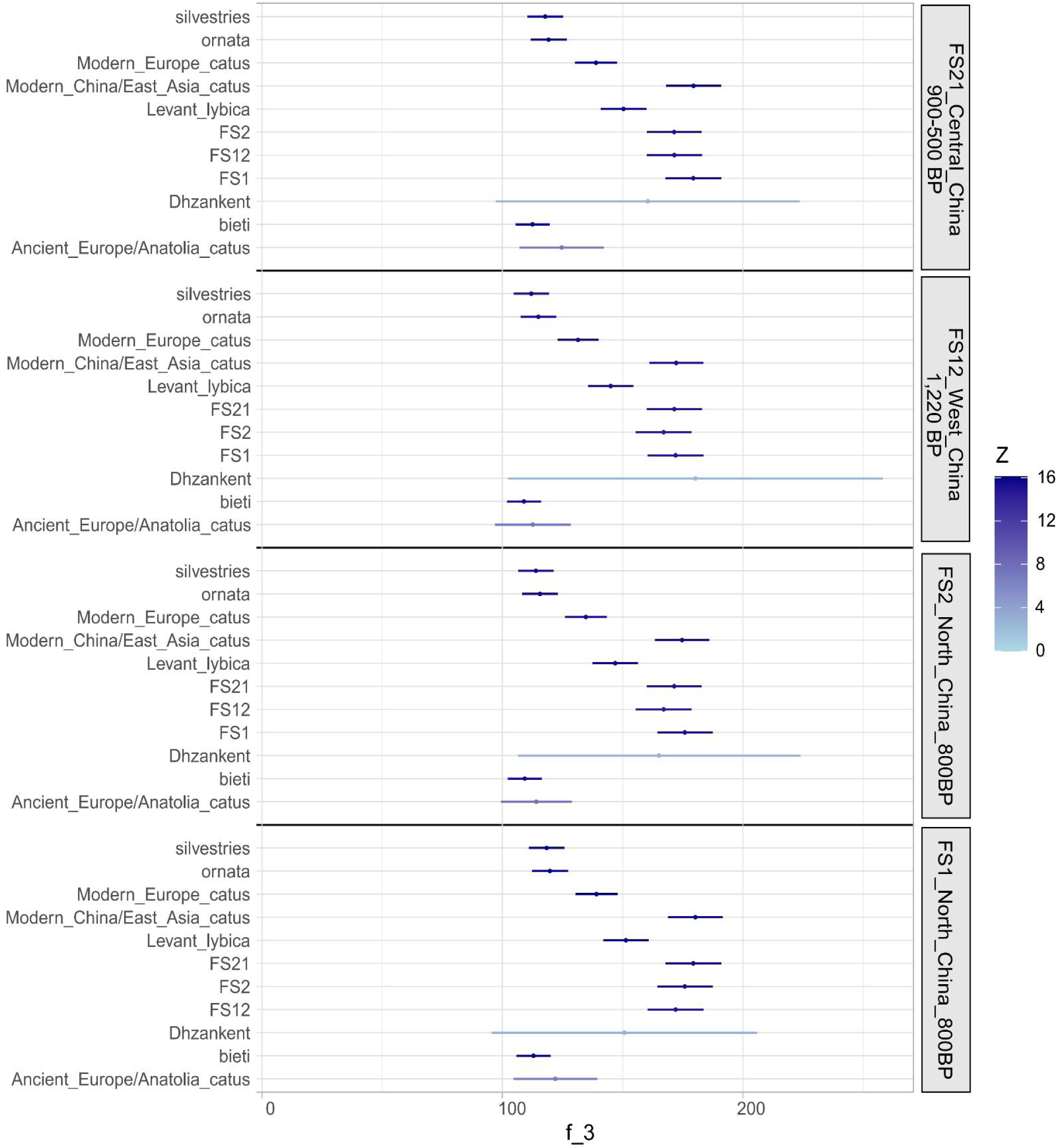
The result of outgroup *f_3_*-statistics illustrates the affinity between ancient domestic cats from China and other domestic cat populations. The error bars in the four stacked blocks, from bottom to top, correspond to the f3 values between FS1/FS2/ FS12/FS21 and other cat populations (labeled on the right). The color gradient, ranging from light to dark blue, shows an increased magnitude of the absolute value of Z. This figure, compared to Fig. 3B, highlights the relationship between ancient domestic cats and modern domestic cats.

**Fig. S6.**
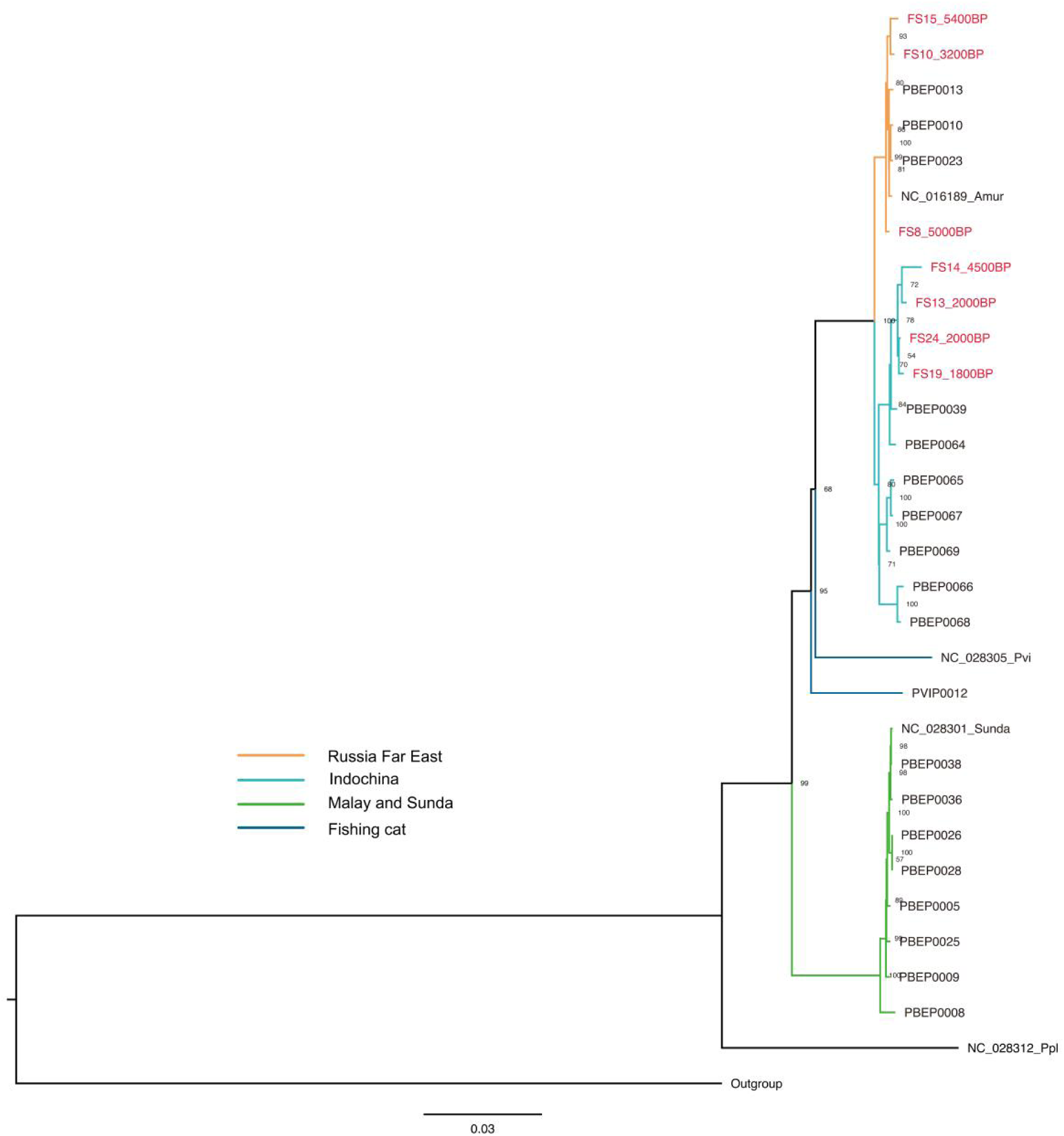
A maximum likelihood (ML) phylogenetic tree based on leopard cat mitochondrial genomes. The tree is reconstructed based on ancient leopard cats from the middle Yellow River Basin of China (n=7) and modern leopard cats from the Russian Far East (n=4), Indochina (n=6), Malay Peninsula (n=6), and Sundaland (n=4). Two fishing cats (*P. viverrinus*) and a flat-headed cat (*P. planiceps*) are included in the analysis. The tiger reference genome (NC_010642.1) is used as an outgroup. The sample names of ancient leopard cats from the Yellow River Basin are marked in red.

**Fig. S7.**
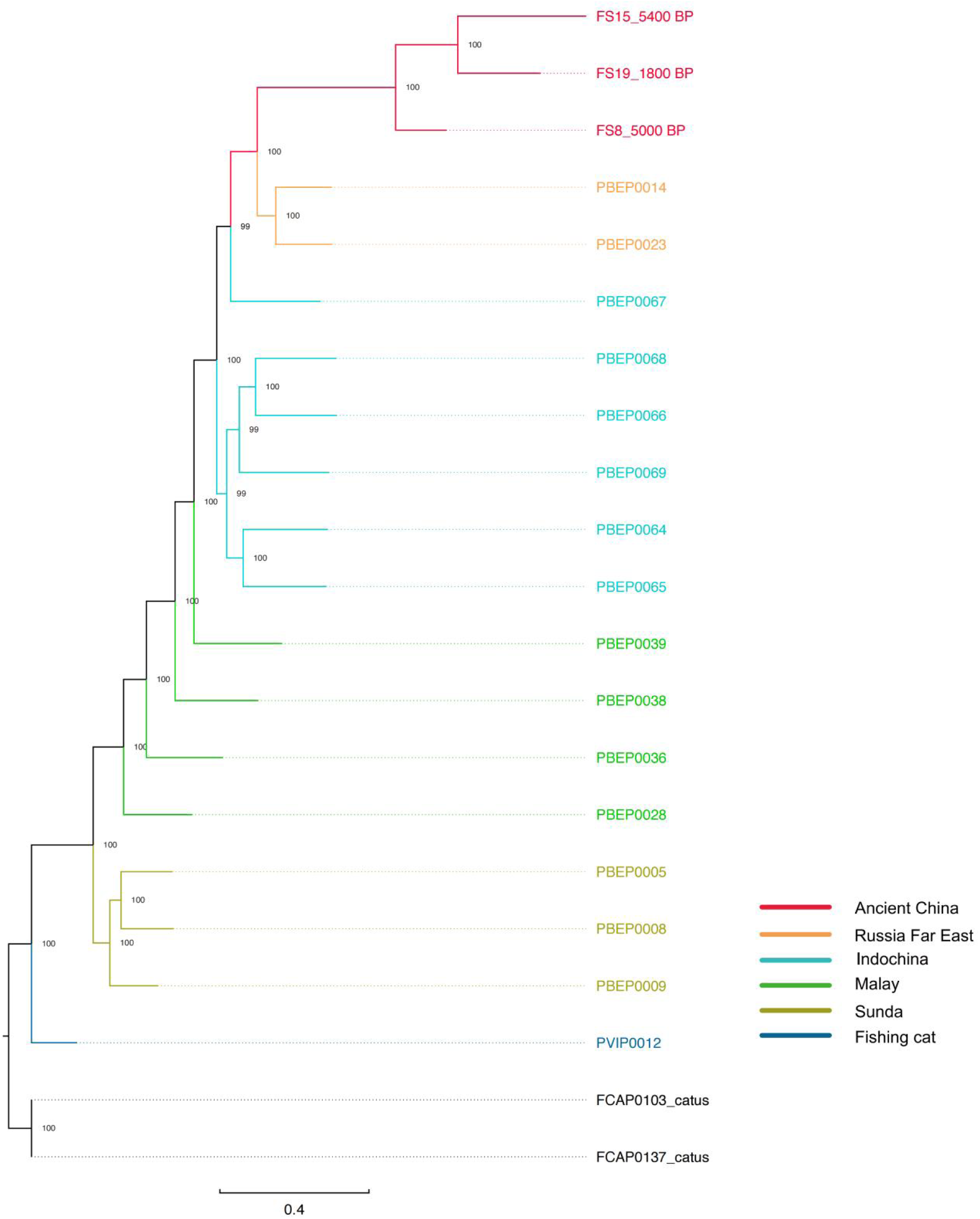
A maximum likelihood (ML) phylogenetic tree based on ancient and modern leopard cats. The tree is reconstructed based on autosomal SNPs from ancient specimens from China (n=3) and modern samples from the Russian Far East (n=2), Indochina (n=6), Malay Peninsula (n=4), and Sunda (n=3). A fishing cat (*P. viverrinus*) is included in the analysis. Two domestic cats are used as outgroups. Numbers at each node represent the statistical support values based on 200 bootstrap replicates.

**Fig. S8.**
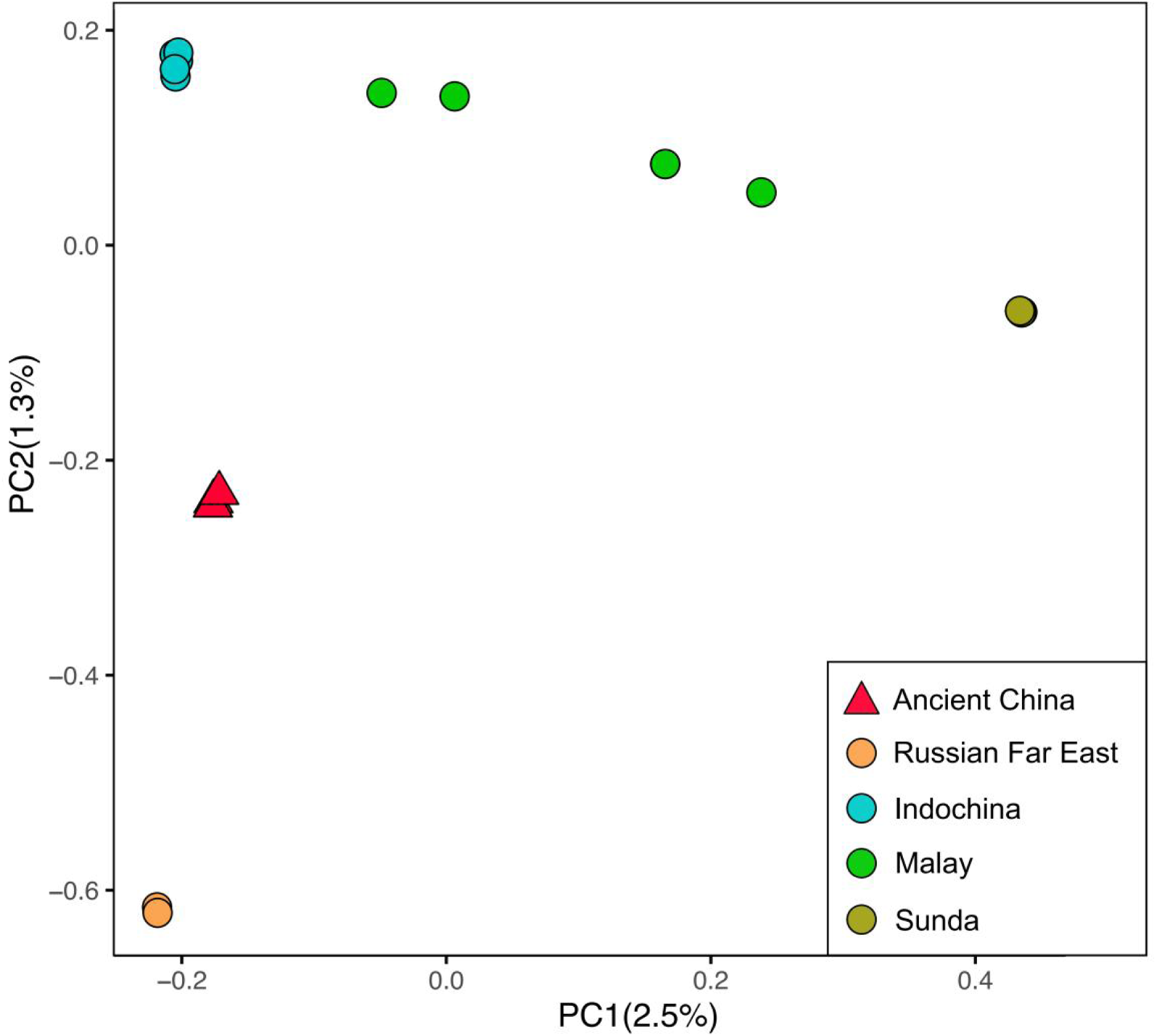
Principal component analysis (PCA) of leopard cats. PCA is first conducted based on autosomal SNPs from high-quality genome data from modern leopard cats. Data from ancient leopard cats are then projected onto it. Three ancient leopard cats from China and 15 modern leopard cats from range wide, including the Russian Far East (n=2), Indochina (n=6), Malay Peninsula (n=4), and Sundaland (n=3), are included in the analysis.

**Fig. S9.**
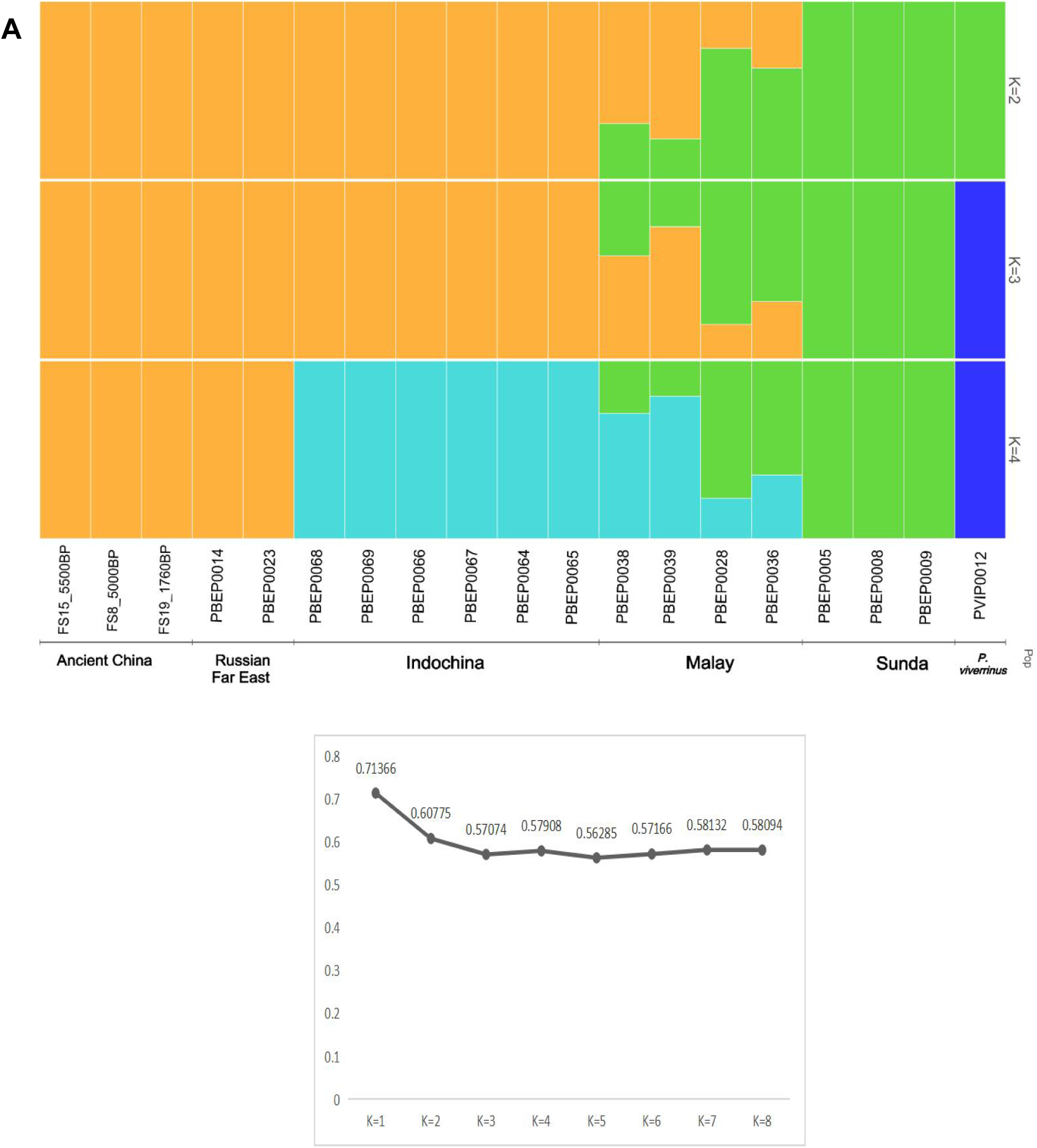
ADMIXTURE analysis of leopard cats. (**A**) Clustering results based on the number of assumed ancestral populations (*K*) from 2 to 4. A total of 18 specimens are used in the analysis including ancient leopard cats from China (n=3) and modern leopard cats from the Russian Far East (n=2), Indochina (n=6), Malay (n=4), and Sundaland (n=3). A fishing cat was used as an outgroup. (**B**) Cross-validation (CV) errors for each number of assumed ancestral populations (*K*) from 1 to 8.

**Fig. S10.**
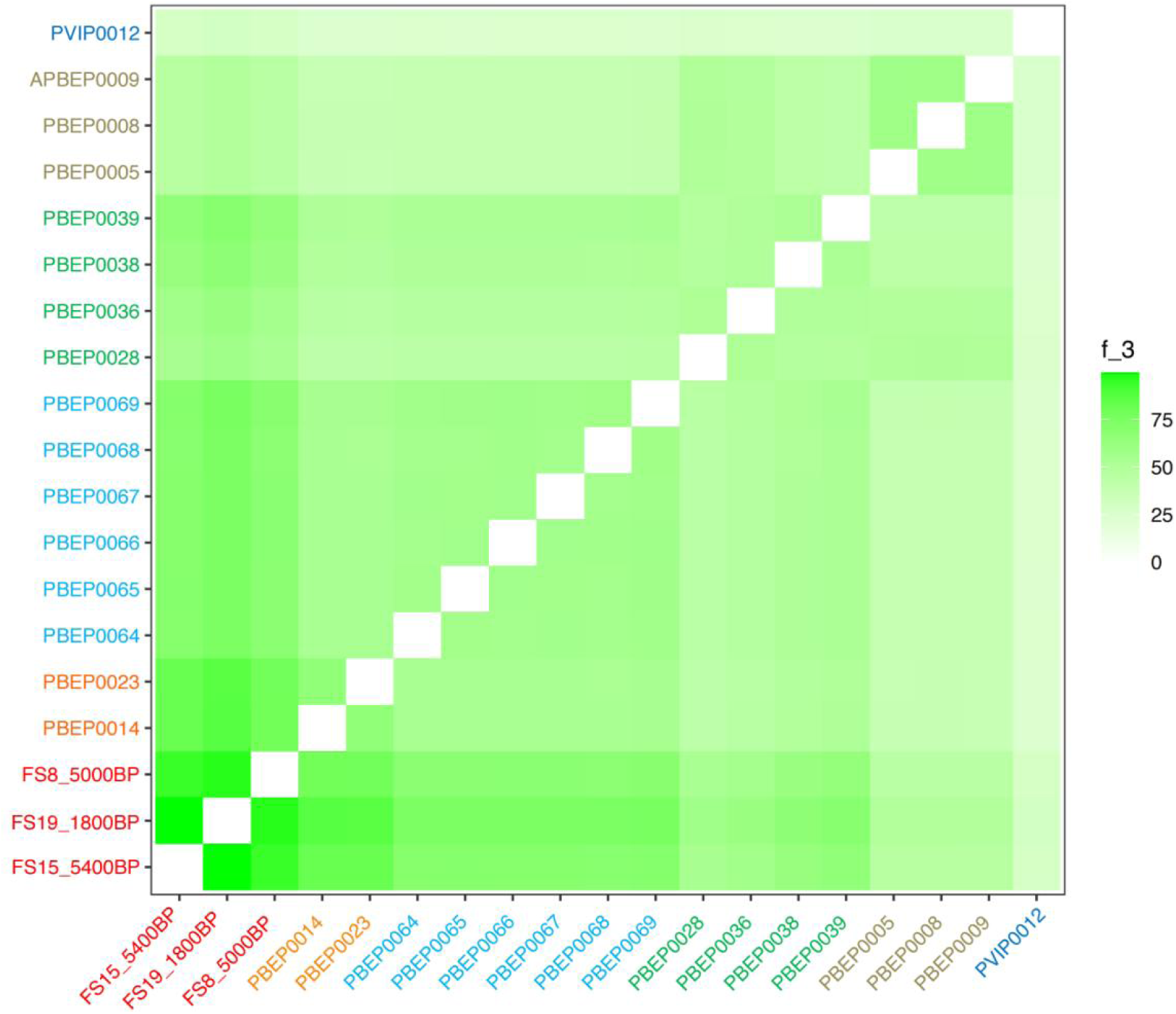
Outgroup *f_3_*-statistics analysis of leopard cats. A heatmap of the outgroup *f_3_*-statistics analysis. Two domestic cats are used as outgroups. The shared drifts of different pairs of individuals of leopard cat or fishing cat are tested. The color gradient, ranging from light to dark green, shows the increased magnitude of f3 values. The sample names from different populations were labeled with different colors. Ancient Chinese leopard cats (n=2) were marked in red; Russian Far East (n=2) in orange; Indochina (n=6) in light blue; Malay (n=4) in green; Sunda (n=3) in brownish yellow; and fishing cat (n=1) in dark blue.

**Fig. S11.**
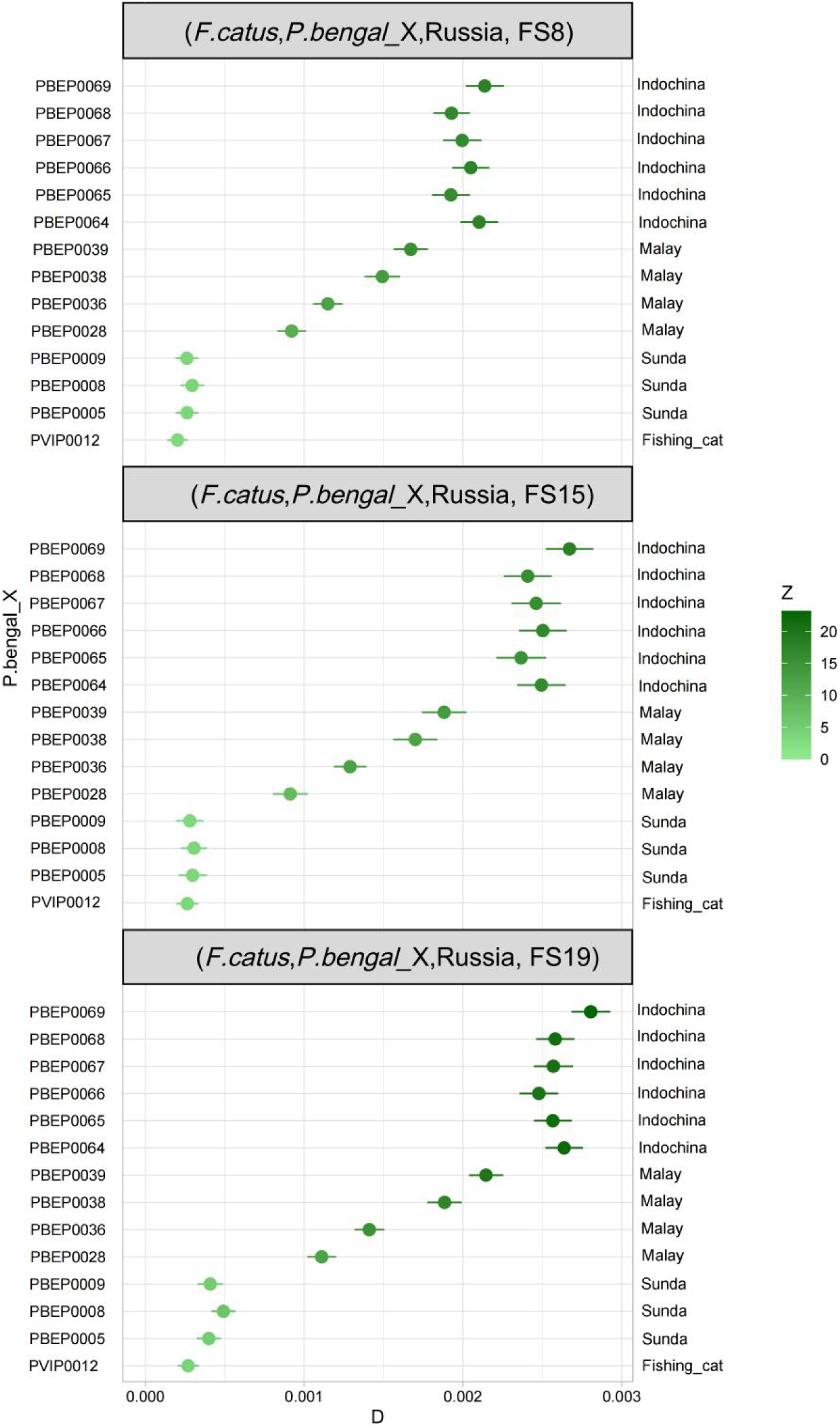
*D-*statistics analysis of leopard cats. The population combinations were set as (Outgroup, *P. bengalensis*_X, Russian Far East, ancient China) (labeled on the top of each block). Two domestic cats were used as outgroups. Leopard cats from Indochina, Malay Peninsula, Sundaland, and a fishing cat are individually placed in P.bengal_X to evaluate the signals of gene flow between the target ancient Chinese leopard cat and a modern leopard cat from P.bengal_X, relative to a Russian Far East leopard cat. The vertical axis shows the ID of leopard cats in P.bengal_X (on the left) and their population assignments (on the right). The horizontal axis shows the *D* scores. The color gradient, ranging from light to dark green, indicates an increased magnitude of |Z| values.

**Fig. S12.**
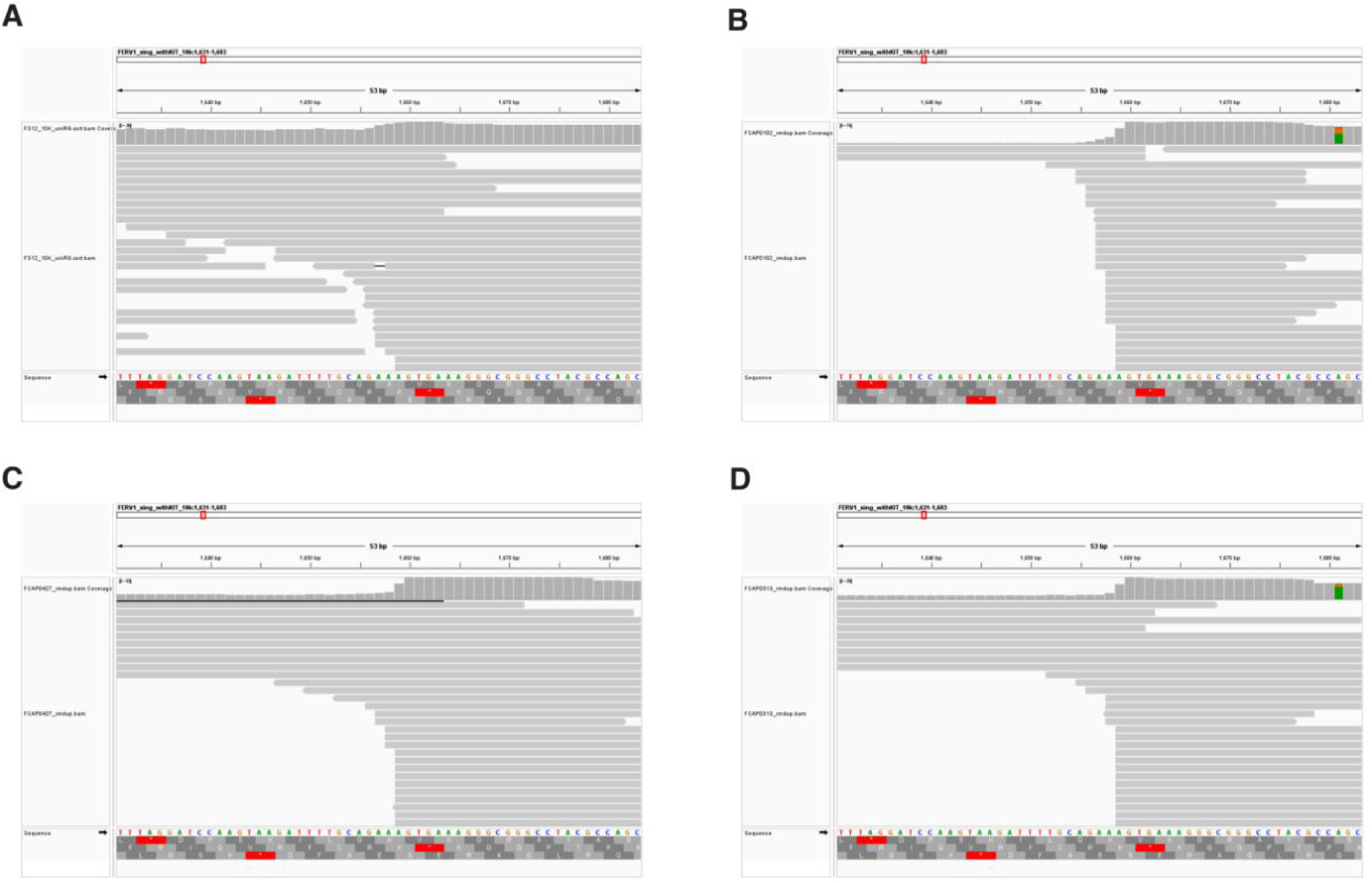
Mapping patterns of the sequencing data to the domestic cat’s *KIT* gene with *FERV1* insertion. **(A)** Alignment of FS12’s *KIT* gene with *FERV1* insertion. **(B)** Alignment of a wild-type domestic cat *KIT* gene without *FERV1* insertion. **(C)** Alignment of a domestic cat homozygous of *FERV1* insertion. **(D)** Alignment of a domestic cat heterozygous of *FERV1* insertion.

## Supplementary Tables

**Table S1.** (A) Information of the ancient samples from this study.

| Sample |  | Geographical |  |  | Endogenous | Num_reads | Length mt | Coverage | Mean_depth | Num_reads | Mean_depth | SNPs_used | Date | MT haplotype | Species |
| --- | --- | --- | --- | --- | --- | --- | --- | --- | --- | --- | --- | --- | --- | --- | --- |
| ID | Site | City and Province | Unit | Country | % | (mt) | (bp) | (mt) % | (mt) x | (Nuc) | (Nuc) x | (Nuc) |  |  |  |
| FS1 | Xitucheng City | Zhangjiakou Hebei | North China | China | 12.5 | 24336 | 16732 | 98.3789 | 86.5136 | 128005822 | 3.22737 | 95470 | 1,115-1,234 CE | IV-B | *F. catus* |
| FS2 | Xitucheng City | Zhangjiakou Hebei | North China | China | 13.82 | 21736 | 16865 | 99.158 | 82.8478 | 101903410 | 2.43784 | 99631 | 1,115-1,234 CE | IV-B | *F. catus* |
| FS3 | Xitucheng City | Zhangjiakou Hebei | North China | China | 11.4 | 11063 | 16473 | 96.8487 | 34.7035 | N/A | N/A | N/A | 1,115-1,234 CE | IV-B | *F. catus* |
| FS4 | Xitucheng City | Zhangjiakou Hebei | North China | China | 10.69 | 15936 | 16570 | 97.419 | 59.8712 | N/A | N/A | N/A | 1,115-1,234 CE | IV-B | *F. catus* |
| FS5 | Xitucheng City | Zhangjiakou Hebei | North China | China | 11.2 | 29916 | 16769 | 98.589 | 39.2182 | N/A | N/A | N/A | 1,115-1,234 CE | IV-B | *F. catus* |
| FS6 | Xinxiang Song Tomb | Xinxiang Henan | Central China | China | 0.34 | 15982 | 16961 | 99.7178 | 112.491 | N/A | N/A | N/A | 960-1,127 CE | IV-B | *F. catus* |
| FS7 | Huoshiliang | Yulin Shaanxi | West China | China | 0.91 | 17321 | 16730 | 99.53 | 16.9127 | N/A | N/A | N/A | 1,800 BCE | *Felis ornata* | *F. ornata* |
| FS8 | Wuzhuangguoliang | Jingbian Shaanxi | West China | China | 8.76 | 9870 | 16416 | 98.2935 | 47.3244 | 153959749 | 5.96269 | 17194519 | 3,000-221 BCE | PBE-RFE | *P. bengalensis* |
| FS10 | Laoniupo | Xi'an Shaanxi | Central China | China | 0.45 | 24215 | 16338 | 97.8265 | 11.5208 | N/A | N/A | N/A | 1,200 BCE | PBE-RFE | *P. bengalensis* |
| FS12* | Tongwan City | Jingbian Shaanxi | West China | China | 13.21 | 23207 | 16742 | 98.4377 | 78.8921 | 111504420 | 3.52959 | 107849 | 706-883 cal CE | IV-B | *F. catus* |
| FS13 | Yangling | Xianyang Shaanxi | Central China | China | 0.06 | 4117 | 15792 | 94.5572 | 26.2551 | N/A | N/A | N/A | 168 - 48 cal BCE | PBE-Indochina | *P. bengalensis* |
| FS14 | Xinjie | Lantian Shaanxi | Central China | China | 0.06 | 12377 | 15584 | 93.3118 | 20.1013 | N/A | N/A | N/A | ~3,500-1,800 BCE | PBE-Indochina | *P. bengalensis* |
| FS15 | Quanhucun | Weinan Shaanxi | Central China | China | 2.3 | 2826 | 16114 | 96.4852 | 15.6682 | 162299570 | 3.32325 | 14322059 | 4,000-2,700 BCE | PBE-RFE | *P. bengalensis* |
| FS16 | Zhengzhoushang City | Zhengzhou Henan | Central China | China | 4.04 | 6512 | 16991 | 99.8942 | 207.187 | N/A | N/A | N/A | 1,413 - 1,480 cal CE | IV-B | *F. catus* |
| FS17 | Shaliangzi | Hohhot Inner Mongolia | North China | China | 17.89 | 5274 | 16871 | 99.1887 | 45.9693 | N/A | N/A | N/A | 1,797 - 1,944 cal CE | IV-B | *F. catus* |
| FS18 | Puyang Song Tomb | Puyang Henan | Central China | China | 0.05 | 4969 | 16975 | 99.8001 | 145.439 | N/A | N/A | N/A | 618-970 CE | IV-B | *F. catus* |
| FS19 | Zheng and Han City | Xinzheng Henan | Central China | China | 6.48 | 27643 | 16518 | 98.9043 | 51.2599 | 175356056 | 4.70431 | 15747266 | 25-220 CE | PBE-Indochina | *P. bengalensis* |
| FS20 | Zheng and Han City | Xinzheng Henan | Central China | China | 0.05 | 10271 | 16694 | 98.148 | 99.1598 | N/A | N/A | N/A | 960-1,368 CE | IV-B | *F. catus* |
| FS21 | Zheng and Han City | Xinzheng Henan | Central China | China | 10.24 | 19562 | 16708 | 98.237 | 77.6107 | 115273634 | 3.23206 | 95137 | 960-1,368 CE | IV-B | *F. catus* |
| FS22 | Zheng and Han City | Xinzheng Henan | Central China | China | 0.1 | 25761 | 16593 | 97.5542 | 59.5793 | N/A | N/A | N/A | 960-1,368 CE | IV-B | *F. catus* |
| FS23 | Zheng and Han City | Xinzheng Henan | Central China | China | 3.42 | 13341 | 17009 | 100 | 304.291 | N/A | N/A | N/A | 960-1,368 CE | IV-B | *F. catus* |
| FS24 | Han Chang'an City | Xi'an Shaanxi | Central China | China | 2.7 | 20836 | 16617 | 99.497 | 90.8493 | N/A | N/A | N/A | 44 cal BCE-76 cal CE | PBE-Indochina | *P. bengalensis* |

\* We conducted additional deep sequencing on the sample to reach a final genome sequencing depth of 15.97x.

**Table S1.**
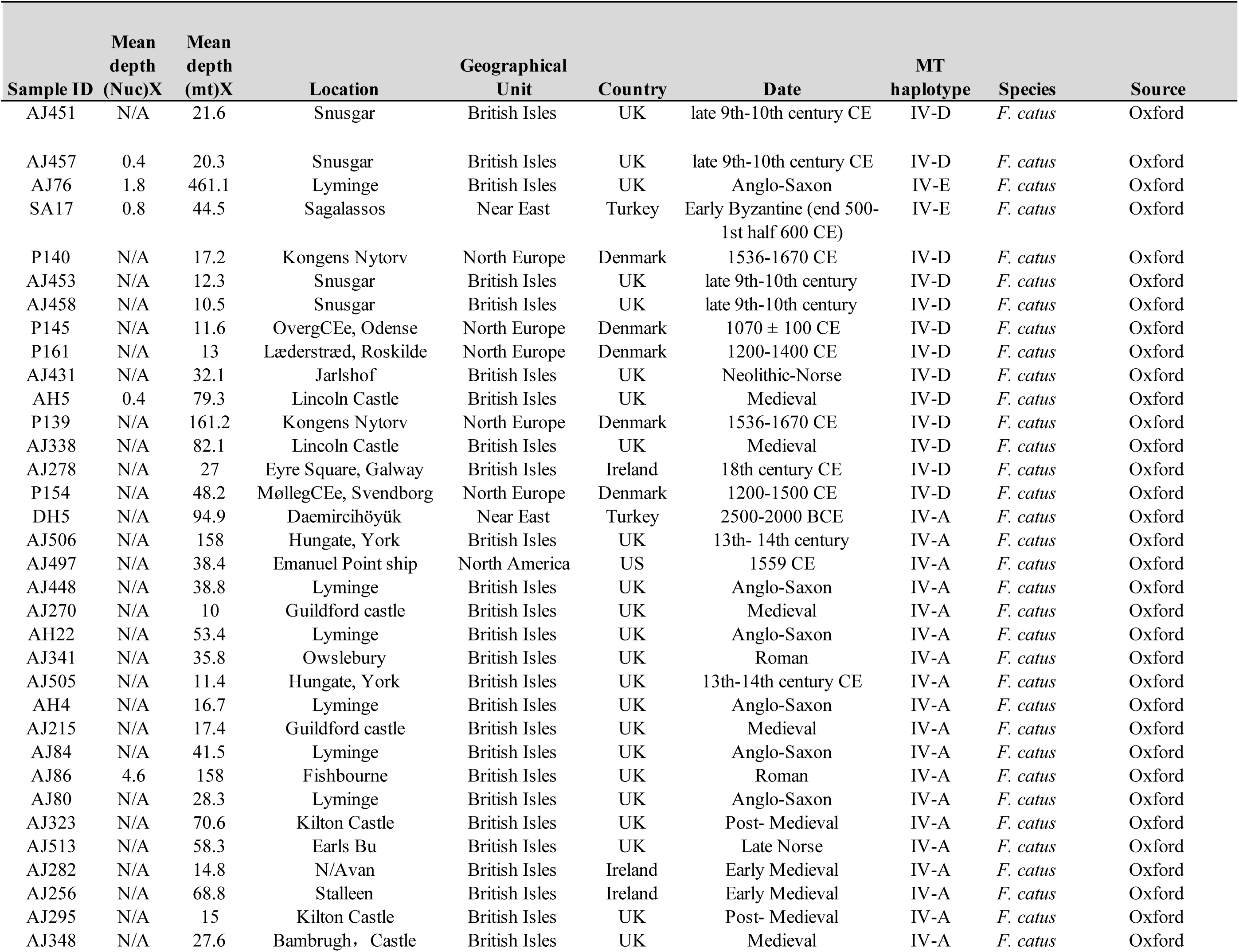
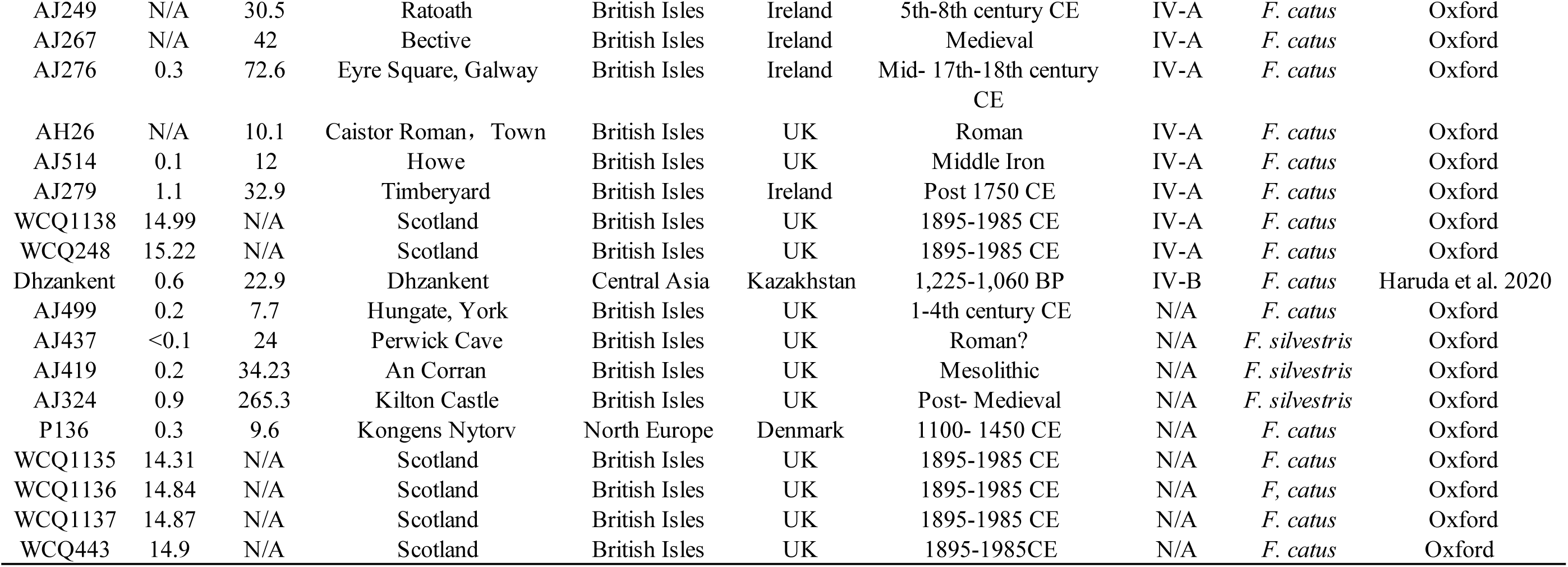
(B) Information of ancient samples from other studies.

**Table S1.** (C) Information of modern Felis samples from other resources.

| Sample ID | Source | Mean depth (Nuc) X | Mean depth (mt) X | Location | Geographical Unit | Country | Date | Species | MT haplotype |
| --- | --- | --- | --- | --- | --- | --- | --- | --- | --- |
| FCAP0103 | SRR7621222 | 14.65 | 1042.19 | Hunan | South China | China | modern | *F. catus* | IV-A |
| FCAP0202 | SRR7621253 | N/A | 2305.22 | Jiangsu | East Coast of China | China | modern | *F. catus* | IV-A |
| FCAP0059 | SRR7621259 | N/A | 355 | Guangdong | South Coast of China | China | modern | *F. catus* | IV-A |
| FCAP0067 | SRR7621233 | N/A | 803.31 | Guangdong | South Coast of China | China | modern | *F. catus* | IV-A |
| FCAP0230 | SRR7621256 | N/A | 823.40 | Jiangxi | South China | China | modern | *F. catus* | IV-A |
| FLI4 | SRR15116525 | 10.1 | N/A | Tajikistan | Central Asia | Tajikistan | modern | *F. ornata* | IV-A |
| FCAP0056 | SRR7621254 | 10.52 | 283.96 | Guangdong | South Coast of China | China | modern | *F. catus* | IV-A |
| FCAP0203 | SRR7621252 | 9.48 | 830.91 | Jiangsu | East Coast of China | China | modern | *F. catus* | IV-A |
| FCAP0219 | SRR7621250 | N/A | 898.11 | Zhejiang | East Coast of China | China | modern | *F. catus* | IV-A |
| FCAP0128 | SRR7621225 | 12.63 | 2231.54 | Hebei | North China | China | modern | *F. catus* | N/A |
| FSI51 | ERR10494336-ERR10494341 | 21.21 | N/A | Israel | the Levant | Israel | modern | *F. lybica* | IV-A |
| FSI47 | ERR10494332 | 11.17 | N/A | Israel | the Levant | Israel | modern | *F. lybica* | IV-A |
| FSX392 | ERR10494345 | 12.09 | N/A | Scotland | British Isles | Scotland | modern | *F. silvestris* | IV-A |
| FLI3 | SRR15116526 | 9.04 | N/A | Tajikistan | Central Asia | Tajikistan | modern | *F. ornata* | IV-A |
| FSI48 | ERR10494333-ERR10494335 | 18.76 | N/A | Israel | the Levant | Israel | modern | *F. lybica* | IV-B |
| FCAP0400 | SRR7621207 | N/A | 729.82 | Sichuan | Southwest China | China | modern | *F. catus* | IV-B |
| FCAP0404 | SRR7621243 | N/A | 704.08 | Sichuan | Southwest China | China | modern | *F. catus* | IV-B |
| FCAP0432 | SRR7621244 | N/A | 861.71 | Sichuan | Southwest China | China | modern | *F. catus* | IV-B |
| FCAP0401 | SRR7621206 | N/A | 510.36 | Sichuan | Southwest China | China | modern | *F. catus* | IV-B |
| FCAP0066 | SRR7621232 | N/A | 325.04 | Guangdong | South Coast of China | China | modern | *F. catus* | IV-B |
| FCAP0137 | SRR7621218 | N/A | 801.72 | Beijing | North China | China | modern | *F. catus* | IV-B |
| FCAP0069 | SRR7621230 | N/A | 436.44 | Guangdong | South Coast of China | China | modern | *F. catus* | IV-B |
| FCAP0263 | SRR7621223 | N/A | 790.72 | Shannxi | North China | China | modern | *F. catus* | IV-B |
| FCAP0055 | SRR7621255 | N/A | 376.78 | Guangdong | South Coast of China | China | modern | *F. catus* | IV-B |
| FCAP0072 | provided by PKU | N/A | 756.09 | Guangdong | South Coast of China | China | modern | *F. catus* | IV-B |
| FCAP0221 | provided by PKU | N/A | N/A | Zhejiang | East Coast of China | China | modern | *F. catus* | IV-B |
| FCAP0408 | SRR7621242 | N/A | 585.48 | Sichuan | Southwest China | China | modern | *F. catus* | IV-B |
| FCAP0373 | SRR7621246 | N/A | 796.02 | Sichuan | Southwest China | China | modern | *F. catus* | IV-B |
| FCAP0411 | SRR7621245 | N/A | 620.53 | Sichuan | Southwest China | China | modern | *F. catus* | IV-B |
| FCAP0444 | SRR7621247 | N/A | 1125.63 | Sichuan | Southwest China | China | modern | *F. catus* | IV-B |
| FCAP0309 | SRR7621211 | N/A | 1053.49 | Gansu | West China | China | modern | *F. catus* | IV-B |
| FCAP0333 | SRR7621248 | N/A | 681.41 | Gansu | West China | China | modern | *F. catus* | IV-B |
| FCAP0102 | SRR7621240 | N/A | 669.54 | Hunan | South China | China | modern | *F. catus* | IV-B |
| FCAP0172 | SRR7621217 | N/A | 1040.97 | Shandong | Central China | China | modern | *F. catus* | IV-B |
| FCAP0268 | SRR7621224 | N/A | 1091.05 | Shannxi | North China | China | modern | *F. catus* | IV-B |
| FCAP0315 | SRR7621209 | N/A | 1084.61 | Gansu | West China | China | modern | *F. catus* | IV-B |
| FCAP0420 | SRR7621208 | N/A | 615.50 | Qinghai | West China | China | modern | *F. catus* | IV-B |
| FCAP0427 | SRR7621215 | N/A | 1132.00 | Qinghai | West China | China | modern | *F. catus* | IV-B |
| FCAP0130 | SRR7621219 | N/A | 600.37 | Liaoning | North China | China | modern | *F. catus* | IV-B |

| Sample ID | Source | Mean depth<br>(Nuc) X | Mean depth<br>(mt) X | Location | Geographical Unit | Country | Date | Species | MT<br>haplotype |
| --- | --- | --- | --- | --- | --- | --- | --- | --- | --- |
| FCAP0132 | SRR7621220 | N/A | 984.46 | Liaoning | North China | China | modern | *F. catus* | IV-B |
| FCAP0099 | SRR7621241 | N/A | 408.39 | Hunan | South China | China | modern | *F. catus* | IV-B |
| FCAP0235 | SRR7621216 | N/A | 933.45 | Liaoning | North China | China | modern | *F. catus* | IV-B |
| FCAP0393 | SRR7621212 | 15.46 | 832.63 | Qinghai | West China | China | modern | *F. catus* | IV-B |
| FCAP0383 | SRR7621214 | N/A | 539.14 | Qinghai | West China | China | modern | *F. catus* | IV-B |
| FCAP0392 | SRR7621213 | N/A | 1589.83 | Qinghai | West China | China | modern | *F. catus* | IV-B |
| SRR5040120 | SRR5040120 | 14.67 | N/A | Thailand | East Asia | Thailand | modern | *F. catus* | N/A |
| SRR5040126 | SRR5040126 | 14.58 | N/A | South Korea | East Asia | South Korea | modern | *F. catus* | N/A |
| SRR5040114 | SRR5040114 | 9.95 | N/A | Portugal | Mainland Europe | Portugal | modern | *F. catus* | N/A |
| SRR5040114 | SRR5040114 | 9.95 | N/A | Portugal | Mainland Europe | Portugal | modern | *F. catus* | N/A |
| SRR5040117 | SRR5040117 | 13.04 | N/A | Italy | Mainland Europe | Italy | modern | *F. catus* | N/A |
| FSI1 | SRR8401848 | 16.78 | N/A | Azerbaijan | Central Asia | Azerbaijan | modern | *F. silvestris* | N/A |
| FSX360 | ERR10494342-ERR10494344 | 17.35 | N/A | Portugal | Mainland Europe | Portugal | modern | *F. silvestris* | N/A |
| FSX405 | ERR10494347-ERR10494349 | 15.3 | N/A | Scotland | British Isles | Scotland | modern | *F. silvestris* | N/A |
| FBIP0004 | SRR7621226 | 13.20 | 911.04 | Gansu | West China | China | modern | *F. bieti* | N/A |
| FBIP0008 | SRR7621227 | 11.48 | 929.99 | Sichuan | Southwest China | China | modern | *F. bieti* | N/A |
| FBIP0003 | SRR7621229 | 11.90 | 871.06 | Qinghai | West China | China | modern | *F. bieti* | N/A |
| FSIP0006 | SRR7621238 | 11.08 | 4789.78 | Xinjiang | West China | China | modern | *F. ornata* | N/A |
| FBIP0026 | SRR7621235 | 11.87 | 642.33 | Gansu | West China | China | modern | *F. bieti* | N/A |
| FSI204 | ERR10494331 | 10.74 | N/A | Kazakhstan | Central Asia | Kazakhstan | modern | *F. ornata* | N/A |
| SRR2062538 | SRR2062538 | N/A | N/A | N/A | N/A | N/A | modern | *F. margarita* | N/A |
| FMA8 | ERR10494328-ERR10494330 | 22.74 | N/A | N/A | N/A | N/A | modern | *F. margarita* | N/A |

**Table S1.** (D) Information of Leopard cat and fishing cat samples from other studies.

| **ID** | **Source** | **location** | **Mean_depth**<br>**(Nuc)X** | **Mean_depth**<br>**(mt)X** | **Date** | **MT_haplotype** |
| --- | --- | --- | --- | --- | --- | --- |
| PBEP0014 | SRR4426166 | Russian Far East | 12.17 | 1719 | modern | PBE-RFE |
| PBEP0023 | SRR4426167 | Russian Far East | 11.1209 | 621 | modern | PBE-RFE |
| PBEP0010 | SRR4426164 | Russian Far East | N/A | 747 | modern | PBE-RFE |
| PVIP0012 | SRR4426169 | N/A | 13.9554 | 612 | modern | PVI |
| PBEP0005 | SRR4426170 | Sunda | 11.3489 | 530 | modern | PBE-Sunda |
| PBEP0008 | SRR4426171 | Sunda | 11.7308 | 1415 | modern | PBE-Sunda |
| PBEP0009 | SRR4426172 | Sunda | 11.0208 | 1094 | modern | PBE-Sunda |
| PBEP0068 | SRR4426173 | Indochina | 13.1926 | 1800 | modern | PBE-Indochina |
| PBEP0069 | SRR4426174 | Indochina | 11.9837 | 948 | modern | PBE-Indochina |
| PBEP0066 | SRR4426175 | Indochina | 11.4505 | 579 | modern | PBE-Indochina |
| PBEP0067 | SRR4426176 | Indochina | 11.6046 | 1216 | modern | PBE-Indochina |
| PBEP0064 | SRR4426177 | Indochina | 11.3286 | 911 | modern | PBE-Indochina |
| PBEP0065 | SRR4426178 | Indochina | 11.4206 | 1302 | modern | PBE-Indochina |
| PBEP0038 | SRR4426179 | Malay | 13.0935 | 1320 | modern | PBE-Sunda |
| PBEP0039 | SRR4426180 | Malay | 12.8588 | 840 | modern | PBE-Indochina |
| PBEP0028 | SRR4426181 | Malay | 12.7609 | 1209 | modern | PBE-Sunda |
| PBEP0036 | SRR4426182 | Malay | 11.7445 | 2397 | modern | PBE-Sunda |
| PBEP0026 | SRR4426163 | Malay | N/A | 1324 | modern | PBE-Sunda |
| PBEP0025 | SRR4426162 | Malay | N/A | 1290 | modern | PBE-Sunda |
| NC_016189.1 | N/A | N/A | N/A | N/A | modern | PBE-RFE |
| NC_028301.1 | N/A | N/A | N/A | N/A | modern | PBE-Sunda |

**Table S2.** Radiocarbon dating of small cat specimens from archeological sites of China.

| ID | Site | Location | Context date | Uncalibrated date (BP) | Date cal. BCE/CE (95.4%, overall range IntCal 20) |
| --- | --- | --- | --- | --- | --- |
| FS12 | Tongwan City | Western China | Song Dynasty | 1,220+/-20 | 706-728 cal CE (7.5%) (1,244 -1,222 cal BP) 772 - 883 cal CE (88.0%) (1,178 -1,067 cal BP) |
| FS13 | Yangling Tomb | Central China | Han Dynasty | 2,095+/-20 | 168 - 48 cal BCE (95.4%) (2,118 - 1,998 cal BP ) |
| FS16 | Zhengzhou Shang City | Central China | Tang Dynasty | 450+/-30 | 1,413 - 1,480 cal CE (95.4%) (537 - 470 cal BP) |
| FS17 | Shanliangzi | Northern China | Han Dynasty | 140+/-30 | 1,797 - 1,944 cal CE (153 - 6 cal BP) (58.2%); 1,671 - 1,779 cal CE (37.2%) (279 - 171 cal BP) |
| FS19 | Zheng and Han City | Central China | Han Dynasty | 1,760+/-30 | 200 - 380 cal CE (95.4%) (1,750 - 1,570 cal BP) |
| FS24 | Han Chang'an City | Central China | Han Dynasty | 1,995+/-20 | 44 cal BCE - 76 cal CE (95.4%) (1,994 -1,874 cal BP) |

**Notes:** Six samples were submitted for accelerator mass spectrometry (AMS) dating. Three samples (FS16, FS17, and FS19) were dated at Beta Analytic in the United States, while the remaining samples (FS12, FS13, and FS24) were processed at the School of Archaeology, Peking University, China. All radiocarbon dates reported were calibrated using the INTCAL20 calibration curve (Reimer et al., 2020).

**Table S3.** Pairwise outgroup f3-statistics of ancient and modern leopard cats (Data of Fig. S10)

| Pop1 | Pop2 | Outgroup | f 3 | std.err | Z | SNPs |
| --- | --- | --- | --- | --- | --- | --- |
| FS15 | FS15 | *Felis catus* | 0 | -1 | 0 | -1 |
| FS15 | FS8 | *Felis catus* | 94.479252 | 3.161095 | 29.888 | 703660 |
| FS15 | FS19 | *Felis catus* | 99.536262 | 3.275273 | 30.39 | 642087 |
| FS15 | PBEP0014 | *Felis catus* | 82.320955 | 2.499533 | 32.935 | 785997 |
| FS15 | PBEP0023 | *Felis catus* | 82.094914 | 2.511656 | 32.686 | 780733 |
| FS15 | PVIP0012 | *Felis catus* | 28.737047 | 0.917571 | 31.319 | 700068 |
| FS15 | PBEP0005 | *Felis catus* | 46.123892 | 1.4261 | 32.343 | 858528 |
| FS15 | PBEP0008 | *Felis catus* | 46.164718 | 1.428339 | 32.321 | 857682 |
| FS15 | PBEP0009 | *Felis catus* | 45.917531 | 1.403135 | 32.725 | 858237 |
| FS15 | PBEP0068 | *Felis catus* | 69.950411 | 2.10229 | 33.273 | 894240 |
| FS15 | PBEP0069 | *Felis catus* | 71.417436 | 2.211924 | 32.287 | 882128 |
| FS15 | PBEP0066 | *Felis catus* | 70.576964 | 2.153776 | 32.769 | 884207 |
| FS15 | PBEP0067 | *Felis catus* | 70.503829 | 2.15352 | 32.739 | 851996 |
| FS15 | PBEP0064 | *Felis catus* | 70.628207 | 2.143967 | 32.943 | 885700 |
| FS15 | PBEP0065 | *Felis catus* | 70.682337 | 2.159481 | 32.731 | 887683 |
| FS15 | PBEP0038 | *Felis catus* | 62.92979 | 1.919277 | 32.788 | 915250 |
| FS15 | PBEP0039 | *Felis catus* | 65.392363 | 1.992234 | 32.824 | 907600 |
| FS15 | PBEP0028 | *Felis catus* | 54.652416 | 1.671718 | 32.692 | 911959 |
| FS15 | PBEP0036 | *Felis catus* | 57.967153 | 1.790102 | 32.382 | 907314 |
| FS8 | FS8 | *Felis catus* | 0 | -1 | 0 | -1 |
| FS8 | FS15 | *Felis catus* | 94.479252 | 3.161095 | 29.888 | 703660 |
| FS8 | FS19 | *Felis catus* | 96.856843 | 3.252197 | 29.782 | 760198 |
| FS8 | PBEP0014 | *Felis catus* | 79.309033 | 2.459222 | 32.25 | 974613 |
| FS8 | PBEP0023 | *Felis catus* | 78.955336 | 2.462822 | 32.059 | 969790 |
| FS8 | PVIP0012 | *Felis catus* | 28.183928 | 0.906908 | 31.077 | 888823 |
| FS8 | PBEP0005 | *Felis catus* | 45.603351 | 1.436178 | 31.753 | 1074427 |
| FS8 | PBEP0008 | *Felis catus* | 45.538826 | 1.439044 | 31.645 | 1072055 |
| FS8 | PBEP0009 | *Felis catus* | 45.358966 | 1.405665 | 32.269 | 1073212 |
| FS8 | PBEP0068 | *Felis catus* | 68.206189 | 2.100244 | 32.475 | 1107559 |
| FS8 | PBEP0069 | *Felis catus* | 69.39284 | 2.205899 | 31.458 | 1090504 |
| FS8 | PBEP0066 | *Felis catus* | 68.65733 | 2.146608 | 31.984 | 1094816 |
| FS8 | PBEP0067 | *Felis catus* | 68.672333 | 2.137885 | 32.122 | 1055827 |
| FS8 | PBEP0064 | *Felis catus* | 68.828178 | 2.138988 | 32.178 | 1095339 |
| FS8 | PBEP0065 | *Felis catus* | 68.826358 | 2.144858 | 32.089 | 1098540 |
| FS8 | PBEP0038 | *Felis catus* | 61.557468 | 1.923598 | 32.001 | 1135282 |
| FS8 | PBEP0039 | *Felis catus* | 64.069036 | 1.998685 | 32.056 | 1123996 |
| FS8 | PBEP0028 | *Felis catus* | 53.800577 | 1.676275 | 32.095 | 1134106 |
| FS8 | PBEP0036 | *Felis catus* | 56.795691 | 1.798561 | 31.578 | 1126856 |
| FS19 | FS19 | *Felis catus* | 0 | -1 | 0 | -1 |
| FS19 | FS15 | *Felis catus* | 99.536262 | 3.275273 | 30.39 | 642087 |
| FS19 | FS8 | *Felis catus* | 96.856843 | 3.252197 | 29.782 | 760198 |
| FS19 | PBEP0014 | *Felis catus* | 87.648307 | 2.650553 | 33.068 | 852906 |
| FS19 | PBEP0023 | *Felis catus* | 87.220844 | 2.666313 | 32.712 | 847048 |
| FS19 | PVIP0012 | *Felis catus* | 30.388756 | 0.97351 | 31.216 | 766663 |
| FS19 | PBEP0005 | *Felis catus* | 49.332079 | 1.51644 | 32.532 | 940866 |
| FS19 | PBEP0008 | *Felis catus* | 49.320579 | 1.528168 | 32.274 | 939748 |
| FS19 | PBEP0009 | *Felis catus* | 49.121241 | 1.508188 | 32.57 | 940176 |
| FS19 | PBEP0068 | *Felis catus* | 75.043667 | 2.265701 | 33.122 | 974334 |
| FS19 | PBEP0069 | *Felis catus* | 75.997938 | 2.354484 | 32.278 | 962071 |
| FS19 | PBEP0066 | *Felis catus* | 75.1633 | 2.299679 | 32.684 | 964114 |
| FS19 | PBEP0067 | *Felis catus* | 75.071234 | 2.287896 | 32.812 | 928363 |
| FS19 | PBEP0064 | *Felis catus* | 75.280656 | 2.28575 | 32.935 | 965174 |
| FS19 | PBEP0065 | *Felis catus* | 75.449196 | 2.293242 | 32.901 | 966873 |
| FS19 | PBEP0038 | *Felis catus* | 67.478661 | 2.06905 | 32.613 | 998387 |
| FS19 | PBEP0039 | *Felis catus* | 69.956229 | 2.136028 | 32.751 | 990065 |
| FS19 | PBEP0028 | *Felis catus* | 58.522595 | 1.79248 | 32.649 | 997491 |
| FS19 | PBEP0036 | *Felis catus* | 61.711387 | 1.896681 | 32.537 | 992332 |
| PBEP0014 | PBEP0014 | *Felis catus* | 0 | -1 | 0 | -1 |
| PBEP0014 | FS15 | *Felis catus* | 82.320955 | 2.499533 | 32.935 | 785997 |
| PBEP0014 | FS8 | *Felis catus* | 79.309033 | 2.459222 | 32.25 | 974613 |
| PBEP0014 | FS19 | *Felis catus* | 87.648307 | 2.650553 | 33.068 | 852906 |
| PBEP0014 | PBEP0023 | *Felis catus* | 66.197117 | 2.082669 | 31.785 | 1190087 |
| PBEP0014 | PVIP0012 | *Felis catus* | 23.948674 | 0.758141 | 31.589 | 1186896 |
| PBEP0014 | PBEP0005 | *Felis catus* | 37.477295 | 1.185413 | 31.615 | 1377780 |
| PBEP0014 | PBEP0008 | *Felis catus* | 37.671307 | 1.204158 | 31.284 | 1371085 |
| PBEP0014 | PBEP0009 | *Felis catus* | 37.416043 | 1.177416 | 31.778 | 1376009 |
| PBEP0014 | PBEP0068 | *Felis catus* | 53.490127 | 1.666208 | 32.103 | 1419107 |
| PBEP0014 | PBEP0069 | *Felis catus* | 54.881495 | 1.770938 | 30.99 | 1379219 |
| PBEP0014 | PBEP0066 | *Felis catus* | 54.002324 | 1.700729 | 31.752 | 1394879 |
| PBEP0014 | PBEP0067 | *Felis catus* | 53.889496 | 1.691444 | 31.86 | 1349399 |
| PBEP0014 | PBEP0064 | *Felis catus* | 54.118805 | 1.708585 | 31.675 | 1393811 |
| PBEP0014 | PBEP0065 | *Felis catus* | 54.12236 | 1.701883 | 31.801 | 1399547 |
| PBEP0014 | PBEP0038 | *Felis catus* | 48.951724 | 1.532748 | 31.937 | 1449191 |
| PBEP0014 | PBEP0039 | *Felis catus* | 50.68284 | 1.589837 | 31.879 | 1429287 |
| PBEP0014 | PBEP0028 | *Felis catus* | 43.426948 | 1.377289 | 31.531 | 1445233 |
| PBEP0014 | PBEP0036 | *Felis catus* | 45.649228 | 1.458796 | 31.292 | 1430275 |
| PBEP0023 | PBEP0023 | *Felis catus* | 0 | -1 | 0 | -1 |
| PBEP0023 | FS15 | *Felis catus* | 82.094914 | 2.511656 | 32.686 | 780733 |
| PBEP0023 | FS8 | *Felis catus* | 78.955336 | 2.462822 | 32.059 | 969790 |
| PBEP0023 | FS19 | *Felis catus* | 87.220844 | 2.666313 | 32.712 | 847048 |
| PBEP0023 | PBEP0014 | *Felis catus* | 66.197117 | 2.082669 | 31.785 | 1190087 |
| PBEP0023 | PVIP0012 | *Felis catus* | 23.837789 | 0.760999 | 31.324 | 1174225 |
| PBEP0023 | PBEP0005 | *Felis catus* | 37.35184 | 1.195286 | 31.249 | 1366269 |
| PBEP0023 | PBEP0008 | *Felis catus* | 37.504124 | 1.215381 | 30.858 | 1359709 |
| PBEP0023 | PBEP0009 | *Felis catus* | 37.248091 | 1.187525 | 31.366 | 1364656 |
| PBEP0023 | PBEP0068 | *Felis catus* | 53.251322 | 1.677011 | 31.754 | 1410661 |
| PBEP0023 | PBEP0069 | *Felis catus* | 54.652481 | 1.789079 | 30.548 | 1370795 |
| PBEP0023 | PBEP0066 | *Felis catus* | 53.74023 | 1.716968 | 31.299 | 1386675 |
| PBEP0023 | PBEP0067 | *Felis catus* | 53.715082 | 1.708844 | 31.434 | 1340270 |
| PBEP0023 | PBEP0064 | *Felis catus* | 53.891182 | 1.714463 | 31.433 | 1385607 |
| PBEP0023 | PBEP0065 | *Felis catus* | 53.891158 | 1.713174 | 31.457 | 1391553 |
| PBEP0023 | PBEP0038 | *Felis catus* | 48.685729 | 1.54992 | 31.412 | 1441364 |
| PBEP0023 | PBEP0039 | *Felis catus* | 50.451412 | 1.603709 | 31.459 | 1420513 |
| PBEP0023 | PBEP0028 | *Felis catus* | 43.196526 | 1.382828 | 31.238 | 1435808 |
| PBEP0023 | PBEP0036 | *Felis catus* | 45.449093 | 1.469906 | 30.92 | 1420822 |
| PVIP0012 | PVIP0012 | *Felis catus* | 0 | -1 | 0 | -1 |
| PVIP0012 | FS15 | *Felis catus* | 28.737047 | 0.917571 | 31.319 | 700068 |
| PVIP0012 | FS8 | *Felis catus* | 28.183928 | 0.906908 | 31.077 | 888823 |
| PVIP0012 | FS19 | *Felis catus* | 30.388756 | 0.97351 | 31.216 | 766663 |
| PVIP0012 | PBEP0014 | *Felis catus* | 23.948674 | 0.758141 | 31.589 | 1186896 |
| PVIP0012 | PBEP0023 | *Felis catus* | 23.837789 | 0.760999 | 31.324 | 1174225 |
| PVIP0012 | PBEP0005 | *Felis catus* | 25.634429 | 0.832234 | 30.802 | 1039423 |
| PVIP0012 | PBEP0008 | *Felis catus* | 25.673187 | 0.844428 | 30.403 | 1032625 |
| PVIP0012 | PBEP0009 | *Felis catus* | 25.638723 | 0.825598 | 31.055 | 1037937 |
| PVIP0012 | PBEP0068 | *Felis catus* | 24.329028 | 0.767674 | 31.692 | 1298418 |
| PVIP0012 | PBEP0069 | *Felis catus* | 24.554984 | 0.814497 | 30.147 | 1244679 |
| PVIP0012 | PBEP0066 | *Felis catus* | 24.290088 | 0.777031 | 31.26 | 1265061 |
| PVIP0012 | PBEP0067 | *Felis catus* | 24.293093 | 0.780768 | 31.114 | 1200500 |
| PVIP0012 | PBEP0064 | *Felis catus* | 24.218761 | 0.776431 | 31.192 | 1265322 |
| PVIP0012 | PBEP0065 | *Felis catus* | 24.356293 | 0.783999 | 31.067 | 1272804 |
| PVIP0012 | PBEP0038 | *Felis catus* | 24.560739 | 0.782955 | 31.369 | 1278932 |
| PVIP0012 | PBEP0039 | *Felis catus* | 24.501967 | 0.786403 | 31.157 | 1269500 |
| PVIP0012 | PBEP0028 | *Felis catus* | 25.061814 | 0.803433 | 31.193 | 1202629 |
| PVIP0012 | PBEP0036 | *Felis catus* | 24.969234 | 0.808118 | 30.898 | 1209863 |
| PBEP0005 | PBEP0005 | *Felis catus* | 0 | -1 | 0 | -1 |
| PBEP0005 | FS15 | *Felis catus* | 46.123892 | 1.4261 | 32.343 | 858528 |
| PBEP0005 | FS8 | *Felis catus* | 45.603351 | 1.436178 | 31.753 | 1074427 |
| PBEP0005 | FS19 | *Felis catus* | 49.332079 | 1.51644 | 32.532 | 940866 |
| PBEP0005 | PBEP0014 | *Felis catus* | 37.477295 | 1.185413 | 31.615 | 1377780 |
| PBEP0005 | PBEP0023 | *Felis catus* | 37.35184 | 1.195286 | 31.249 | 1366269 |
| PBEP0005 | PVIP0012 | *Felis catus* | 25.634429 | 0.832234 | 30.802 | 1039423 |
| PBEP0005 | PBEP0008 | *Felis catus* | 59.109367 | 1.954609 | 30.241 | 1046564 |
| PBEP0005 | PBEP0009 | *Felis catus* | 58.574828 | 1.900872 | 30.815 | 1052285 |
| PBEP0005 | PBEP0068 | *Felis catus* | 38.187278 | 1.224464 | 31.187 | 1466928 |
| PBEP0005 | PBEP0069 | *Felis catus* | 38.972885 | 1.292857 | 30.145 | 1424331 |
| PBEP0005 | PBEP0066 | *Felis catus* | 38.377598 | 1.235497 | 31.062 | 1441564 |
| PBEP0005 | PBEP0067 | *Felis catus* | 38.422704 | 1.239383 | 31.001 | 1384954 |
| PBEP0005 | PBEP0064 | *Felis catus* | 38.320087 | 1.237798 | 30.958 | 1442591 |
| PBEP0005 | PBEP0065 | *Felis catus* | 38.46402 | 1.24436 | 30.911 | 1448224 |
| PBEP0005 | PBEP0038 | *Felis catus* | 43.72363 | 1.403629 | 31.15 | 1392550 |
| PBEP0005 | PBEP0039 | *Felis catus* | 42.41455 | 1.37545 | 30.837 | 1402302 |
| PBEP0005 | PBEP0028 | *Felis catus* | 50.648952 | 1.643772 | 30.813 | 1261340 |
| PBEP0005 | PBEP0036 | *Felis catus* | 48.824514 | 1.58665 | 30.772 | 1290000 |
| PBEP0008 | PBEP0008 | *Felis catus* | 0 | -1 | 0 | -1 |
| PBEP0008 | FS15 | *Felis catus* | 46.164718 | 1.428339 | 32.321 | 857682 |
| PBEP0008 | FS8 | *Felis catus* | 45.538826 | 1.439044 | 31.645 | 1072055 |
| PBEP0008 | FS19 | *Felis catus* | 49.320579 | 1.528168 | 32.274 | 939748 |
| PBEP0008 | PBEP0014 | *Felis catus* | 37.671307 | 1.204158 | 31.284 | 1371085 |
| PBEP0008 | PBEP0023 | *Felis catus* | 37.504124 | 1.215381 | 30.858 | 1359709 |
| PBEP0008 | PVIP0012 | *Felis catus* | 25.673187 | 0.844428 | 30.403 | 1032625 |
| PBEP0008 | PBEP0005 | *Felis catus* | 59.109367 | 1.954609 | 30.241 | 1046564 |
| PBEP0008 | PBEP0009 | *Felis catus* | 58.850098 | 1.936179 | 30.395 | 1046302 |
| PBEP0008 | PBEP0068 | *Felis catus* | 38.376315 | 1.24739 | 30.765 | 1458998 |
| PBEP0008 | PBEP0069 | *Felis catus* | 39.171144 | 1.314889 | 29.79 | 1419140 |
| PBEP0008 | PBEP0066 | *Felis catus* | 38.564098 | 1.259783 | 30.612 | 1434541 |
| PBEP0008 | PBEP0067 | *Felis catus* | 38.617735 | 1.261538 | 30.612 | 1378483 |
| PBEP0008 | PBEP0064 | *Felis catus* | 38.535139 | 1.25923 | 30.602 | 1436094 |
| PBEP0008 | PBEP0065 | *Felis catus* | 38.647117 | 1.265993 | 30.527 | 1441722 |
| PBEP0008 | PBEP0038 | *Felis catus* | 43.885647 | 1.430986 | 30.668 | 1385733 |
| PBEP0008 | PBEP0039 | *Felis catus* | 42.62025 | 1.396354 | 30.523 | 1396048 |
| PBEP0008 | PBEP0028 | *Felis catus* | 50.883481 | 1.662097 | 30.614 | 1255154 |
| PBEP0008 | PBEP0036 | *Felis catus* | 49.032855 | 1.617148 | 30.321 | 1285300 |
| PBEP0009 | PBEP0009 | *Felis catus* | 0 | -1 | 0 | -1 |
| PBEP0009 | FS15 | *Felis catus* | 45.917531 | 1.403135 | 32.725 | 858237 |
| PBEP0009 | FS8 | *Felis catus* | 45.358966 | 1.405665 | 32.269 | 1073212 |
| PBEP0009 | FS19 | *Felis catus* | 49.121241 | 1.508188 | 32.57 | 940176 |
| PBEP0009 | PBEP0014 | *Felis catus* | 37.416043 | 1.177416 | 31.778 | 1376009 |
| PBEP0009 | PBEP0023 | *Felis catus* | 37.248091 | 1.187525 | 31.366 | 1364656 |
| PBEP0009 | PVIP0012 | *Felis catus* | 25.638723 | 0.825598 | 31.055 | 1037937 |
| PBEP0009 | PBEP0005 | *Felis catus* | 58.574828 | 1.900872 | 30.815 | 1052285 |
| PBEP0009 | PBEP0008 | *Felis catus* | 58.850098 | 1.936179 | 30.395 | 1046302 |
| PBEP0009 | PBEP0068 | *Felis catus* | 38.118426 | 1.216623 | 31.331 | 1465594 |
| PBEP0009 | PBEP0069 | *Felis catus* | 38.861658 | 1.284087 | 30.264 | 1422781 |
| PBEP0009 | PBEP0066 | *Felis catus* | 38.251629 | 1.225227 | 31.22 | 1440249 |
| PBEP0009 | PBEP0067 | *Felis catus* | 38.330881 | 1.235669 | 31.02 | 1383051 |
| PBEP0009 | PBEP0064 | *Felis catus* | 38.248548 | 1.229581 | 31.107 | 1440539 |
| PBEP0009 | PBEP0065 | *Felis catus* | 38.352801 | 1.237209 | 30.999 | 1446463 |
| PBEP0009 | PBEP0038 | *Felis catus* | 43.641591 | 1.394671 | 31.292 | 1391361 |
| PBEP0009 | PBEP0039 | *Felis catus* | 42.336655 | 1.367342 | 30.963 | 1400471 |
| PBEP0009 | PBEP0028 | *Felis catus* | 50.585553 | 1.634403 | 30.95 | 1259817 |
| PBEP0009 | PBEP0036 | *Felis catus* | 48.71554 | 1.581173 | 30.81 | 1288682 |
| PBEP0068 | PBEP0068 | *Felis catus* | 0 | -1 | 0 | -1 |
| PBEP0068 | FS15 | *Felis catus* | 69.950411 | 2.10229 | 33.273 | 894240 |
| PBEP0068 | FS8 | *Felis catus* | 68.206189 | 2.100244 | 32.475 | 1107559 |
| PBEP0068 | FS19 | *Felis catus* | 75.043667 | 2.265701 | 33.122 | 974334 |
| PBEP0068 | PBEP0014 | *Felis catus* | 53.490127 | 1.666208 | 32.103 | 1419107 |
| PBEP0068 | PBEP0023 | *Felis catus* | 53.251322 | 1.677011 | 31.754 | 1410661 |
| PBEP0068 | PVIP0012 | *Felis catus* | 24.329028 | 0.767674 | 31.692 | 1298418 |
| PBEP0068 | PBEP0005 | *Felis catus* | 38.187278 | 1.224464 | 31.187 | 1466928 |
| PBEP0068 | PBEP0008 | *Felis catus* | 38.376315 | 1.24739 | 30.765 | 1458998 |
| PBEP0068 | PBEP0009 | *Felis catus* | 38.118426 | 1.216623 | 31.331 | 1465594 |
| PBEP0068 | PBEP0069 | *Felis catus* | 58.441624 | 1.913531 | 30.541 | 1411327 |
| PBEP0068 | PBEP0066 | *Felis catus* | 57.58135 | 1.824227 | 31.565 | 1426744 |
| PBEP0068 | PBEP0067 | *Felis catus* | 56.680701 | 1.803141 | 31.434 | 1398764 |
| PBEP0068 | PBEP0064 | *Felis catus* | 56.393183 | 1.796036 | 31.399 | 1443298 |
| PBEP0068 | PBEP0065 | *Felis catus* | 56.737073 | 1.812383 | 31.305 | 1446165 |
| PBEP0068 | PBEP0038 | *Felis catus* | 50.571 | 1.608067 | 31.448 | 1511299 |
| PBEP0068 | PBEP0039 | *Felis catus* | 52.322734 | 1.654707 | 31.621 | 1487696 |
| PBEP0068 | PBEP0028 | *Felis catus* | 44.533386 | 1.423341 | 31.288 | 1518078 |
| PBEP0068 | PBEP0036 | *Felis catus* | 46.901036 | 1.520153 | 30.853 | 1498384 |
| PBEP0069 | PBEP0069 | *Felis catus* | 0 | -1 | 0 | -1 |
| PBEP0069 | FS15 | *Felis catus* | 71.417436 | 2.211924 | 32.287 | 882128 |
| PBEP0069 | FS8 | *Felis catus* | 69.39284 | 2.205899 | 31.458 | 1090504 |
| PBEP0069 | FS19 | *Felis catus* | 75.997938 | 2.354484 | 32.278 | 962071 |
| PBEP0069 | PBEP0014 | *Felis catus* | 54.881495 | 1.770938 | 30.99 | 1379219 |
| PBEP0069 | PBEP0023 | *Felis catus* | 54.652481 | 1.789079 | 30.548 | 1370795 |
| PBEP0069 | PVIP0012 | *Felis catus* | 24.554984 | 0.814497 | 30.147 | 1244679 |
| PBEP0069 | PBEP0005 | *Felis catus* | 38.972885 | 1.292857 | 30.145 | 1424331 |
| PBEP0069 | PBEP0008 | *Felis catus* | 39.171144 | 1.314889 | 29.79 | 1419140 |
| PBEP0069 | PBEP0009 | *Felis catus* | 38.861658 | 1.284087 | 30.264 | 1422781 |
| PBEP0069 | PBEP0068 | *Felis catus* | 58.441624 | 1.913531 | 30.541 | 1411327 |
| PBEP0069 | PBEP0066 | *Felis catus* | 58.552373 | 1.922908 | 30.45 | 1395854 |
| PBEP0069 | PBEP0067 | *Felis catus* | 57.836758 | 1.902992 | 30.393 | 1363949 |
| PBEP0069 | PBEP0064 | *Felis catus* | 57.746294 | 1.89774 | 30.429 | 1407299 |
| PBEP0069 | PBEP0065 | *Felis catus* | 58.039258 | 1.919653 | 30.234 | 1410521 |
| PBEP0069 | PBEP0038 | *Felis catus* | 51.890591 | 1.71557 | 30.247 | 1468425 |
| PBEP0069 | PBEP0039 | *Felis catus* | 53.71418 | 1.762043 | 30.484 | 1450464 |
| PBEP0069 | PBEP0028 | *Felis catus* | 45.55532 | 1.507132 | 30.226 | 1477220 |
| PBEP0069 | PBEP0036 | *Felis catus* | 47.900895 | 1.604823 | 29.848 | 1461155 |
| PBEP0066 | PBEP0066 | *Felis catus* | 0 | -1 | 0 | -1 |
| PBEP0066 | FS15 | *Felis catus* | 70.576964 | 2.153776 | 32.769 | 884207 |
| PBEP0066 | FS8 | *Felis catus* | 68.65733 | 2.146608 | 31.984 | 1094816 |
| PBEP0066 | FS19 | *Felis catus* | 75.1633 | 2.299679 | 32.684 | 964114 |
| PBEP0066 | PBEP0014 | *Felis catus* | 54.002324 | 1.700729 | 31.752 | 1394879 |
| PBEP0066 | PBEP0023 | *Felis catus* | 53.74023 | 1.716968 | 31.299 | 1386675 |
| PBEP0066 | PVIP0012 | *Felis catus* | 24.290088 | 0.777031 | 31.26 | 1265061 |
| PBEP0066 | PBEP0005 | *Felis catus* | 38.377598 | 1.235497 | 31.062 | 1441564 |
| PBEP0066 | PBEP0008 | *Felis catus* | 38.564098 | 1.259783 | 30.612 | 1434541 |
| PBEP0066 | PBEP0009 | *Felis catus* | 38.251629 | 1.225227 | 31.22 | 1440249 |
| PBEP0066 | PBEP0068 | *Felis catus* | 57.58135 | 1.824227 | 31.565 | 1426744 |
| PBEP0066 | PBEP0069 | *Felis catus* | 58.552373 | 1.922908 | 30.45 | 1395854 |
| PBEP0066 | PBEP0067 | *Felis catus* | 57.043378 | 1.830176 | 31.168 | 1377887 |
| PBEP0066 | PBEP0064 | *Felis catus* | 56.830802 | 1.8231 | 31.173 | 1421893 |
| PBEP0066 | PBEP0065 | *Felis catus* | 57.080744 | 1.832016 | 31.157 | 1425738 |
| PBEP0066 | PBEP0038 | *Felis catus* | 50.912261 | 1.634442 | 31.15 | 1487144 |
| PBEP0066 | PBEP0039 | *Felis catus* | 52.786739 | 1.686925 | 31.292 | 1465598 |
| PBEP0066 | PBEP0028 | *Felis catus* | 44.785824 | 1.435889 | 31.19 | 1493950 |
| PBEP0066 | PBEP0036 | *Felis catus* | 47.165809 | 1.535822 | 30.71 | 1476621 |
| PBEP0067 | PBEP0067 | *Felis catus* | 0 | -1 | 0 | -1 |
| PBEP0067 | FS15 | *Felis catus* | 70.503829 | 2.15352 | 32.739 | 851996 |
| PBEP0067 | FS8 | *Felis catus* | 68.672333 | 2.137885 | 32.122 | 1055827 |
| PBEP0067 | FS19 | *Felis catus* | 75.071234 | 2.287896 | 32.812 | 928363 |
| PBEP0067 | PBEP0014 | *Felis catus* | 53.889496 | 1.691444 | 31.86 | 1349399 |
| PBEP0067 | PBEP0023 | *Felis catus* | 53.715082 | 1.708844 | 31.434 | 1340270 |
| PBEP0067 | PVIP0012 | *Felis catus* | 24.293093 | 0.780768 | 31.114 | 1200500 |
| PBEP0067 | PBEP0005 | *Felis catus* | 38.422704 | 1.239383 | 31.001 | 1384954 |
| PBEP0067 | PBEP0008 | *Felis catus* | 38.617735 | 1.261538 | 30.612 | 1378483 |
| PBEP0067 | PBEP0009 | *Felis catus* | 38.330881 | 1.235669 | 31.02 | 1383051 |
| PBEP0067 | PBEP0068 | *Felis catus* | 56.680701 | 1.803141 | 31.434 | 1398764 |
| PBEP0067 | PBEP0069 | *Felis catus* | 57.836758 | 1.902992 | 30.393 | 1363949 |
| PBEP0067 | PBEP0066 | *Felis catus* | 57.043378 | 1.830176 | 31.168 | 1377887 |
| PBEP0067 | PBEP0064 | *Felis catus* | 57.033927 | 1.826729 | 31.222 | 1378841 |
| PBEP0067 | PBEP0065 | *Felis catus* | 57.425261 | 1.845202 | 31.121 | 1380255 |
| PBEP0067 | PBEP0038 | *Felis catus* | 50.865001 | 1.628588 | 31.233 | 1444001 |
| PBEP0067 | PBEP0039 | *Felis catus* | 52.606408 | 1.68338 | 31.25 | 1424429 |
| PBEP0067 | PBEP0028 | *Felis catus* | 44.835111 | 1.441555 | 31.102 | 1445197 |
| PBEP0067 | PBEP0036 | *Felis catus* | 47.149441 | 1.536355 | 30.689 | 1429407 |
| PBEP0064 | PBEP0064 | *Felis catus* | 0 | -1 | 0 | -1 |
| PBEP0064 | FS15 | *Felis catus* | 70.628207 | 2.143967 | 32.943 | 885700 |
| PBEP0064 | FS8 | *Felis catus* | 68.828178 | 2.138988 | 32.178 | 1095339 |
| PBEP0064 | FS19 | *Felis catus* | 75.280656 | 2.28575 | 32.935 | 965174 |
| PBEP0064 | PBEP0014 | *Felis catus* | 54.118805 | 1.708585 | 31.675 | 1393811 |
| PBEP0064 | PBEP0023 | *Felis catus* | 53.891182 | 1.714463 | 31.433 | 1385607 |
| PBEP0064 | PVIP0012 | *Felis catus* | 24.218761 | 0.776431 | 31.192 | 1265322 |
| PBEP0064 | PBEP0005 | *Felis catus* | 38.320087 | 1.237798 | 30.958 | 1442591 |
| PBEP0064 | PBEP0008 | *Felis catus* | 38.535139 | 1.25923 | 30.602 | 1436094 |
| PBEP0064 | PBEP0009 | *Felis catus* | 38.248548 | 1.229581 | 31.107 | 1440539 |
| PBEP0064 | PBEP0068 | *Felis catus* | 56.393183 | 1.796036 | 31.399 | 1443298 |
| PBEP0064 | PBEP0069 | *Felis catus* | 57.746294 | 1.89774 | 30.429 | 1407299 |
| PBEP0064 | PBEP0066 | *Felis catus* | 56.830802 | 1.8231 | 31.173 | 1421893 |
| PBEP0064 | PBEP0067 | *Felis catus* | 57.033927 | 1.826729 | 31.222 | 1378841 |
| PBEP0064 | PBEP0065 | *Felis catus* | 57.47477 | 1.844152 | 31.166 | 1420493 |
| PBEP0064 | PBEP0038 | *Felis catus* | 50.949881 | 1.634895 | 31.164 | 1487328 |
| PBEP0064 | PBEP0039 | *Felis catus* | 52.791184 | 1.687717 | 31.28 | 1466847 |
| PBEP0064 | PBEP0028 | *Felis catus* | 44.772867 | 1.436449 | 31.169 | 1494998 |
| PBEP0064 | PBEP0036 | *Felis catus* | 47.143791 | 1.536757 | 30.677 | 1477374 |
| PBEP0065 | PBEP0065 | *Felis catus* | 0 | -1 | 0 | -1 |
| PBEP0065 | FS15 | *Felis catus* | 70.682337 | 2.159481 | 32.731 | 887683 |
| PBEP0065 | FS8 | *Felis catus* | 68.826358 | 2.144858 | 32.089 | 1098540 |
| PBEP0065 | FS19 | *Felis catus* | 75.449196 | 2.293242 | 32.901 | 966873 |
| PBEP0065 | PBEP0014 | *Felis catus* | 54.12236 | 1.701883 | 31.801 | 1399547 |
| PBEP0065 | PBEP0023 | *Felis catus* | 53.891158 | 1.713174 | 31.457 | 1391553 |
| PBEP0065 | PVIP0012 | *Felis catus* | 24.356293 | 0.783999 | 31.067 | 1272804 |
| PBEP0065 | PBEP0005 | *Felis catus* | 38.46402 | 1.24436 | 30.911 | 1448224 |
| PBEP0065 | PBEP0008 | *Felis catus* | 38.647117 | 1.265993 | 30.527 | 1441722 |
| PBEP0065 | PBEP0009 | *Felis catus* | 38.352801 | 1.237209 | 30.999 | 1446463 |
| PBEP0065 | PBEP0068 | *Felis catus* | 56.737073 | 1.812383 | 31.305 | 1446165 |
| PBEP0065 | PBEP0069 | *Felis catus* | 58.039258 | 1.919653 | 30.234 | 1410521 |
| PBEP0065 | PBEP0066 | *Felis catus* | 57.080744 | 1.832016 | 31.157 | 1425738 |
| PBEP0065 | PBEP0067 | *Felis catus* | 57.425261 | 1.845202 | 31.121 | 1380255 |
| PBEP0065 | PBEP0064 | *Felis catus* | 57.47477 | 1.844152 | 31.166 | 1420493 |
| PBEP0065 | PBEP0038 | *Felis catus* | 51.049324 | 1.641499 | 31.099 | 1493130 |
| PBEP0065 | PBEP0039 | *Felis catus* | 52.893594 | 1.693741 | 31.229 | 1471647 |
| PBEP0065 | PBEP0028 | *Felis catus* | 44.908315 | 1.455199 | 30.861 | 1499890 |
| PBEP0065 | PBEP0036 | *Felis catus* | 47.319543 | 1.547917 | 30.57 | 1482257 |
| PBEP0038 | PBEP0038 | *Felis catus* | 0 | -1 | 0 | -1 |
| PBEP0038 | FS15 | *Felis catus* | 62.92979 | 1.919277 | 32.788 | 915250 |
| PBEP0038 | FS8 | *Felis catus* | 61.557468 | 1.923598 | 32.001 | 1135282 |
| PBEP0038 | FS19 | *Felis catus* | 67.478661 | 2.06905 | 32.613 | 998387 |
| PBEP0038 | PBEP0014 | *Felis catus* | 48.951724 | 1.532748 | 31.937 | 1449191 |
| PBEP0038 | PBEP0023 | *Felis catus* | 48.685729 | 1.54992 | 31.412 | 1441364 |
| PBEP0038 | PVIP0012 | *Felis catus* | 24.560739 | 0.782955 | 31.369 | 1278932 |
| PBEP0038 | PBEP0005 | *Felis catus* | 43.72363 | 1.403629 | 31.15 | 1392550 |
| PBEP0038 | PBEP0008 | *Felis catus* | 43.885647 | 1.430986 | 30.668 | 1385733 |
| PBEP0038 | PBEP0009 | *Felis catus* | 43.641591 | 1.394671 | 31.292 | 1391361 |
| PBEP0038 | PBEP0068 | *Felis catus* | 50.571 | 1.608067 | 31.448 | 1511299 |
| PBEP0038 | PBEP0069 | *Felis catus* | 51.890591 | 1.71557 | 30.247 | 1468425 |
| PBEP0038 | PBEP0066 | *Felis catus* | 50.912261 | 1.634442 | 31.15 | 1487144 |
| PBEP0038 | PBEP0067 | *Felis catus* | 50.865001 | 1.628588 | 31.233 | 1444001 |
| PBEP0038 | PBEP0064 | *Felis catus* | 50.949881 | 1.634895 | 31.164 | 1487328 |
| PBEP0038 | PBEP0065 | *Felis catus* | 51.049324 | 1.641499 | 31.099 | 1493130 |
| PBEP0038 | PBEP0039 | *Felis catus* | 52.325796 | 1.674354 | 31.251 | 1462386 |
| PBEP0038 | PBEP0028 | *Felis catus* | 48.251508 | 1.536723 | 31.399 | 1453227 |
| PBEP0038 | PBEP0036 | *Felis catus* | 49.911117 | 1.611623 | 30.969 | 1441111 |
| PBEP0039 | PBEP0039 | *Felis catus* | 0 | -1 | 0 | -1 |
| PBEP0039 | FS15 | *Felis catus* | 65.392363 | 1.992234 | 32.824 | 907600 |
| PBEP0039 | FS8 | *Felis catus* | 64.069036 | 1.998685 | 32.056 | 1123996 |
| PBEP0039 | FS19 | *Felis catus* | 69.956229 | 2.136028 | 32.751 | 990065 |
| PBEP0039 | PBEP0014 | *Felis catus* | 50.68284 | 1.589837 | 31.879 | 1429287 |
| PBEP0039 | PBEP0023 | *Felis catus* | 50.451412 | 1.603709 | 31.459 | 1420513 |
| PBEP0039 | PVIP0012 | *Felis catus* | 24.501967 | 0.786403 | 31.157 | 1269500 |
| PBEP0039 | PBEP0005 | *Felis catus* | 42.41455 | 1.37545 | 30.837 | 1402302 |
| PBEP0039 | PBEP0008 | *Felis catus* | 42.62025 | 1.396354 | 30.523 | 1396048 |
| PBEP0039 | PBEP0009 | *Felis catus* | 42.336655 | 1.367342 | 30.963 | 1400471 |
| PBEP0039 | PBEP0068 | *Felis catus* | 52.322734 | 1.654707 | 31.621 | 1487696 |
| PBEP0039 | PBEP0069 | *Felis catus* | 53.71418 | 1.762043 | 30.484 | 1450464 |
| PBEP0039 | PBEP0066 | *Felis catus* | 52.786739 | 1.686925 | 31.292 | 1465598 |
| PBEP0039 | PBEP0067 | *Felis catus* | 52.606408 | 1.68338 | 31.25 | 1424429 |
| PBEP0039 | PBEP0064 | *Felis catus* | 52.791184 | 1.687717 | 31.28 | 1466847 |
| PBEP0039 | PBEP0065 | *Felis catus* | 52.893594 | 1.693741 | 31.229 | 1471647 |
| PBEP0039 | PBEP0038 | *Felis catus* | 52.325796 | 1.674354 | 31.251 | 1462386 |
| PBEP0039 | PBEP0028 | *Felis catus* | 47.650117 | 1.532367 | 31.096 | 1459517 |
| PBEP0039 | PBEP0036 | *Felis catus* | 49.743939 | 1.61294 | 30.841 | 1444072 |
| PBEP0028 | PBEP0028 | *Felis catus* | 0 | -1 | 0 | -1 |
| PBEP0028 | FS15 | *Felis catus* | 54.652416 | 1.671718 | 32.692 | 911959 |
| PBEP0028 | FS8 | *Felis catus* | 53.800577 | 1.676275 | 32.095 | 1134106 |
| PBEP0028 | FS19 | *Felis catus* | 58.522595 | 1.79248 | 32.649 | 997491 |
| PBEP0028 | PBEP0014 | *Felis catus* | 43.426948 | 1.377289 | 31.531 | 1445233 |
| PBEP0028 | PBEP0023 | *Felis catus* | 43.196526 | 1.382828 | 31.238 | 1435808 |
| PBEP0028 | PVIP0012 | *Felis catus* | 25.061814 | 0.803433 | 31.193 | 1202629 |
| PBEP0028 | PBEP0005 | *Felis catus* | 50.648952 | 1.643772 | 30.813 | 1261340 |
| PBEP0028 | PBEP0008 | *Felis catus* | 50.883481 | 1.662097 | 30.614 | 1255154 |
| PBEP0028 | PBEP0009 | *Felis catus* | 50.585553 | 1.634403 | 30.95 | 1259817 |
| PBEP0028 | PBEP0068 | *Felis catus* | 44.533386 | 1.423341 | 31.288 | 1518078 |
| PBEP0028 | PBEP0069 | *Felis catus* | 45.55532 | 1.507132 | 30.226 | 1477220 |
| PBEP0028 | PBEP0066 | *Felis catus* | 44.785824 | 1.435889 | 31.19 | 1493950 |
| PBEP0028 | PBEP0067 | *Felis catus* | 44.835111 | 1.441555 | 31.102 | 1445197 |
| PBEP0028 | PBEP0064 | *Felis catus* | 44.772867 | 1.436449 | 31.169 | 1494998 |
| PBEP0028 | PBEP0065 | *Felis catus* | 44.908315 | 1.455199 | 30.861 | 1499890 |
| PBEP0028 | PBEP0038 | *Felis catus* | 48.251508 | 1.536723 | 31.399 | 1453227 |
| PBEP0028 | PBEP0039 | *Felis catus* | 47.650117 | 1.532367 | 31.096 | 1459517 |
| PBEP0028 | PBEP0036 | *Felis catus* | 51.988861 | 1.676199 | 31.016 | 1360165 |
| PBEP0036 | PBEP0036 | *Felis catus* | 0 | -1 | 0 | -1 |
| PBEP0036 | FS15 | *Felis catus* | 57.967153 | 1.790102 | 32.382 | 907314 |
| PBEP0036 | FS8 | *Felis catus* | 56.795691 | 1.798561 | 31.578 | 1126856 |
| PBEP0036 | FS19 | *Felis catus* | 61.711387 | 1.896681 | 32.537 | 992332 |
| PBEP0036 | PBEP0014 | *Felis catus* | 45.649228 | 1.458796 | 31.292 | 1430275 |
| PBEP0036 | PBEP0023 | *Felis catus* | 45.449093 | 1.469906 | 30.92 | 1420822 |
| PBEP0036 | PVIP0012 | *Felis catus* | 24.969234 | 0.808118 | 30.898 | 1209863 |
| PBEP0036 | PBEP0005 | *Felis catus* | 48.824514 | 1.58665 | 30.772 | 1290000 |
| PBEP0036 | PBEP0008 | *Felis catus* | 49.032855 | 1.617148 | 30.321 | 1285300 |
| PBEP0036 | PBEP0009 | *Felis catus* | 48.71554 | 1.581173 | 30.81 | 1288682 |
| PBEP0036 | PBEP0068 | *Felis catus* | 46.901036 | 1.520153 | 30.853 | 1498384 |
| PBEP0036 | PBEP0069 | *Felis catus* | 47.900895 | 1.604823 | 29.848 | 1461155 |
| PBEP0036 | PBEP0066 | *Felis catus* | 47.165809 | 1.535822 | 30.71 | 1476621 |
| PBEP0036 | PBEP0067 | *Felis catus* | 47.149441 | 1.536355 | 30.689 | 1429407 |
| PBEP0036 | PBEP0064 | *Felis catus* | 47.143791 | 1.536757 | 30.677 | 1477374 |
| PBEP0036 | PBEP0065 | *Felis catus* | 47.319543 | 1.547917 | 30.57 | 1482257 |
| PBEP0036 | PBEP0038 | *Felis catus* | 49.911117 | 1.611623 | 30.969 | 1441111 |
| PBEP0036 | PBEP0039 | *Felis catus* | 49.743939 | 1.61294 | 30.841 | 1444072 |
| PBEP0036 | PBEP0028 | *Felis catus* | 51.988861 | 1.676199 | 31.016 | 1360165 |

**Table S4.** D-statistics of leopard cats from modern Asia and ancient China (Data of Fig. S11).

| Outgroup | Pop1 | Pop2 | Pop3 | D | std.err | Z | SNPs |
| --- | --- | --- | --- | --- | --- | --- | --- |
| *Felis catus* | PBEP0068 | Russian Far East | FS8 | 0.001928 | 0.000114 | 16.92 | 2089721 |
| *Felis catus* | PBEP0069 | Russian Far East | FS8 | 0.002137 | 0.00012 | 17.775 | 2086193 |
| *Felis catus* | PBEP0066 | Russian Far East | FS8 | 0.002049 | 0.000116 | 17.679 | 2082559 |
| *Felis catus* | PBEP0067 | Russian Far East | FS8 | 0.001995 | 0.00012 | 16.601 | 2087328 |
| *Felis catus* | PBEP0064 | Russian Far East | FS8 | 0.002102 | 0.000117 | 17.903 | 2095996 |
| *Felis catus* | PBEP0065 | Russian Far East | FS8 | 0.001924 | 0.000118 | 16.301 | 2081063 |
| *Felis catus* | PBEP0038 | Russian Far East | FS8 | 0.001492 | 0.00011 | 13.516 | 2090739 |
| *Felis catus* | PBEP0039 | Russian Far East | FS8 | 0.001671 | 0.000107 | 15.668 | 2089949 |
| *Felis catus* | PBEP0028 | Russian Far East | FS8 | 0.000921 | 0.000089 | 10.387 | 2089592 |
| *Felis catus* | PBEP0036 | Russian Far East | FS8 | 0.001149 | 0.000092 | 12.492 | 2090655 |
| *Felis catus* | PBEP0005 | Russian Far East | FS8 | 0.000262 | 0.000071 | 3.703 | 2096009 |
| *Felis catus* | PBEP0008 | Russian Far East | FS8 | 0.000294 | 0.000074 | 3.954 | 2090335 |
| *Felis catus* | PBEP0009 | Russian Far East | FS8 | 0.000261 | 0.000073 | 3.573 | 2090861 |
| *Felis catus* | PVIP0012 | Russian Far East | FS8 | 0.000202 | 0.000064 | 3.173 | 2085955 |
| *Felis catus* | PBEP0068 | Russian Far East | FS15 | 0.002408 | 0.00015 | 16.032 | 2335310 |
| *Felis catus* | PBEP0069 | Russian Far East | FS15 | 0.002671 | 0.000148 | 18.103 | 2331886 |
| *Felis catus* | PBEP0066 | Russian Far East | FS15 | 0.002503 | 0.000149 | 16.786 | 2328008 |
| *Felis catus* | PBEP0067 | Russian Far East | FS15 | 0.002461 | 0.000155 | 15.885 | 2333039 |
| *Felis catus* | PBEP0064 | Russian Far East | FS15 | 0.002494 | 0.000152 | 16.439 | 2341941 |
| *Felis catus* | PBEP0065 | Russian Far East | FS15 | 0.002366 | 0.000155 | 15.312 | 2326744 |
| *Felis catus* | PBEP0038 | Russian Far East | FS15 | 0.0017 | 0.000138 | 12.338 | 2336410 |
| *Felis catus* | PBEP0039 | Russian Far East | FS15 | 0.001881 | 0.000139 | 13.565 | 2335622 |
| *Felis catus* | PBEP0028 | Russian Far East | FS15 | 0.000912 | 0.000109 | 8.337 | 2335303 |
| *Felis catus* | PBEP0036 | Russian Far East | FS15 | 0.001289 | 0.000102 | 12.622 | 2336387 |
| *Felis catus* | PBEP0005 | Russian Far East | FS15 | 0.000297 | 0.000087 | 3.432 | 2342306 |
| *Felis catus* | PBEP0008 | Russian Far East | FS15 | 0.000306 | 0.000083 | 3.709 | 2336127 |
| *Felis catus* | PBEP0009 | Russian Far East | FS15 | 0.00028 | 0.000086 | 3.264 | 2336820 |
| *Felis catus* | PVIP0012 | Russian Far East | FS15 | 0.000264 | 0.000071 | 3.728 | 2331636 |
| *Felis catus* | PBEP0068 | Russian Far East | FS19 | 0.002581 | 0.00012 | 21.513 | 2613447 |
| *Felis catus* | PBEP0069 | Russian Far East | FS19 | 0.002805 | 0.000121 | 23.159 | 2608527 |
| *Felis catus* | PBEP0066 | Russian Far East | FS19 | 0.002478 | 0.000121 | 20.545 | 2603553 |
| *Felis catus* | PBEP0067 | Russian Far East | FS19 | 0.002569 | 0.000122 | 21.055 | 2609780 |
| *Felis catus* | PBEP0064 | Russian Far East | FS19 | 0.002637 | 0.000118 | 22.31 | 2621480 |
| *Felis catus* | PBEP0065 | Russian Far East | FS19 | 0.002566 | 0.000118 | 21.719 | 2600836 |
| *Felis catus* | PBEP0038 | Russian Far East | FS19 | 0.001884 | 0.000108 | 17.379 | 2614086 |
| *Felis catus* | PBEP0039 | Russian Far East | FS19 | 0.002145 | 0.000107 | 20.029 | 2612845 |
| *Felis catus* | PBEP0028 | Russian Far East | FS19 | 0.00111 | 0.00009 | 12.295 | 2612407 |
| *Felis catus* | PBEP0036 | Russian Far East | FS19 | 0.00141 | 0.000093 | 15.156 | 2614254 |
| *Felis catus* | PBEP0005 | Russian Far East | FS19 | 0.000399 | 0.000075 | 5.324 | 2621116 |
| *Felis catus* | PBEP0008 | Russian Far East | FS19 | 0.000491 | 0.000076 | 6.437 | 2613456 |
| *Felis catus* | PBEP0009 | Russian Far East | FS19 | 0.000408 | 0.000078 | 5.198 | 2614325 |
| *Felis catus* | PVIP0012 | Russian Far East | FS19 | 0.000269 | 0.000066 | 4.061 | 2607464 |

**Table S5.** Pairwise f3-statistics of the genetic relationships between various populations of ancient or modern Felis species (Data of Fig. 3B).

| Pop1 | Pop2 | Outgroup | f_3 | std.err | Z | SNPs |
| --- | --- | --- | --- | --- | --- | --- |
| FS1 | Modern_China/East_Asia_catus | *Felis margarita* | 180.21786 | 11.322142 | 15.92 | 81007 |
| FS1 | FS21 | *Felis margarita* | 179.38697 | 11.588262 | 15.48 | 50863 |
| FS1 | FS2 | *Felis margarita* | 175.83828 | 11.48101 | 15.32 | 53082 |
| FS1 | FS12 | *Felis margarita* | 172.01661 | 11.573873 | 14.86 | 58323 |
| FS1 | Levant lybica | *Felis margarita* | 151.32251 | 9.438672 | 16.03 | 77866 |
| FS1 | Dhzankent | *Felis margarita* | 150.71702 | 55.144777 | 2.733 | 1878 |
| FS1 | modern_European_catus | *Felis margarita* | 139.11256 | 8.677047 | 16.03 | 81937 |
| FS1 | Ancient_European/Anatolia_catus | *Felis margarita* | 122.01078 | 17.430569 | 7 | 8738 |
| FS1 | ornata | *Felis margarita* | 119.73996 | 7.444453 | 16.08 | 66125 |
| FS1 | silvestries | *Felis margarita* | 118.399 | 7.344451 | 16.09 | 70271 |
| FS1 | bieti | *Felis margarita* | 112.93601 | 7.047862 | 16.02 | 60043 |
| FS2 | FS1 | *Felis margarita* | 175.83828 | 11.48101 | 15.32 | 53082 |
| FS2 | Modern_China/East_Asia_catus | *Felis margarita* | 174.72111 | 11.255582 | 15.52 | 84297 |
| FS2 | FS21 | *Felis margarita* | 171.42282 | 11.359815 | 15.09 | 53303 |
| FS2 | FS12 | *Felis margarita* | 167.05408 | 11.598646 | 14.4 | 60457 |
| FS2 | Dhzankent | *Felis margarita* | 165.10907 | 58.683773 | 2.814 | 1966 |
| FS2 | Levant lybica | *Felis margarita* | 146.89282 | 9.435001 | 15.57 | 81263 |
| FS2 | modern_European_catus | *Felis margarita* | 134.72156 | 8.629847 | 15.61 | 85434 |
| FS2 | ornata | *Felis margarita* | 115.6054 | 7.403732 | 15.61 | 69362 |
| FS2 | Ancient_European/Anatolia_catus | *Felis margarita* | 114.05922 | 14.730644 | 7.743 | 9158 |
| FS2 | silvestries | *Felis margarita* | 113.90879 | 7.306579 | 15.59 | 73504 |
| FS2 | bieti | *Felis margarita* | 109.28301 | 7.022039 | 15.56 | 63007 |
| FS12 | Dhzankent | *Felis margarita* | 180.25164 | 77.922663 | 2.313 | 2125 |
| FS12 | Modern_China/East_Asia_catus | *Felis margarita* | 172.25718 | 11.165351 | 15.43 | 91443 |
| FS12 | FS1 | *Felis margarita* | 172.01661 | 11.573873 | 14.86 | 58323 |
| FS12 | FS21 | *Felis margarita* | 171.49798 | 11.460161 | 14.97 | 57531 |
| FS12 | FS2 | *Felis margarita* | 167.05408 | 11.598646 | 14.4 | 60457 |
| FS12 | Levant lybica | *Felis margarita* | 145.00226 | 9.403209 | 15.42 | 88355 |
| FS12 | modern_European_catus | *Felis margarita* | 131.4479 | 8.489932 | 15.48 | 93524 |
| FS12 | ornata | *Felis margarita* | 114.95499 | 7.393877 | 15.55 | 75907 |
| FS12 | Ancient_European/Anatolia_catus | *Felis margarita* | 112.61497 | 15.776276 | 7.138 | 10099 |
| FS12 | silvestries | *Felis margarita* | 111.98298 | 7.232932 | 15.48 | 80847 |
| FS12 | bieti | *Felis margarita* | 108.95904 | 7.022332 | 15.52 | 69196 |
| FS21 | Modern_China/East_Asia_catus | *Felis margarita* | 179.48462 | 11.363918 | 15.79 | 80396 |
| FS21 | FS1 | *Felis margarita* | 179.38697 | 11.588262 | 15.48 | 50863 |
| FS21 | FS12 | *Felis margarita* | 171.49798 | 11.460161 | 14.97 | 57531 |
| FS21 | FS2 | *Felis margarita* | 171.42282 | 11.359815 | 15.09 | 53303 |
| FS21 | Dhzankent | *Felis margarita* | 160.43269 | 63.095555 | 2.543 | 1855 |
| FS21 | Levant lybica | *Felis margarita* | 150.35494 | 9.507685 | 15.81 | 77340 |
| FS21 | modern_European_catus | *Felis margarita* | 138.90525 | 8.728777 | 15.91 | 81089 |
| FS21 | Ancient_European/Anatolia_catus | *Felis margarita* | 124.68283 | 17.452172 | 7.144 | 8715 |
| FS21 | ornata | *Felis margarita* | 119.25503 | 7.460807 | 15.98 | 65445 |
| FS21 | silvestries | *Felis margarita* | 117.79287 | 7.376722 | 15.97 | 69232 |
| FS21 | bieti | *Felis margarita* | 112.53383 | 7.053864 | 15.95 | 59429 |
| FS21 | FS21 | *Felis margarita* | 0 | -1 | 0 | -1 |
| Dhzankent | FS1 | *Felis margarita* | 150.71702 | 55.144777 | 2.733 | 1878 |
| Dhzankent | FS2 | *Felis margarita* | 165.10907 | 58.683773 | 2.814 | 1966 |
| Dhzankent | FS12 | *Felis margarita* | 180.25164 | 77.922663 | 2.313 | 2125 |
| Dhzankent | FS21 | *Felis margarita* | 160.43269 | 63.095555 | 2.543 | 1855 |
| Dhzankent | Dhzankent | *Felis margarita* | 0 | -1 | 0 | -1 |
| Dhzankent | Ancient_European/Anatolia_catus | *Felis margarita* | 116.97977 | 23.262014 | 5.029 | 5356 |
| Dhzankent | Modern_China/East_Asia_catus | *Felis margarita* | 120.93274 | 23.63015 | 5.118 | 7215 |
| Dhzankent | modern_European_catus | *Felis margarita* | 109.55539 | 21.377054 | 5.125 | 8111 |
| Dhzankent | Levant lybica | *Felis margarita* | 113.87666 | 22.249337 | 5.118 | 6969 |
| Dhzankent | silvestries | *Felis margarita* | 97.587149 | 19.052361 | 5.122 | 6541 |
| Dhzankent | ornata | *Felis margarita* | 96.563797 | 18.809416 | 5.134 | 5567 |
| Dhzankent | bieti | *Felis margarita* | 91.958451 | 17.888265 | 5.141 | 4915 |
| Modern China/E Asia_cat | FS1 | *Felis margarita* | 180.21786 | 11.322142 | 15.92 | 81007 |
| Modern China/E Asia_cat | FS2 | *Felis margarita* | 174.72111 | 11.255582 | 15.52 | 84297 |
| Modern China/E Asia_cat | FS12 | *Felis margarita* | 172.25718 | 11.165351 | 15.43 | 91443 |
| Modern China/E Asia_cat | FS21 | *Felis margarita* | 179.48462 | 11.363918 | 15.79 | 80396 |
| Modern China/E Asia_cat | Dhzankent | *Felis margarita* | 120.93274 | 23.63015 | 5.118 | 7215 |
| Modern China/E Asia_cat | Ancient_European/Anatolia_catus | *Felis margarita* | 85.16037 | 6.992342 | 12.18 | 30964 |
| Modern China/E Asia_cat | Modern_China/East_Asia_catus | *Felis margarita* | 0 | -1 | 0 | -1 |
| Modern China/E Asia_cat | modern_European_catus | *Felis margarita* | 98.470755 | 2.751153 | 35.79 | 350175 |
| Modern China/E Asia_cat | Levant lybica | *Felis margarita* | 99.70676 | 2.790807 | 35.73 | 342163 |
| Modern China/E Asia_cat | silvestries | *Felis margarita* | 91.460678 | 2.55638 | 35.78 | 350244 |
| Modern China/E Asia_cat | ornata | *Felis margarita* | 89.930103 | 2.514533 | 35.76 | 338484 |
| Modern China/E Asia_cat | bieti | *Felis margarita* | 87.520902 | 2.442639 | 35.83 | 334647 |

**Table S6.**
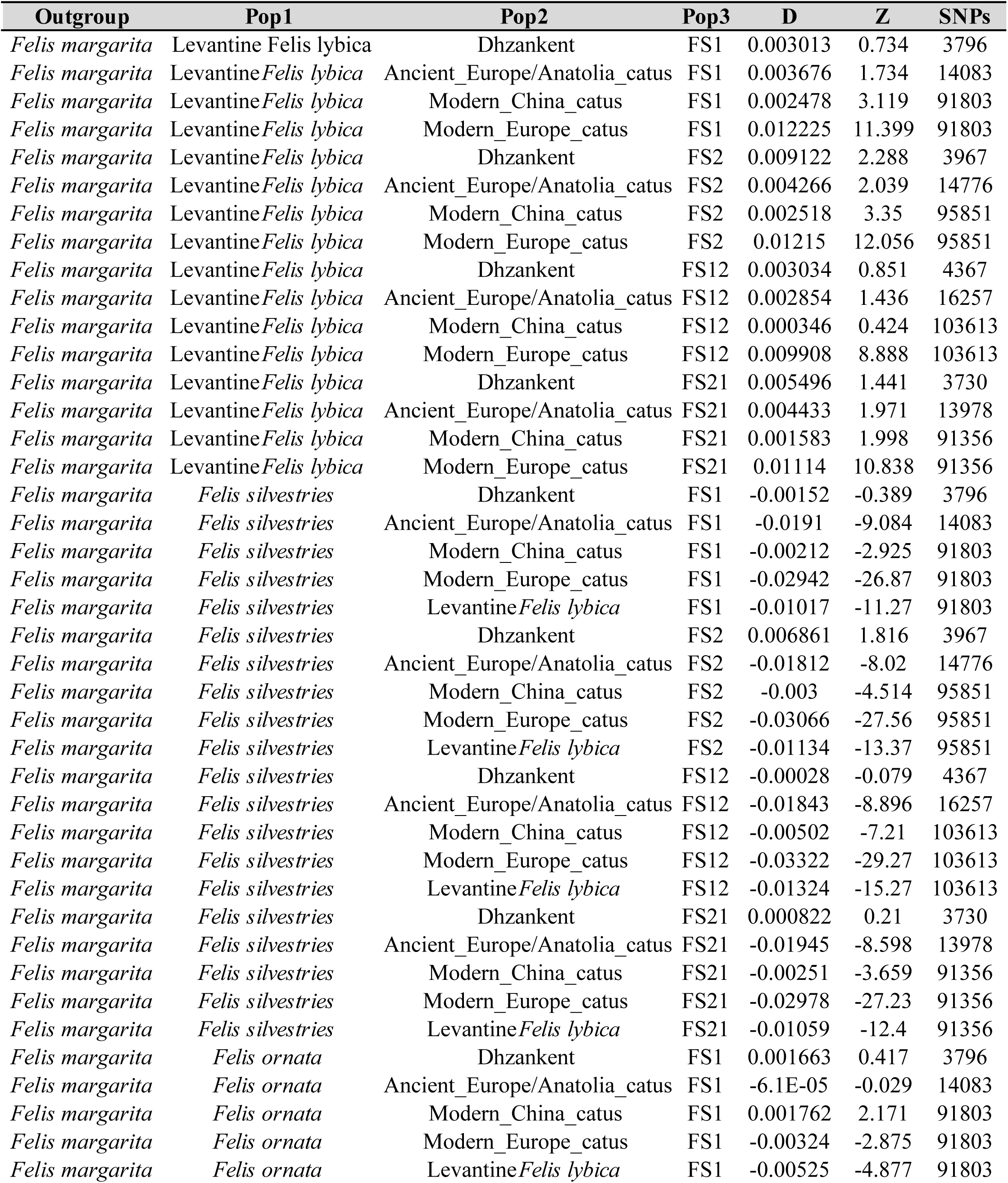
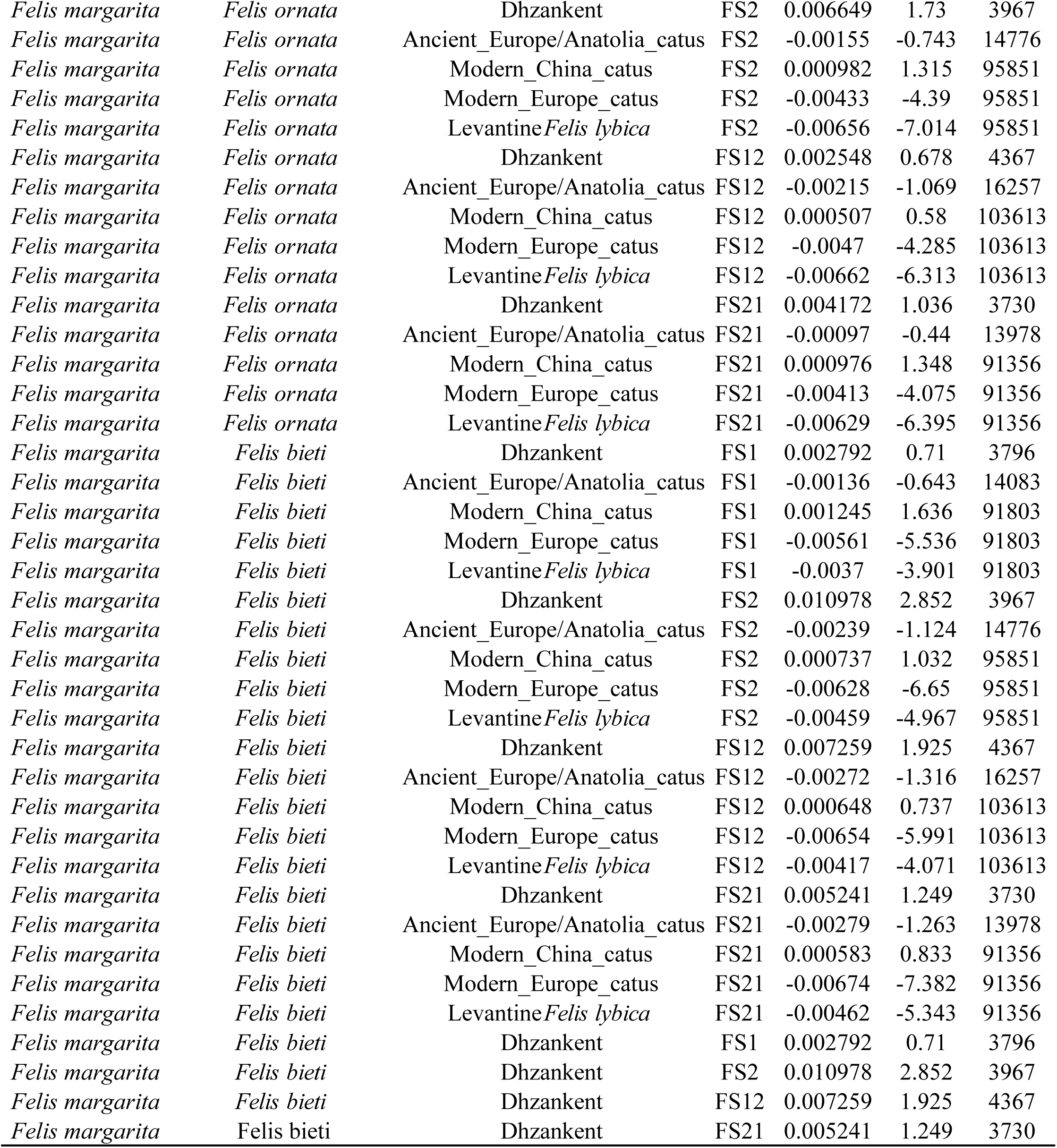
D-statistics test for gene flow between ancient Chinese domestic cats (FS1, FS2, FS12, FS21) and wildcats (F. lybica, F. silvestris, F. bieti, F. ornata) with the analysis set up as D(Outgroup, wildcatsX, Pop2, ancient_China) (Data of Fig. 3D).

**Table S7.** (A) Genotypes and inferred phenotypes of FS12 associated with coat colors.

| Coat color | Gene | Ref genome | Chr | Position | mutation | If FS12 carries |  |
| --- | --- | --- | --- | --- | --- | --- | --- |
|  |  |  |  |  |  | the allele | Reference |
| Black | *ASIP* | catus_9.0 | A3 | 25086566-25086567 | Deletion | No | (Eizirik et al., 2003) |
| Amber | *MC1R* | catus_9.0 | E2 | 63829934 | G>A | No | (Peterschmitt et al., 2009) |
| Russet | *MC1R* | catus_9.0 | E2 | 63830124-63830126 | Deletion | No | (Gustafson et al., 2017) |
| Copal | *MC1R* | catus_9.0 | E2 | 63830324-63830353 | Deletion | No | (Abitbol & Gache, 2019) |
| Dilution | *MLPH* | catus_9.0 | C1 | 219396820 | Deletion | NA | (Ishida et al., 2006) |
| Siamese coat colour pattern | *TYR* | catus_9.0 | D1 | 45898803 | C>T | No | (Lyons, Imes, et al., 2005) |
| Burmese coat colour pattern | *TYR* | catus_9.0 | D1 | 45907839 | C>A | No | (Lyons, Imes, et al., 2005) |
| Complete albinism | *TYR* | catus_9.0 | D1 | 45898772 | Deletion | No | (Imes et al., 2006) |
| Albinism | *TYR* | catus_9.0 | D1 | 45812806 | G>A | No | (Abitbol et al., 2017) |
| Mocha | *TYR* | catus_9.0 | D1 | 45898609-45898771 | Duplication | NA | (Y. Yu et al., 2019) |
| Cinnamon | *TYRP1* | catus_9.0 | D4 | 40069161 | C>T | No | (Lyons, Foe, et al., 2005) |
| Chocolate | *TYRP1* | catus_9.0 | D4 | 40081229 | G>A | No | (Lyons, Foe, et al., 2005) |
| White spotting | *KIT* | catus_9.0 | B1 | 164038110-164038111 | Insertion | Yes | (David et al., 2014) |

**Table S7.** (B) Genotypes and inferred phenotypes of FS12 associated with coat patterns.

| Pigmentation pattern | Gene | Ref genome | Chr | Position | mutation | If FS12 carries |  |
| --- | --- | --- | --- | --- | --- | --- | --- |
|  |  |  |  |  |  | the allele | Reference |
| Ticked | *DKK4* | catus_9.0 | B1 | 42620835 | C>T | No | (Lyons et al., 2021) |
| Ticked | *DKK4* | catus_9.0 | B1 | 42621481 | G>A | No | (Lyons et al., 2021) |
| Blotched tabby | *LVRN* | catus_9.0 | A1 | 96753652 | C>A | No | (Kaelin et al., 2012) |
| Blotched tabby | *LVRN* | catus_9.0 | A1 | 96754158 | G>A | No | (Kaelin et al., 2012) |
| Blotched tabby | *LVRN* | catus_9.0 | A1 | 96813484 | G>A | No | (Kaelin et al., 2012) |

**Table S7.** (C) Genotypes and inferred phenotypes of FS12 associated hair morphology.

| Hair type | Gene | Ref genome | Chr | Position | mutation | If FS12 carries the allele | Reference |
| --- | --- | --- | --- | --- | --- | --- | --- |
| Long hair | *FGF5* | catus_9.0 | B1 | 142165952 | T>G | No | (Drögemüller et al., 2007) |
| Long hair | *FGF5* | catus_9.0 | B1 | 142165953 | Deletion | No | (Drögemüller et al., 2007) |
| Long hair | *FGF5* | catus_9.0 | B1 | 142177919 | G>A | No | (Kehler et al., 2007) |
| Long hair | *FGF5* | catus_9.0 | B1 | 142185646-142185647 | Insertion A | No | (Kehler et al., 2007) |
| Long hair | *FGF5* | catus_9.0 | B1 | 142165850 | C>T | No | (Shaffer et al., 2021) |
| Lykoi | *HR* | catus_9.0 | B1 | 36040784-36040785 | dup | NA | (Buckley, Gandolfi, et al., 2020) |
| Lykoi | *HR* | catus_9.0 | B1 | 36040933 | Indel CAG | No | (Buckley, Gandolfi, et al., 2020) |
| Lykoi | *HR* | catus_9.0 | B1 | 36045776 | G>A | No | (Buckley, Gandolfi, et al., 2020) |
| Lykoi | *HR* | catus_9.0 | B1 | 36047047 | C>T | No | (Buckley, Gandolfi, et al., 2020) |
| Lykoi | *HR* | catus_9.0 | B1 | 36047518 | C>T | No | (Buckley, Gandolfi, et al., 2020) |
| Lykoi | *HR* | catus_9.0 | B1 | 36051555-36051556 | Insertion GACA | NA | (Buckley, Gandolfi, et al., 2020) |
| Sphynx hairless | *KRT71* | catus_9.0 | B4 | 81048680 | C>T | No | (Gandolfi et al., 2010) |
| Curly | *KRT71* | catus_9.0 | B4 | 81046358-81046359 | Insertion A | No | (Gandolfi et al., 2010) |
| Curly | *LPAR6* | catus_9.0 | A1 | 22919307_22919310 | Deletion | No | (Gandolfi, Alhaddad, Affolter, et al., 2013) |
| Curly | *LIPH* | catus_9.0 | C2 | 84136341_84136347 | Indel | No | (Manakhov et al., 2020) |
| Curly | *KRT71* | catus_9.0 | B4 | 81050264 | C>G | No | (Gandolfi, Alhaddad, Joslin, et al., 2013) |

**Table S7.** (D) Genotypes and inferred phenotypes of FS12 associated with tail length and morphology.

| Tail length | Gene | Ref genome | Chr | Position | mutation | If FS12 carries |  |
| --- | --- | --- | --- | --- | --- | --- | --- |
|  |  |  |  |  |  | the allele | Reference |
| Short tail | *HES7* | catus_9.0 | E1 | 2918735 | A>G | No | (Xu et al., 2016) |
| Short tail | *T* | catus_9.0 | B2 | 152019395 | Deletion | No | (Buckingham et al., 2013) |
| Short tail | *T* | catus_9.0 | B2 | 152019424 | Deletion | No | (Buckingham et al., 2013) |
| Short tail | *T* | catus_9.0 | B2 | 152021360-152021362 | Deletion | No | (Buckingham et al., 2013) |
| Short tail | *T* | catus_9.0 | B2 | 152021363-152021379 | Duplication | N/A | (Buckingham et al., 2013) |

**Table S7.** (E) Genotypes and inferred phenotypes of FS12 associated with genetic diseases.

| Disease | Gene | Chr | Ref genome | Position | Mutation | If FS12 carries the genotype | Reference |
| --- | --- | --- | --- | --- | --- | --- | --- |
| Factor XII Deficiency (Variant 1) | *F12* | A1 | catus_9.0 | 175381113-175381114 | Deletion | No | (Bender et al., 2015) |
| Factor XII Deficiency (Variant 2) | *F12* | A1 | catus_9.0 | 175382065 | Deletion | No | (Maruyama et al., 2017) |
| Hypertrophic Cardiomyopathy | *MYBPC* | D1 | catus_9.0 | 101329646 | C>T | No | (Meurs et al., 2007) |
| Hypertrophic Cardiomyopathy | *MYBPC* | D1 | catus_9.0 | 1340772 | C>G | No | (Meurs et al., 2005) |
| Polycystic Kidney Disease (PKD) | *PKD1* | E3 | catus_9.0 | 42858112 | C>A | No | (Lyons et al., 2004) |
| Polycystic Kidney Disease (PKD) | *PKD1* | E3 | catus_9.0 | 42848725 | Deletion | No | (Shitamori et al., 2023) |
| Polycystic Kidney Disease (PKD) | *PKD1* | E3 | catus_9.0 | 42850283 | C>T | No | (Shitamori et al., 2023) |
| Mucopolysaccharidosis Type VI Modifier | *ARSB* | A1 | catus_9.0 | 145138738 | G>A | No | (Crawley et al., 1998) |
| Mucopolysaccharidosis Type VI Modifier | *ARSB* | A1 | catus_9.0 | 145138869 | A>G | No | (Yogalingam et al., 1996) |
| Acute Intermittent Porphyria (Variant 1) | *HMBS* | D1 | catus_9.0 | 16540928-16540931 | Deletion | No | (Clavero et al., 2013) |
| Acute Intermittent Porphyria (Variant 2) | *HMBS* | D1 | catus_9.0 | 16544575 | G>A | No | (Clavero et al., 2013) |
| Acute Intermittent Porphyria (Variant 3) | *HMBS* | D1 | catus_9.0 | 16541469 | Insertion | No | (Clavero et al., 2013) |
| Acute Intermittent Porphyria (Variant 4) | *HMBS* | D1 | catus_9.0 | 16544592-16544594 | Deletion | No | (Clavero et al., 2013) |
| Acute Intermittent Porphyria (Variant 5) | *HMBS* | D1 | catus_9.0 | 16541613 | G>A | No | (Clavero et al., 2013) |
| Autoimmune Lymphoproliferative Syndrome | *FASL* | F1 | catus_9.0 | 16871911 | Insertion | No | (Aberdein et al., 2017) |
| Burmese Head Defect | *ALXI* | B4 | catus_9.0 | 110088245-110088254 | Deletion | No | (Lyons, Erdman, et al., 2016) |
| Chediak-Higashi Syndrome | *LYST* | D2 | catus_9.0 | 13289499-13308862 | Insertion | N/A | (Buckley, Grah, et al., 2020) |
| Congenital Adrenal Hyperplasia | *CYP11B1* | F2 | catus_9.0 | 84247412 | G>A | No | (Owens et al., 2012) |
| Congenital Erythropoietic Porphyria | *UROS* | D2 | catus_9.0 | 83467845 | C>T | No | (Clavero et al., 2010) |
| Congenital Myasthenic Syndrome | *COLQ* | C2 | catus_9.0 | 135068287 | G>A | No | (Abitbol, Hitte, et al., 2015) |
| Cystinuria Type 1A | *SCL3A1* | A3 | catus_9.0 | 66539609 | C>T | No | (Mizukami et al., 2015) |
| Cystinuria Type B (Variant 1) | *SCL7A9* | E2 | catus_9.0 | 20228722 | C>T | No | (Mizukami et al., 2016) |
| Cystinuria Type B (Variant 2) | *SCL7A9* | E2 | catus_9.0 | 20218196 | G>A | No | (Mizukami et al., 2016) |
| Cystinuria Type B (Variant 3) | *SCL7A9* | E2 | catus_9.0 | 20219555 | T>A | No | (Mizukami et al., 2016) |
| Dihydropyrimidinase Deficiency | *DPYS* | F2 | catus_9.0 | 52064442 | G>A | No | (Chang et al., 2012) |
| Earfold and Osteochondrodysplasia | *TRPV4* | D3 | catus_9.0 | 18884250 | G>T | No | (Gandolfi et al., 2016) |
| Familial Episodic Hypokalemic Polymyopathy | *WNK4* | E1 | catus_9.0 | 43304422 | C>T | No | (Gandolfi et al., 2012) |
| Glutaric Aciduria Type II | *ETFDH* | B1 | catus_9.0 | 71374289 | T>G | No | (Wakitani et al., 2014) |
| Glycogen Storage Disease | *GBE1* | C2 | catus_9.0 | 34744477-34781895 | Insertion | N/A | (Fyfe et al., 2007) |
| GM1 Gangliosidosis | *GLB1* | C2 | catus_9.0 | 158932167 | C>G | No | (D. R. Martin et al., 2008) |
| GM2 Gangliosidosis | *GM2A* | A1 | catus_9.0 | 198114664-198114667 | Deletion | No | (D. R. Martin et al., 2005) |
| GM2 Gangliosidosis Type II | *HEXB* | A1 | catus_9.0 | 141571788-141571812 | Inversion | N/A | (D. R. Martin et al., 2004) |
| GM2 Gangliosidosis Type II | *HEXB* | A1 | catus_9.0 | 141565348 | C>T | No | (Kanae et al., 2007) |
| Hemophilia B (Variant 1) | *F9* | X | catus_9.0 | 117111577 | C>T | No | (Goree et al., 2005) |
| Hemophilia B (Variant 2) | *F9* | X | catus_9.0 | 117091961 | G>A | No | (Goree et al., 2005) |
| Hypertrophic Cardiomyopathy | *MYBPC* | D1 | catus_9.0 | 101340772 | G>C | No | (Meurs et al., 2005) |
| Hypertrophic Cardiomyopathy | *MYBPC* | D1 | catus_9.0 | 101329646 | G>A | No | (Meurs et al., 2007) |
| Hypotrichosis | *FONXN1* | E1 | catus_9.0 | 18255880-18255883 | Deletion | No | (Abitbol, Bossé, et al., 2015) |
| Lipoprotein Lipase Deficiency | *LPL* | B1 | catus_9.0 | 38078551 | G>A | No | (Ginzinger et al., 1996) |
| MDR1 Medication Sensitivity | *ABCB1* | A2 | catus_9.0 | 93144356-93144357 | Deletion | No | (Mealey & Burke, 2015) |
| Mucopolysaccharidosis Type I | *IDUA* | B1 | catus_9.0 | 207800585- 207800587 | Deletion | No | (X. He et al., 1999) |
| Mucopolysaccharidosis Type VI | *ARSB* | A1 | catus_9.0 | 145138869 | A>G | No | (Yogalingam et al., 1996) |
| Mucopolysaccharidosis Type VI | *ARSB* | A1 | catus_9.0 | 145138738 | C>T | N/A | (Crawley et al., 1998) |
| Mucopolysaccharidosis Type VII (Variant 1) | *GUSB* | E3 | catus_9.0 | 16120173 | G>A | No | (Fyfe et al., 1999) |
| Mucopolysaccharidosis Type VII (Variant 2) | *GUSB* | E3 | catus_9.0 | 16123229/16123232 | T>G/C>T | No | (Wang et al., 2015) |
| Myotonia Congenita | *CLCN1* | A2 | catus_9.0 | 158986498 | G>T | No | (Gandolfi et al., 2014) |
| Progressive Retinal Atrophy | *CEP290* | B4 | catus_9.0 | 112522818 | A>C | No | (Menotti-Raymond et al., 2007) |
| Progressive Retinal Atrophy | *KIF3B* | A3 | catus_9.0 | 26784019 | G>A | No | (Cogné et al., 2020) |
| Progressive Retinal Atrophy | *AIPL1* | E1 | catus_9.0 | 940445 | C>T | No | (Lyons, Creighton, et al., 2016) |
| Pyruvate Kinase Deficiency | *PKLR* | F1 | catus_9.0 | 70310110 | G>A | No | (Grah, Grah, et al., 2012) |
| Sphingomyelinosis (Variant 1) | *NPC1* | D3 | catus_9.0 | 48234217 | C>G | No | (Somers et al., 2003) |
| Sphingomyelinosis (Variant 1) | *NPC1* | D3 | catus_9.0 | 48250290 | T>G | N/A | (Mauler et al., 2017) |
| Sphingomyelinosis (Variant 2) | *NPC2* | B3 | catus_9.0 | 121872988 | C>T | No | (Zampieri et al., 2014) |
| Spinal Muscular Atrophy | *LIX1* | A1 | catus_9.0 | 161036890-161176706 | Deletion | No | (Fyfe et al., 2006) |
| Vitamin D-Dependent Rickets | *CYP27B1* | B4 | catus_9.0 | 86180281 | Deletion | N/A | (Geisen et al., 2009) |
| Vitamin D-Dependent Rickets | *CYP27B1* | B4 | catus_9.0 | 86180375 | C>A | No | (Grah, Ellis, et al., 2012) |
| Polydactyly | *LMBR1* | A2 | catus_9.0 | 169532842 | T>A | No | (Lettice et al., 2008) |
| Polydactyly | *LMBR1* | A2 | catus_9.0 | 169532844 | T>C | No | (Lettice et al., 2008) |
| Polydactyly | *SHH* | A2 | catus_9.0 | 169533066 | C>G | No | (Lettice et al., 2008) |

**Table S8.** Information of ancient artwork related to cats in China between 827 CE and 1930 CE within the time frame of this research.

| Theme | Date | Period | Type | Black | White | Unclear | Number of individuals | Collector/Source |
| --- | --- | --- | --- | --- | --- | --- | --- | --- |
| *M126 of the Tombs of the Tang and Song Dynasties at Liujiazhuang Locus North in Anyang City, Henan* | 827 CE | Tang | Tomb mural | 1 | 1 | 2 | 3 | The excavation report of the Tombs of the Tang and Song Dynasties at Liujiazhuang Locus North in Anyang City, Henan |
| *Tang Dynasty Mural Tomb at Beiguan, Anyang City, Henan* | 835-907 CE | Tang | Tomb mural | 1 | 1 | 0 | 1 | Preliminary Report on the Excavation of Tang Dynasty Mural Tombs at Beiguan, Anyang City, Henan |
| *Anyang Xiaonanhai Mural* | 960-1,127 CE | Song | Tomb mural | 1 | 0 | 0 | 1 | Zhaoling's Painted Brick Chamber from the Tang Dynasty |
| *Dengfeng Heishangou Song Tomb* | 11th - 12th centuries CE | Song | Tomb mural | 1 | 1 | 0 | 1 | Research on Images from Song Dynasty Tomb in Dengfeng's Heishan Gou by Yi Qing |
| *Xu Congbin's Tomb* | 982 CE | Song | Tomb mural | 1 | 1 | 0 | 1 | Newly Added Fort's Liao Dynasty Tomb of Xu Congyuan |
| *Chen Shu's Cat and butterfly Painting* | 17th - 18th centuries CE | Qing | Legacy painting | 1 | 2 | 0 | 2 | The Palace Museum in Beijing |
| *Cheng Zhang's Two Cats Peering at Fish* | 1930 CE | modern times | Legacy painting | 2 | 2 | 0 | 2 | The Palace Museum in Beijing |
| *Qiu Ying's Portrait of a Lady* | 16th century CE | Ming | Legacy painting | 2 | 3 | 0 | 3 | The Palace Museum in Beijing |
| *Li Di's Winter Play with Infant* | 10th - 11th centuries CE | Song | Legacy painting | 1 | 1 | 0 | 1 | National Palace Museum in Taipei |
| *Wealthy FloralCat Painting* | 12th - 13th centuries CE | Song | Legacy painting | 1 | 1 | 0 | 1 | National Palace Museum in Taipei |
| *Siblings in Unity Painting* | 13th - 14th centuries CE | Yuan | Legacy painting | 1 | 1 | 0 | 1 | National Palace Museum in Taipei |
| *Yi Yuanji's Cat and Monkey* | 11th century CE | Song | Legacy painting | 2 | 2 | 0 | 2 | National Palace Museum in Taipei |
| *Wealth in the Pot Painting* | 15th century CE | Ming | Legacy painting | 1 | 1 | 0 | 1 | National Palace Museum in Taipei |
| *Cat Under the Banana Shade* | 14th-17th centuries CE | Min | Legacy painting | 4 | 4 | 0 | 4 | The Metropolitan Museum of Art |
| *Xuanzong's Scroll of FiveCats* | 15th century CE | Min | Legacy painting | 4 | 5 | 0 | 5 | The Metropolitan Museum of Art |
| *Tao Cheng'sCat in Fragrant Grass* | 15th century CE | Min | Legacy painting | 6 | 5 | 0 | 14 | National Palace Museum in Taipei |
| *Li Di'sCat and Dragonfly* | 10th - 11th centuries CE | Song | Legacy painting | 1 | 1 | 0 | 1 | Collection of Osaka City Museum of Fine Arts, Japan |
| *Cat Painting* | 10th - 11th centuries CE | Song | Legacy painting | 1 | 1 | 0 | 1 | National Palace Museum in Taipei |
| *LittleCat Shadow* | 10th - 11th centuries CE | Song | Legacy painting | 0 | 1 | 0 | 1 | National Palace Museum in Taipei |
| *Cat and Infant Play Painting* | 10th - 13th centuries CE | Song | Legacy painting | 2 | 2 | 0 | 2 | Museum of Fine Arts, Boston |
| *Song Huizong's Cat and butterfly Painting (Signature)* | 12th centuries CE | Song | Legacy painting | 1 | 1 | 0 | 1 | National Palace Museum in Taipei |
| *Li Di's Okra Mountain and Rock* | 13th century CE | Southern Song | Legacy painting | 1 | 1 | 0 | 1 | National Palace Museum in Taipei |
| *Shen Zhenlin's Cat and butterfly in Spring Album, WisteriaCat* | 19th century CE | Qing | Legacy painting | 10 | 15 | 0 | 15 | National Palace Museum in Taipei |
| *Shang Xi's Playful Cat Scroll* | 15th century CE | Ming | Legacy painting | 1 | 2 | 0 | 2 | National Palace Museum in Taipei |
| *Five Cats Painting* | 12th - 13th centuries CE | Song | Legacy painting | 4 | 4 | 0 | 5 | Private Collection |
| *Shen Zhenlin's Cat and Bamboo* | 19th century CE | Qing | Legacy painting | 1 | 3 | 0 | 3 | National Palace Museum in Taipei |
| *Zhou Wenju's Portrait of a Lady* | 10th century CE | Five Dynasties | Legacy painting | 1 | 1 | 0 | 1 | National Palace Museum in Taipei |
| *Liangkai's Scroll of Leisurely Delights of the cats* | 12th century CE | Song | Legacy painting | 2 | 5 | 0 | 5 | Nara Yamato Bunkakan in Japan |
| *Luo Zonggui's Hibiscus Cat Play Fan Painting* | 10th - 13th centuries CE | Song | Legacy painting | 1 | 2 | 0 | 2 | N/A |
| *Mao Yi's Sichuan Hibiscus and Wandering Cat* | 13th century CE | Southern Song | Legacy painting | 3 | 3 | 0 | 4 | Nara Yamato Bunkakan in Japan |
| *Jin Qing's Twin Cats* | 10th - 13th centuries CE | Song | Legacy painting | 1 | 2 | 0 | 2 | N/A |
| *Wang Weilie's Cat and Rock* | 17th century CE | Qing | Legacy painting | 1 | 1 | 0 | 1 | National Art Museum of China |
| *Wang Guxiang's Cat and butterfly Painting* | 16th century CE | Ming | Legacy painting | 1 | 1 | 0 | 1 | N/A |
|  |  |  |  | 62 | 77 | 2 | 91 |  |
|  |  |  |  | 68% | 85% |  |  |  |

**Notes:** In a painting where an individual has both black and white fur, the color statistics would be recorded once under both the black ("Black" in table title) and white ("White" in table title) categories. The final proportion of colors is calculated as the count of records for black or white divided by the total number of observed individuals ("Number of individuals" in table title).

